# Ultra-low-dose same-target combinations control drug resistance across stages

**DOI:** 10.64898/2026.11.08.764983

**Authors:** Zhi-Gao Zhao, Shao-Qing Cai, Rui Tan, Weijing Cai, Qian Zhao, Xuan Wang, Feng Xu

## Abstract

The emergence of drug resistance continues to limit the long-term effectiveness of cancer therapies [1], as tumour cells adapt to therapeutic selective pressure through diverse genetic and non-genetic routes [2,3]. Despite advances in more potent and next-generation inhibitors and mechanism-based combination therapies [4,5], durable disease control remains difficult to achieve. Here we tested an alternative strategy: combining multiple inhibitors directed against the same oncogenic driver, each at ultra-low doses with minimal individual activity, to delay resistance emergence. In epidermal growth factor receptor (EGFR)-driven cancer models, such ultra-low-dose same-target combinations (ULDCs) distributed individually weak perturbations across multiple inhibitors directed against the same target and achieved substantial growth inhibition in vitro and antitumour activity in vivo despite weak effects of the corresponding single agents. Under similar levels of growth inhibition (∼50%), single-agent EGFR inhibitors generated resistant outgrowth, whereas a five-drug ULDC delayed resistance emergence. Increasing the number of inhibitors further delayed resistance emergence, with no detectable resistant outgrowth observed for combinations containing 8–14 inhibitors within the experimental timeframe. The five-drug ULDC also retained measurable growth-inhibitory activity against gefitinib-resistant PC-9/GR cells, and resistant cells arising under five-drug ULDC treatment exhibited markedly greater reversibility following drug withdrawal than those arising under single-agent treatment. Although pharmacological additivity or synergy may contribute to the collective growth-inhibitory effects of ULDCs, the divergent long-term resistance outcomes observed under similar levels of growth inhibition indicate that resistance behaviour is not determined by inhibitory strength alone. Instead, our findings support a model in which increasing same-target inhibitor diversity progressively constrains the adaptive trajectories available during resistance evolution, while therapeutic perturbation architecture may also influence the stability of resistant states that emerge.

More broadly, these findings support the concept of same-target collective suppression, in which multiple individually weak perturbations directed against the same target can collectively produce substantial antitumour activity and constrain resistance evolution across multiple stages. We propose that increasing same-target inhibitor diversity may restrict accessible adaptive trajectories, while therapeutic perturbation architecture may also influence the stability of resistant states.

## INTRODUCTION

Drug resistance remains a central challenge in cancer therapy [1,6,7]. Targeted therapies directed against oncogenic drivers often produce substantial initial responses, yet these responses are rarely durable [4,8,9]. In EGFR-mutant non-small-cell lung cancer (NSCLC), advances in the development of EGFR tyrosine kinase inhibitors (TKIs) have significantly improved clinical outcomes [10,11,12], but resistance frequently emerges through diverse genetic and molecular alterations [2,13,14], as well as non-genetic mechanisms involving epigenetic plasticity and adaptive cellular states [2,3]. Several strategies have been developed to delay acquired resistance, including next-generation inhibitors with improved activity against resistant alterations and mechanism-based combination therapies targeting distinct or acquired resistance mechanisms [4,5,15].

Although these approaches can improve disease control, resistance remains common, reflecting the extensive molecular and phenotypic adaptability of tumour cells under therapeutic selective pressure [2,3,7,13,14].

A defining feature of resistance is that tumour cells can access multiple potential routes of adaptation under therapeutic pressure. These routes include genetic or phenotypic changes that reduce drug sensitivity while preserving sufficient cellular function to sustain growth [2,3,13]. The set of such possibilities can be conceptualized as an evolutionary resistance landscape, in which the availability of adaptive routes influences how readily resistance emerges [16,17]. Current therapeutic strategies may reduce tumour burden, but do not necessarily eliminate the availability of these adaptive routes.

We therefore asked whether the availability of such routes could be influenced by how therapeutic perturbations are structured, rather than solely by the magnitude of growth inhibition. Specifically, we tested whether increasing the diversity of inhibitors directed against the same oncogenic target could alter resistance emergence under approximately matched levels of growth inhibition. In this study, same-target inhibitor diversity is operationally defined by the number of distinct EGFR-TKIs included in a combination. Building on our previous observations that ultra-low-dose combinations of same-target inhibitors can produce collective antitumour effects [18], we defined ULDCs as combinations of multiple inhibitors directed against the same oncogenic driver at concentrations where individual inhibitors produce only minimal growth inhibition when administered alone, typically within the IC_05_–IC_20_ range. Here, “ultra-low-dose” refers to these minimally active concentrations, with IC_05_, IC_10_, IC_15_, and IC_20_ denoting the concentrations of each individual inhibitor corresponding to 5%, 10%, 15%, and 20% growth inhibition, respectively, as determined in the respective cell model. Using EGFR-driven cancer models, we show that such combinations produce substantial same-target collective growth-inhibitory activity in vitro and antitumour activity in vivo despite minimal activity of the individual agents, progressively delay detectable resistance emergence as inhibitor diversity increases, and, in the five-drug setting, retain activity against established resistant cells and are associated with more reversible resistant states (Fig. 1 and Supplementary Fig. 1).

**Fig. 1.**
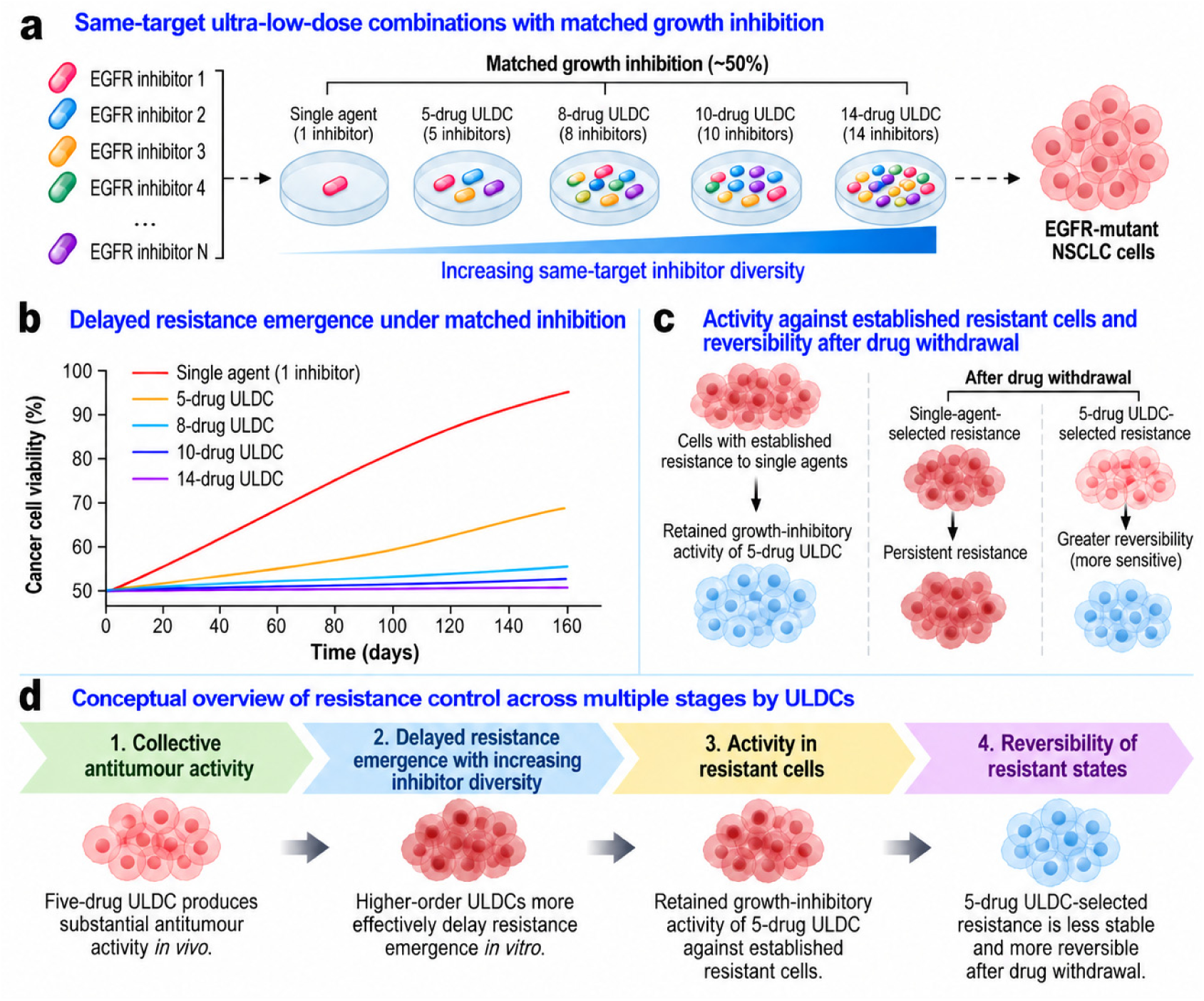
Study overview of how ultra-low-dose same-target combinations control cancer cell drug resistance across multiple stages. **a**, Schematic illustration of the ultra-low-dose same-target combination (ULDC) strategy and resistance-evolution experimental design. Single EGFR-TKIs and their multi-drug same-target combinations were administered under approximately matched levels of growth inhibition (∼50%) during repeated cyclic drug exposure. Individual inhibitors within combinations were used at concentrations producing minimal growth inhibition when administered alone, enabling assessment of how increasing same-target inhibitor diversity influences resistance evolution under comparable levels of inhibitory strength. **b**, Schematic representation of resistance emergence with increasing same-target inhibitor diversity under approximately matched levels of growth inhibition (∼50%). Single-agent treatment is associated with resistance emergence, whereas increasing inhibitor diversity is associated with progressively delayed detectable resistance emergence, with higher-order combinations (≥8 inhibitors) showing no detectable resistant outgrowth within the experimental timeframe. **c**, Schematic overview of resistance after formation, illustrating retained growth-inhibitory activity of the five-drug ULDC against established resistant cells and greater reversibility of five-drug-ULDC-selected resistance following treatment withdrawal compared with single-agent-selected resistance. **d**, Conceptual overview of resistance control across multiple stages by ULDCs. ULDCs produce collective antitumour activity before resistance emergence and delay resistance emergence during evolutionary adaptation. The five-drug ULDC retains activity against established resistant states, and resistant states arising under five-drug ULDC pressure are less stable and more reversible, suggesting that ULDCs influence multiple stages of resistance evolution.

Together, these findings support the concept of same-target collective suppression, in which multiple individually weak perturbations directed against the same target can act collectively to reshape both tumour inhibition and resistance evolution. Beyond direct tumour inhibition, this framework suggests that therapeutic perturbation architecture may also shape resistance evolution by constraining accessible adaptive trajectories and influencing the stability of resistant states that emerge.

## RESULTS

### Ultra-low-dose same-target combinations collectively inhibit cancer cell growth despite minimal single-agent activity

To test whether multiple inhibitors, each administered at an ultra-low dose and thereby providing an individually weak perturbation directed against the same target, can generate collective growth inhibition, we treated EGFR-mutant NSCLC cells with five-drug combinations comprising the EGFR-TKIs gefitinib, erlotinib, afatinib, neratinib, and osimertinib, each at concentrations corresponding to IC_05_–IC_20_ (Supplementary Table 2). Across multiple lung cancer cell models, these combinations produced clear antiproliferative activity despite minimal effects of the corresponding single agents, establishing the pharmacological basis for the growth-inhibitory component of what we term “same-target collective suppression”.

In lung cancer PC-9 cells, individual EGFR-TKIs showed minimal inhibition across the tested ultra-low-dose range, whereas the five-drug ULDC generated substantially greater growth inhibition compared with individual inhibitors at the same IC_05_–IC_20_ conditions (Fig. 2a and Supplementary Tables 3 and 4). The five-drug ULDC displayed a dose-dependent growth inhibition, reaching approximately 50% inhibition when each component was combined at its respective IC_15_ concentration. Combination index (CI) analysis [19,20] yielded CI values below 1 across multiple dose levels, consistent with CI-defined synergistic pharmacological interactions among the combined inhibitors under these experimental conditions (Supplementary Table 5). Removal of any single component slightly reduced overall combination activity, indicating that the observed inhibition reflected distributed weak contributions from multiple components rather than dependence on a dominant inhibitor (Supplementary Fig. 3 and Supplementary Table 7).

**Fig. 2.**
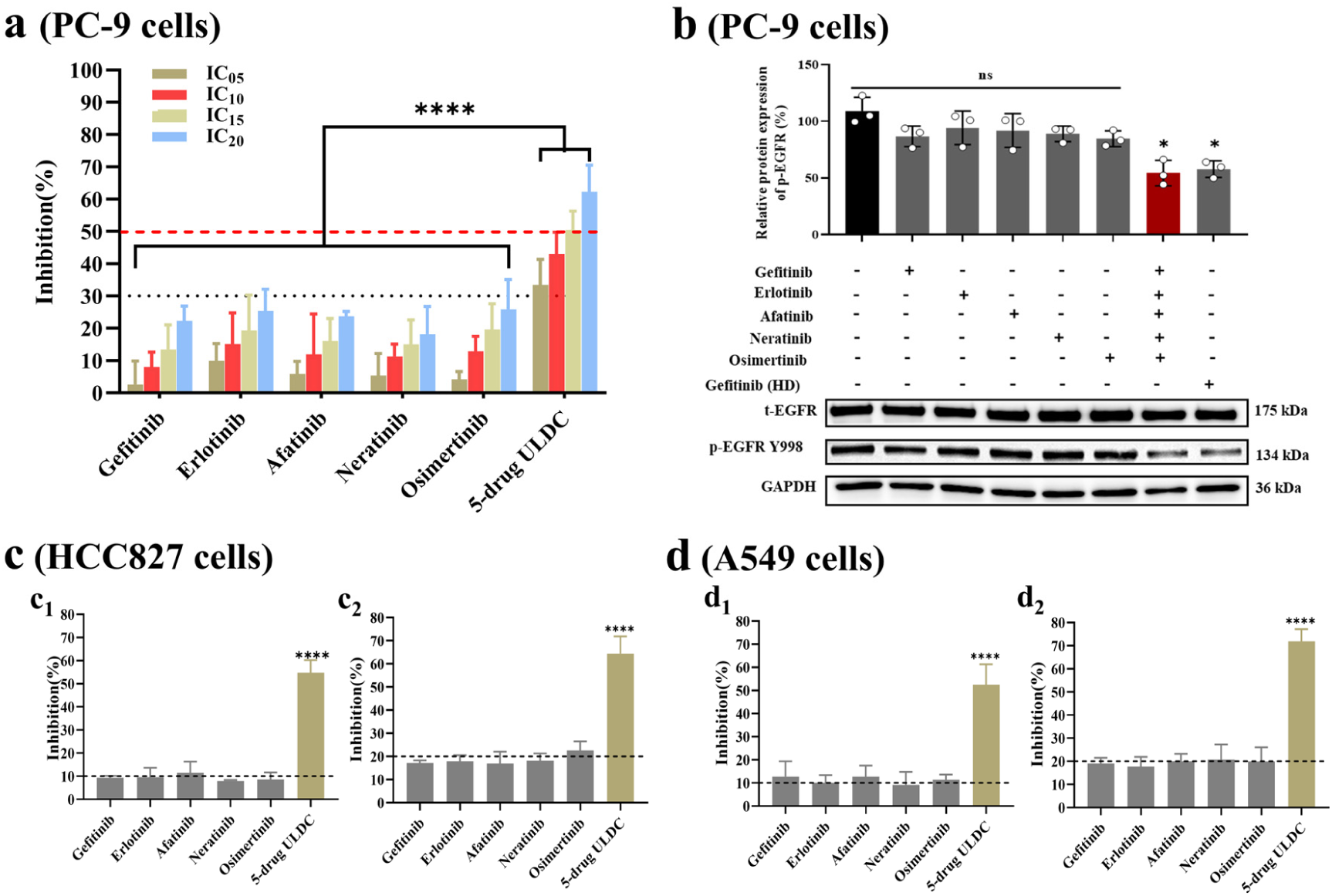
Ultra-low-dose same-target combinations (five-drug ULDCs) generate collective growth-inhibitory activity across lung cancer cell models. **a**, Collective inhibitory effects of the five-drug ULDC in EGFR-mutant PC-9 cells. Individual EGFR-TKIs were administered alone at ultra-low concentrations (IC_05_–IC_20_), and the five-drug ULDC was generated by combining the corresponding IC_05_–IC_20_ concentrations of each individual inhibitor. The five-drug ULDC achieved substantial collective growth-inhibitory activity despite the minimal inhibitory activity of each individual inhibitor. **b**, Inhibition of EGFR phosphorylation by the five-drug ULDC in PC-9 cells. Individual inhibitors were tested alone at IC_15_ concentrations, whereas the five-drug ULDC was generated by combining the corresponding IC_15_ concentrations of each inhibitor. Total EGFR (t-EGFR) remained unchanged, whereas phosphorylation of EGFR (p-EGFR) was markedly reduced following ULDC treatment. Gefitinib (HD), high-dose gefitinib control (IC_50_); GAPDH, loading control. **c**, Validation of collective growth-inhibitory activity in an additional EGFR-mutant lung cancer cell model (HCC827). Individual inhibitors and the five-drug ULDC were evaluated at two dose levels based on the respective IC_10_ and IC_20_ values determined in HCC827 cells. **d**, Validation in KRAS-mutant, EGFR-wild-type A549 cells. The same evaluation was performed at IC_10_ and IC_20_ dose levels determined in A549 cells. Collective growth-inhibitory activity was also observed in this model, although less pronounced than in EGFR-mutant models. The horizontal dotted and dashed lines indicate the corresponding growth inhibition levels on the y-axis. Statistical comparisons were performed relative to the corresponding single-drug groups. ****P<0.0001; *P<0.05; ns, not significant. Data are presented as mean ± SD (n=3).

To examine whether collective growth inhibition was associated with inhibition of the shared molecular target, we assessed EGFR phosphorylation in PC-9 cells. Individual EGFR-TKIs administered at IC_15_ concentrations produced minimal effects on p-EGFR levels, whereas the five-drug ULDC markedly reduced p-EGFR levels (Fig. 2b). Notably, p-EGFR inhibition by the ULDC was comparable to that achieved by the high-dose gefitinib control (IC_50_), whereas total EGFR (t-EGFR) protein abundance remained largely unchanged, suggesting that the ULDC primarily affected EGFR phosphorylation without substantially altering total EGFR abundance (Fig. 2b).

We next evaluated whether this collective growth inhibition could be observed across lung cancer models with distinct genetic backgrounds and degrees of EGFR dependency (Supplementary Note 1.1.5). In EGFR-mutant HCC827 cells, five-drug ULDCs produced substantial growth inhibition under corresponding ultra-low-dose conditions (Fig. 2c). Similar, although less pronounced, growth-inhibitory effects were also observed in KRAS-mutant, EGFR-wild-type A549 cells, which are markedly less sensitive to EGFR-TKIs than EGFR-driven models (Fig. 2d). Together, these findings show that multiple individually weak EGFR-targeted perturbations can generate collective growth-inhibitory activity across genetically distinct lung cancer cell models with varying degrees of EGFR dependency.

### The delayed emergence of drug resistance cannot be explained by growth-inhibitory strength alone

We next examined whether ULDCs alter the emergence of drug resistance under sustained treatment pressure. Under repeated cyclic drug exposure (Supplementary Fig. 13), all five single-agent EGFR-TKIs reproducibly gave rise to resistant outgrowth, with resistance emerging between 20 and 60 days depending on inhibitor identity (Fig. 3a and Supplementary Fig. 15).

**Fig. 3.**
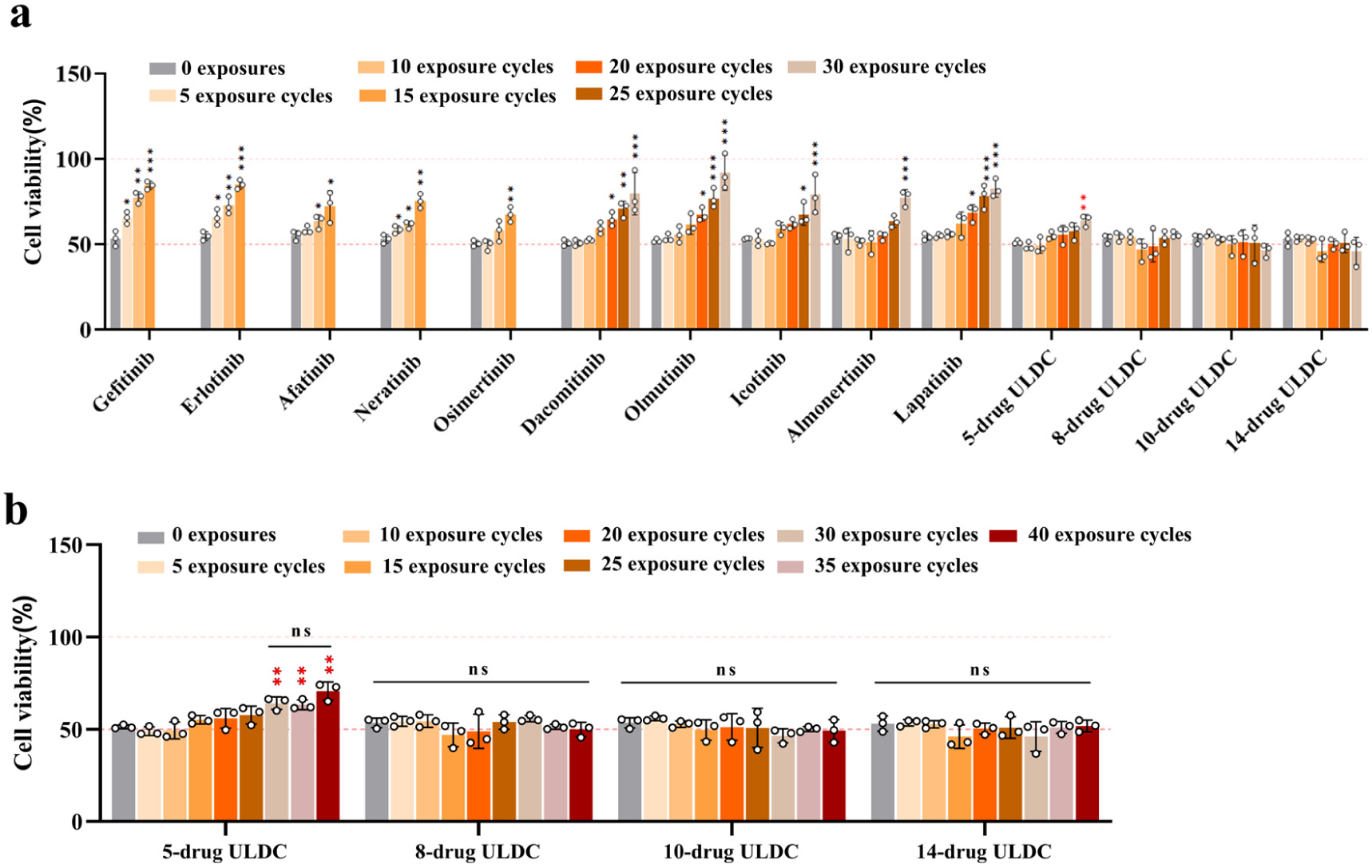
Increasing same-target inhibitor diversity progressively delays resistance emergence in ULDCs. **a**, Resistance emergence during repeated cyclic drug exposure of PC-9 cells treated with individual EGFR-TKIs or same-target ULDCs containing increasing inhibitor diversity under approximately matched levels of growth inhibition (∼50%). Cell viability was measured after 5–30 exposure cycles (20–120 days), as indicated. Each exposure cycle consisted of 2 days of drug treatment followed by 2 days of recovery in drug-free complete medium. **b**, Increasing same-target inhibitor diversity progressively delays resistance emergence. Whereas the five-drug ULDC eventually exhibited detectable resistant outgrowth, ULDCs containing 8, 10, or 14 inhibitors maintained growth inhibition without detectable resistant outgrowth throughout the observation period (40 exposure cycles; 160 days). The horizontal dotted and dashed lines indicate the corresponding cell viability levels on the y-axis. Statistical comparisons were performed within each treatment group, using parental PC-9 cells (0 exposure cycles) as the reference. ***P < 0.001, **P < 0.01, and *P < 0.05; ns, not significant. Data are presented as mean ± SD (n=3).

By contrast, the five-drug ULDC maintained drug sensitivity for substantially longer, with resistant outgrowth becoming evident only after prolonged treatment and reaching a clear resistant phenotype by around day 120 (Fig. 3 and Supplementary Fig. 15). Importantly, all treatments were administered at concentrations producing approximately matched levels of growth inhibition (∼50%). Therefore, the delayed resistance observed under five-drug ULDC treatment cannot be attributed solely to greater inhibitory strength.

These results indicate that resistance behaviour is not determined by inhibitory strength alone but is also influenced by how the therapeutic perturbation is structured within a same-target combination.

### Increasing same-target inhibitor diversity progressively delays resistance emergence

Under fixed per-drug ultra-low-dose conditions, increasing same-target inhibitor diversity progressively enhanced growth inhibition, accompanied by a proportional increase in total drug exposure with inhibitor number (Fig. 2, Supplementary Figs. 10, 11, and 12). Because increased growth inhibition and total drug exposure could potentially contribute to delayed resistance, subsequent resistance experiments were performed under approximately matched levels of growth inhibition (∼50%), allowing resistance outcomes to be compared while minimizing differences in growth-inhibitory strength. Under these conditions, resistance emergence displayed a non-linear relationship with same-target inhibitor diversity (as described below), indicating that resistance outcomes could not be explained by inhibitory strength alone.

To determine how same-target inhibitor diversity influences long-term resistance behaviour, we systematically increased the number of EGFR inhibitors included in ULDCs while maintaining approximately matched growth-inhibitory conditions. The eight-drug ULDC comprised the original five inhibitors plus dacomitinib, olmutinib, and icotinib; the ten-drug ULDC additionally included almonertinib and lapatinib; and the fourteen-drug ULDC further incorporated furmonertinib, rezivertinib, befotertinib, and oritinib. Increasing same-target inhibitor diversity was associated with progressively delayed resistant outgrowth. Although several newly introduced EGFR-TKIs showed slower resistance emergence as single agents than the first five inhibitors tested, all ten single agents tested ultimately generated resistance within 120 days of repeated drug exposure (Fig. 3a). By contrast, no detectable resistant outgrowth was observed in the eight-, ten-, and fourteen-drug ULDC groups throughout the 160-day observation period of repeated cyclic drug exposure (Fig. 3b).

The relationship between same-target inhibitor diversity and resistance behaviour was strongly non-linear, progressing from resistance emergence with all ten single agents tested, to delayed resistance with the five-drug ULDC, and to no detectable resistant outgrowth within the experimental timeframe in higher-order same-target ULDCs (Fig. 3). These findings indicate that the differences in resistance emergence associated with increasing same-target inhibitor diversity cannot be explained by growth-inhibitory strength alone.

### Five-drug ULDC retains activity after resistance formation, and ULDC-selected resistance is more reversible

We next asked whether the five-drug ULDC retains growth-inhibitory activity after resistance has emerged. In gefitinib-resistant PC-9/GR cells, the five-drug ULDC retained measurable growth-inhibitory activity, whereas most individual EGFR-TKIs showed only limited activity under the corresponding ultra-low-dose conditions (Fig. 4a and Supplementary Note 2.2). This relative advantage of the combination over its individual components was observed across multiple dose levels. To assess the stability of acquired resistance, resistant cell populations generated under repeated treatment in this study were serially passaged in drug-free medium. The resistance phenotype that emerged under five-drug ULDC treatment declined rapidly after treatment withdrawal, with sensitivity largely restored toward parental PC-9 cell levels after three recovery periods. By contrast, resistance arising under single-agent EGFR-TKIs remained substantially more persistent, with drug sensitivity failing to return to parental PC-9 levels even after five recovery periods (Fig. 4b; Supplementary Note 2.3).

**Fig. 4.**
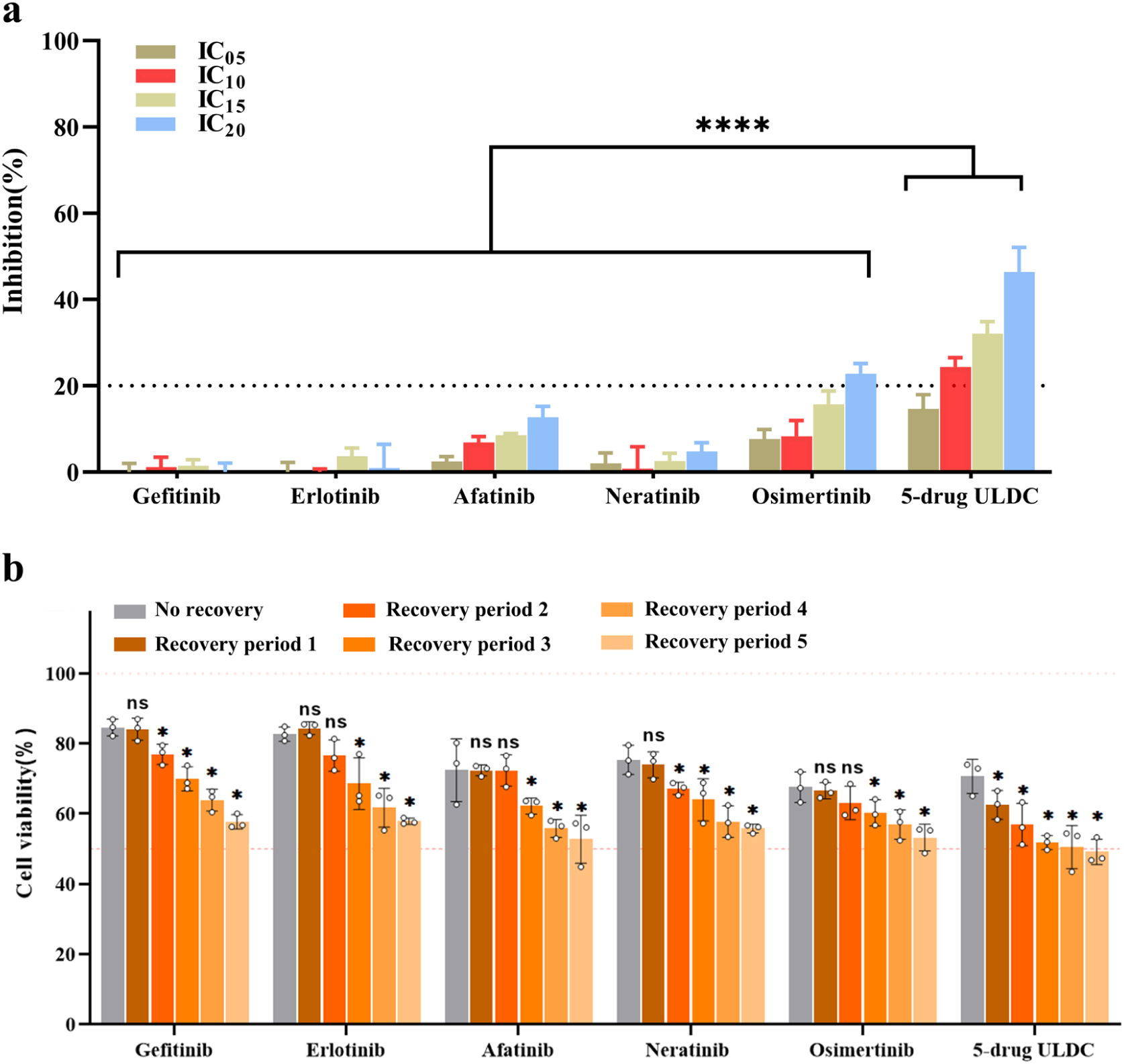
Five-drug ULDC retains activity against resistant cells and is associated with more reversible resistant states. **a**, Collective growth-inhibitory activity of the five-drug ULDC in established gefitinib-resistant PC-9/GR cells. EGFR-TKIs were tested individually at ultra-low concentrations (IC_05_–IC_20_ determined in PC-9 cells), whereas the five-drug ULDC was generated by combining the corresponding concentrations of the five inhibitors. The five-drug ULDC produced substantially greater growth inhibition than the corresponding individual inhibitors under the same ultra-low-dose conditions. Compared with each corresponding single-agent treatment group, ****P < 0.0001. Mean ± SD (n=3). **b**, Reversibility of resistant states after drug withdrawal. Single-agent-selected resistant populations were obtained after 15 exposure cycles with the corresponding individual inhibitors, whereas ULDC-selected resistant populations were obtained after 40 exposure cycles with the five-drug ULDC. Resistant populations were subsequently cultured in drug-free complete medium, and drug sensitivity was reassessed using the same treatment conditions employed during the original repeated cyclic exposure experiments, corresponding to approximately matched 50% growth inhibition in parental PC-9 cells. Compared with single-agent-selected resistant populations, five-drug-ULDC-selected resistant populations exhibited greater reversibility, with drug sensitivity returning toward parental PC-9 cell levels. Compared with the corresponding no-recovery group, *P < 0.05 indicates a statistically significant difference, whereas ns indicates no statistically significant difference (P ≥ 0.05). Mean ± SD (n=3). The horizontal dotted and dashed lines indicate the corresponding growth inhibition or cell viability levels on the y-axis.

These findings indicate that five-drug ULDC treatment is associated not only with delayed resistance emergence but also with the emergence of a more reversible resistant state. This suggests that the architecture of therapeutic perturbation may influence not only the timing of resistance emergence but also the stability and reversibility of the resistant state that arises.

### The five-drug ULDC induces distinct transcriptomic responses

To explore the molecular responses associated with same-target ULDC treatment, we performed transcriptomic profiling of PC-9 cells treated with the five-drug ULDC or individual EGFR-TKIs under approximately matched levels of growth inhibition (∼50%). Compared with individual inhibitors, the ULDC induced a broader transcriptional response, including a substantial set of ULDC-associated differentially expressed genes (Supplementary Figs. 19 and 20). Pathway enrichment analysis further identified biological processes related to DNA replication, genome maintenance, and cancer-associated signalling (Supplementary Figs. 18–22). In addition, differential expression of genes previously associated with drug resistance, including CCND1 [21], HMGB1 [22,23], and CFL1 [24], was observed, further supporting the presence of a distinct transcriptional response to the same-target ULDC. Together, these findings show that five-drug ULDC treatment is associated with a distinct and broader transcriptional response than individual EGFR-TKI treatment under approximately matched levels of growth inhibition (Supplementary Note 3).

### The five-drug ULDC inhibits tumour growth in vivo

To assess in vivo relevance, we evaluated the five-drug EGFR-TKI ULDC in a PC-9 xenograft mouse model. The five-drug ULDC markedly inhibited tumour growth despite the limited activity of the corresponding individual ultra-low-dose EGFR-TKIs and approached the efficacy of the high-dose gefitinib control group, demonstrating substantial collective antitumour activity in vivo (Figs. 5a and 5b).

**Fig. 5.**
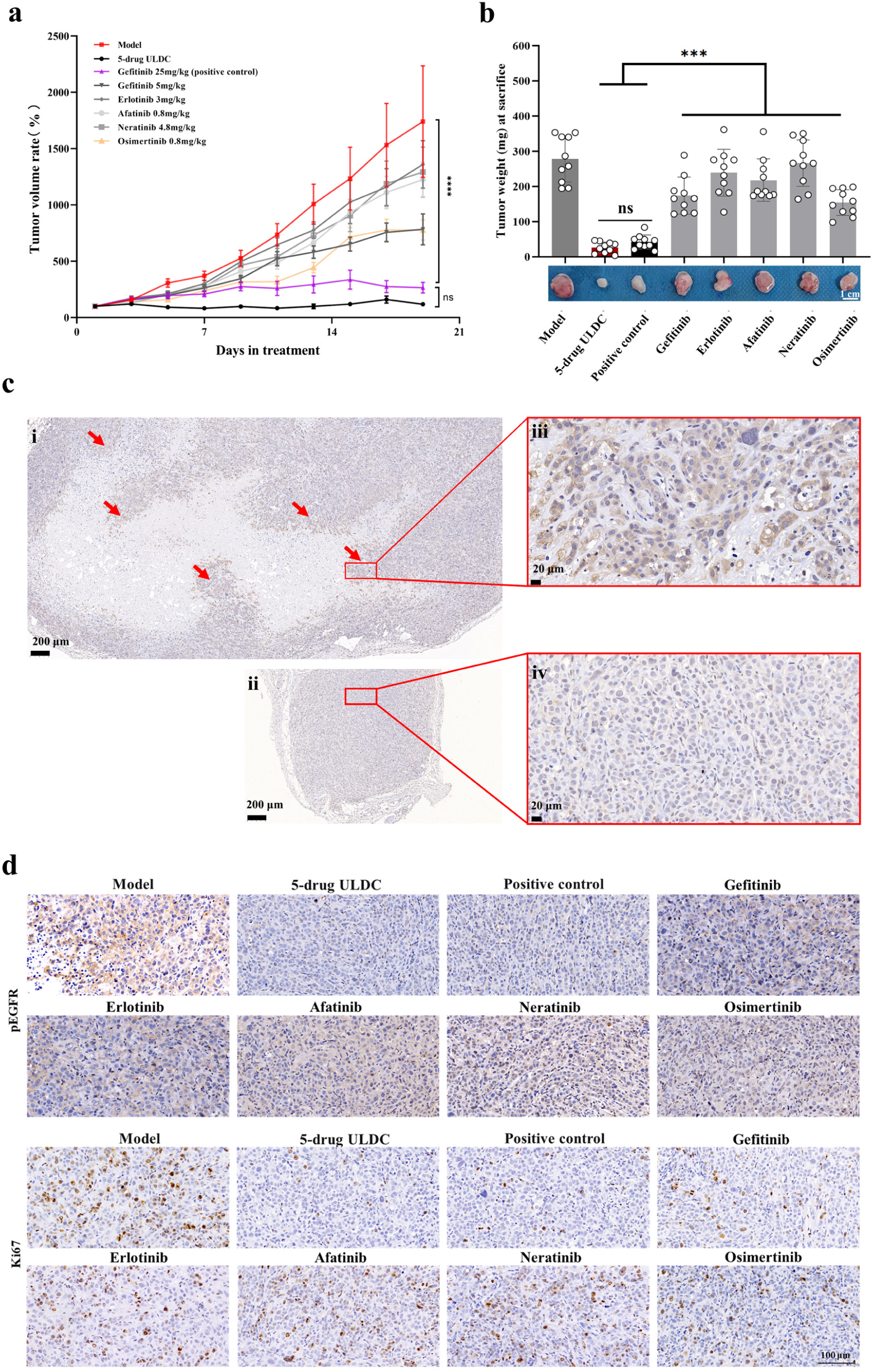
The five-drug ULDC inhibits tumour growth in vivo. PC-9 xenograft-bearing nude mice were treated by oral gavage for 3 weeks (5 consecutive treatment days followed by 2 days off per week). Each group comprised ten mice (five females and five males; n = 10). Individual EGFR-TKIs were administered at ultra-low doses that, after conversion to human-equivalent doses using standard Km factors, corresponded to approximately 5%–10% of their respective WHO-defined daily doses (DDDs). The five-drug ULDC was generated by combining these individually ultra-low doses, whereas gefitinib administered at 25 mg kg⁻¹ day⁻¹ using the same dosing schedule served as a high-dose positive control. **a**, Tumour growth curves of PC-9 xenografts during treatment. The five-drug ULDC markedly inhibited tumour growth compared with the model group and approached the antitumour activity observed with the high-dose gefitinib control group. Tumour volume ratio was calculated relative to tumour volume on the first treatment day. **b**, Final tumour weights at the study endpoint. The five-drug ULDC significantly reduced tumour burden compared with each corresponding single-agent treatment and showed no significant difference from the high-dose gefitinib control group. Individual tumours are shown below the graph. **c**, Representative p-EGFR immunohistochemical staining of tumour sections from model and five-drug ULDC-treated mice. Representative low- and high-magnification images are shown. Red arrows indicate the transition regions between viable tumour tissue and necrotic areas. i and iii, tumour tissues from the model group; ii and iv, tumour tissues from the five-drug ULDC treatment group. **d**, Representative immunohistochemical staining of p-EGFR and Ki67 in tumour tissues from each treatment group. The five-drug ULDC was associated with reduced p-EGFR and Ki67 staining compared with individual ultra-low-dose EGFR-TKI treatments, consistent with reduced EGFR phosphorylation and tumour-cell proliferation. Statistical comparisons were performed as indicated. Data are presented as mean ± SD.

Tumours from mice receiving the five-drug ULDC also showed reduced p-EGFR and lower Ki67 staining compared with individual ultra-low-dose EGFR-TKI treatments, consistent with reduced target phosphorylation and tumour-cell proliferation in vivo (Figs. 5c and 5d). Treatment was well tolerated under the conditions tested, with no apparent treatment-associated body-weight loss or overt histopathological or biochemical abnormalities detected during the treatment period (Supplementary Figs. 23–29). These findings indicate that the five-drug ULDC achieves substantial antitumour activity in vivo with acceptable tolerability under the conditions tested.

Collectively, our findings show that same-target ULDCs can influence both antitumour activity and multiple stages of resistance evolution by maintaining substantial tumour inhibition, progressively delaying detectable resistance emergence as inhibitor diversity increases, and, in the five-drug setting, retaining activity against established resistant cells and being associated with more reversible resistant states.

## DISCUSSION

Drug resistance remains a major obstacle to durable cancer therapy, yet the factors governing its emergence remain incompletely understood. A central finding of this study is that resistance emergence is influenced not only by growth-inhibitory strength but also by same-target inhibitor diversity within the ultra-low-dose same-target combination (ULDC) strategy. While single-agent treatment reproducibly generated resistance under repeated exposure, the five-drug ULDC substantially delayed this process, and resistance emergence was progressively delayed as same-target inhibitor diversity increased, with no detectable resistant outgrowth in higher-order combinations within the experimental timeframe.

Notably, these divergent resistance outcomes were observed under approximately matched levels of growth inhibition (∼50%), indicating that they cannot be explained simply by differences in growth-inhibitory strength. Although total drug exposure and pharmacological interactions may contribute to the effects of higher-order combinations, the strongly non-linear relationship between same-target inhibitor diversity and long-term resistance behaviour suggests that these factors alone may not fully account for the observed pattern. Instead, our findings reveal a qualitative transition from resistance emergence with all ten single agents tested, to delayed resistance with the five-drug ULDC, and to no detectable resistant outgrowth in higher-order combinations within the experimental timeframe.

More broadly, the dissociation between approximately matched growth inhibition and divergent patterns of resistance emergence suggests that growth-inhibitory strength alone does not determine resistance behaviour. One potential contributor to this distinction is that inhibitors directed against the same target do not necessarily impose identical pharmacological perturbations. Although ULDCs combine inhibitors directed against the same oncogenic target, these inhibitors are not pharmacologically identical. Structural and mechanistic differences in how EGFR inhibitors engage the kinase can produce distinct inhibitor–target interactions and differential inhibitor sensitivity across EGFR mutational contexts [25,26]. Thus, the same target can be perturbed through diverse inhibitor-specific interactions, providing a pharmacological basis for the concept of same-target inhibitor diversity.

An important feature of this study is that all inhibitors were directed against the same molecular target. Although we did not directly compare same-target and cross-target strategies, the observed effects may, at least in part, depend on concentrating multiple weak perturbations from different inhibitors on the same target. One possible explanation is that concentrating distributed weak perturbations on the same target may restrict the range of accessible resistance routes. Such a model is compatible with the structural constraints governing EGFR kinase activation and regulation [27,28], as well as with evidence that EGFR mutational context can differentially influence inhibitor sensitivity [25,26,29,30].

By contrast, a previous study using low-dose combinations targeting multiple nodes within a signalling pathway reported loss of efficacy upon further dose reduction [31]. Together, these observations raise the hypothesis that the efficacy of combinations of individually weak perturbations may depend, at least in part, on whether those perturbations are dispersed across multiple targets or concentrated on the same target.

From a structural and functional perspective, resistance evolution under targeted therapy may be constrained by the functional space available to the target. Kinase domains such as EGFR can adopt multiple conformational states [27,28,32], and different kinase inhibitors can engage distinct conformational states or binding modes [26,32,33]. Moreover, individual resistance mutations can exhibit differential sensitivity to different inhibitors [25,29,30], while resistant populations can arise through distinct evolutionary trajectories and cellular states [34]. Together, these structural, pharmacological and evolutionary considerations suggest that resistance evolution may depend on a restricted set of viable adaptive solutions.

Consistent with this view, resistance emergence may depend not only on the existence of resistant variants but also on the availability of viable adaptive trajectories that maintain sufficient functional capacity to support tumour growth. Increasing same-target inhibitor diversity may progressively restrict accessible adaptive trajectories, thereby delaying resistance emergence and, under some conditions, resulting in no detectable resistant outgrowth within the experimental timeframe. Although we did not directly measure the underlying molecular processes, the strongly non-linear relationship between inhibitor diversity and resistance behaviour is consistent with progressive restriction of accessible adaptive trajectories. Together, these observations support a model in which increasing same-target inhibitor diversity within ULDCs may impose progressively greater adaptive constraint during resistance evolution (Fig. 6).

**Fig. 6.**
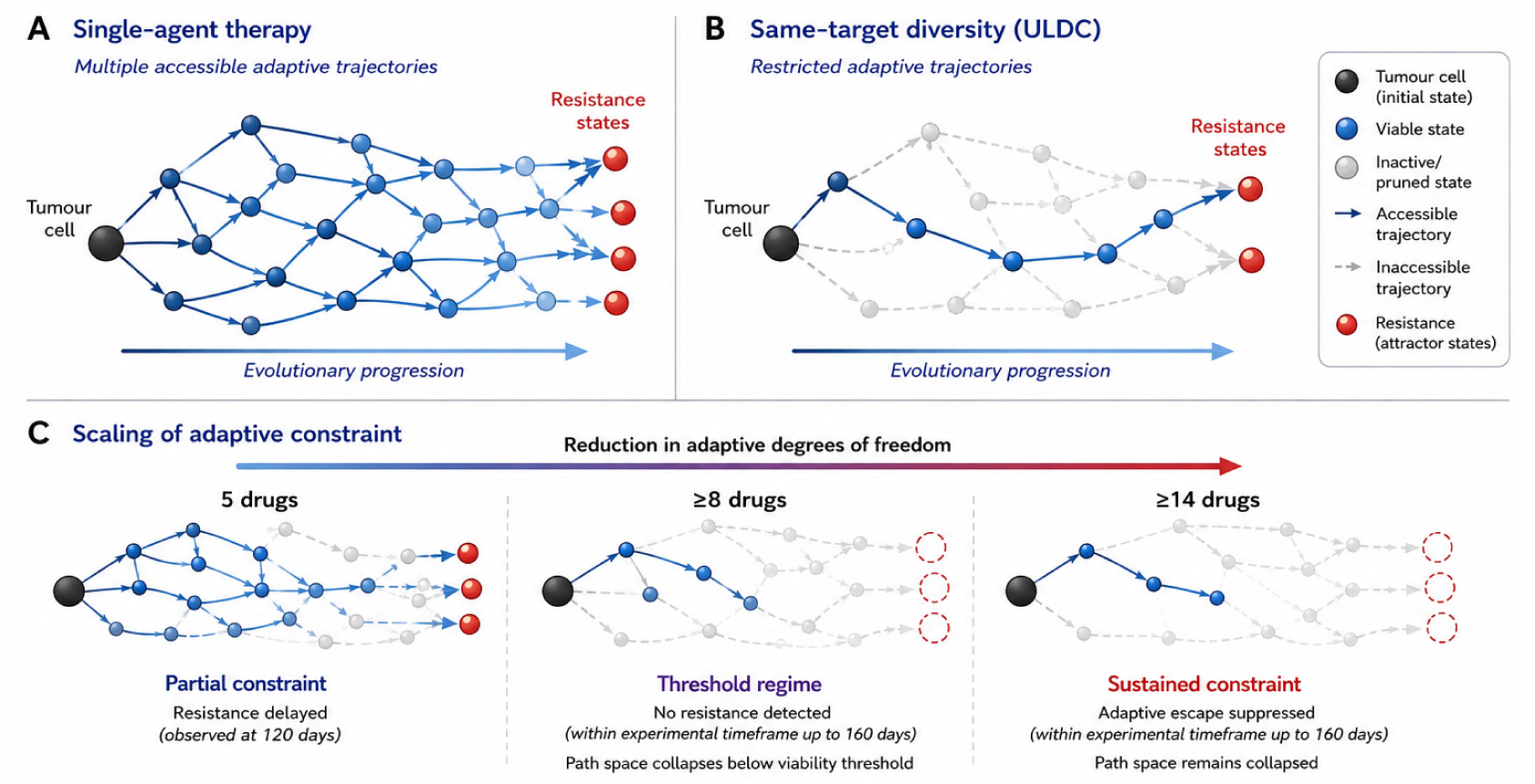
Conceptual model of progressive restriction of adaptive trajectories under ultra-low-dose same-target combinations. **A**, Adaptive landscape under single-agent therapy. Under single-agent treatment, tumour cells retain multiple accessible adaptive trajectories, allowing evolutionary escape toward resistant states through diverse evolutionary routes. **B**, Adaptive landscape under ultra-low-dose same-target combinations (ULDCs). Increasing same-target inhibitor diversity is proposed to progressively restrict accessible adaptive trajectories under approximately matched levels of growth inhibition, thereby limiting the evolutionary routes available for resistance development. **C**, Progressive restriction of adaptive trajectories with increasing same-target inhibitor diversity. Increasing same-target inhibitor diversity is proposed to progressively restrict adaptive degrees of freedom, thereby narrowing the range of accessible evolutionary trajectories. This progressive restriction provides the conceptual basis for the adaptive-constraint component of same-target collective suppression shown in Fig. 7.

More broadly, evolutionary perspectives on cancer have emphasized that adaptation under therapeutic selective pressure can involve genetic and epigenetic alterations as well as phenotypic plasticity [2,3,13,16,17]. Viewed in this context, these findings may reflect a broader restriction in the range of adaptive states accessible to tumour cells.

Within this framework, ULDCs represent a distinct architecture of therapeutic perturbation. Whereas conventional combination strategies distribute therapeutic selective pressure across multiple targets or pathways, the ULDC strategy described here distributes individually weak perturbations across multiple inhibitors directed against the same target. This same-target architecture may enable these distributed weak perturbations to collectively achieve substantial antitumour activity, while increasing inhibitor diversity within that architecture may progressively restrict the range of adaptive trajectories available to tumour cells under therapeutic selective pressure.

The observation of collective inhibitory effects in additional cellular models spanning different degrees of EGFR dependency suggests that collective growth-inhibitory activity of same-target ULDCs is not restricted to the highly EGFR-dependent PC-9 context. However, whether the broader resistance-constraining properties of ULDCs extend across different biological contexts remains to be established.

Consistent with this interpretation, transcriptomic profiling of the five-drug ULDC revealed a distinct transcriptional profile compared with individual inhibitors under approximately matched levels of growth inhibition, characterized by broader transcriptional changes, enrichment of differentially expressed genes in tumour-relevant pathways, and altered expression of genes previously associated with drug resistance. Further comparative analysis identified a substantial set of ULDC-associated transcriptional changes not shared with individual EGFR-TKI DEG sets. Although these observations do not directly resolve the mechanisms underlying the observed patterns of resistance emergence, they provide molecular context for the effects associated with the five-drug ULDC.

Beyond delaying resistance emergence, the five-drug ULDC retained inhibitory activity against an established gefitinib-resistant derivative (PC-9/GR), in which most individual EGFR-TKIs exhibited limited activity. Notably, resistance emerging under five-drug ULDC treatment exhibited substantially greater reversibility than resistance emerging under single-agent treatment. Following serial passages in drug-free medium, sensitivity to the five-drug ULDC was largely restored toward parental PC-9 cell levels, whereas resistance arising under individual EGFR-TKIs remained substantially more persistent. Together, these findings suggest that therapeutic perturbation architecture may influence not only the timing of resistance emergence but also the stability of the resulting resistant states. Tumour cells can reach EGFR inhibitor resistance through distinct evolutionary paths, with adaptive state transitions contributing to the emergence of different resistant states [3,34]. Our findings extend this evolutionary perspective by suggesting that the architecture of therapeutic perturbation may influence both the emergence of resistance and the stability of the resistant states that arise. Specifically, resistant cells arising under five-drug ULDC treatment exhibited greater reversibility following withdrawal of drug selection than those arising under prolonged single-agent treatment, suggesting that the therapeutic architecture under which resistance evolves may influence the stability of the resulting resistant state.

Tumour heterogeneity provides a substrate for adaptive escape during targeted therapy [2,7]. Against this evolutionary backdrop, our findings suggest that increasing same-target inhibitor diversity may impose an additional evolutionary constraint by limiting accessible adaptive trajectories. The distributed contribution pattern observed in drop-one analyses suggests that the inhibitory effects of the five-drug ULDC arise from distributed weak contributions across its components rather than dependence on a single dominant inhibitor. Collectively, these findings support what we term “same-target collective suppression”, in which multiple individually weak perturbations directed against the same target collectively produce substantial antitumour activity while also being associated with constrained resistance evolution (Fig. 7). Within this same-target architecture, increasing inhibitor diversity may progressively strengthen adaptive constraint by restricting the range of adaptive trajectories compatible with continued target function and tumour growth, thereby delaying detectable resistance emergence, with higher-order combinations showing no detectable resistant outgrowth within the experimental timeframe. Separately, the greater reversibility of resistance arising under five-drug ULDC treatment suggests that the architecture of therapeutic perturbation may also influence the stability of the resistant states that emerge. Rather than representing a dosing strategy alone, ULDCs therefore provide an experimental implementation of a broader conceptual framework for understanding how distributed weak perturbations acting on the same target can shape the evolutionary landscape of resistance under therapeutic pressure.

**Fig. 7.**
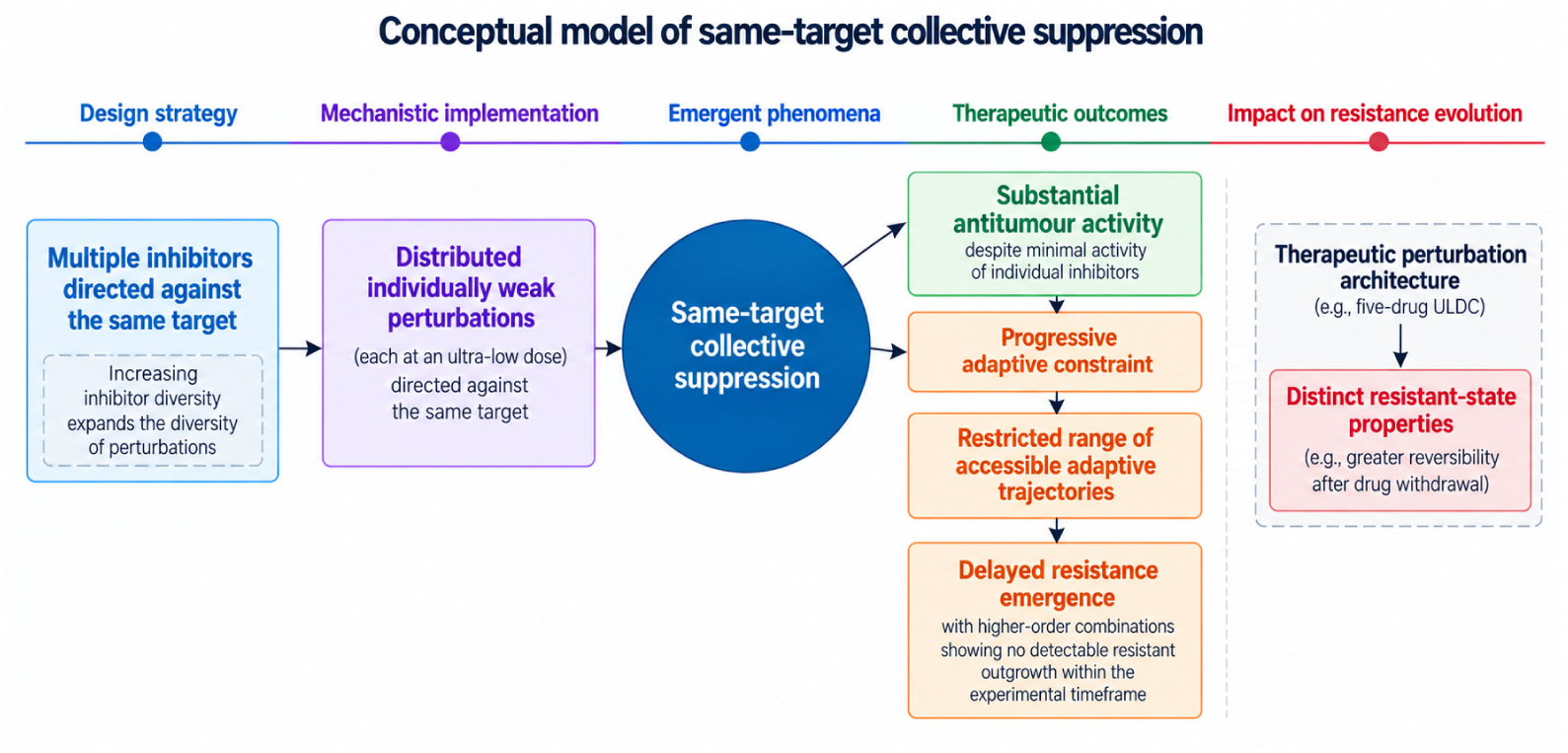
Conceptual model of same-target collective suppression. Administration of multiple inhibitors, each at an ultra-low dose, generates distributed individually weak perturbations directed against the same target. Increasing same-target inhibitor diversity expands the diversity of these perturbations within this architecture, allowing multiple individually weak perturbations to act collectively and give rise to same-target collective suppression. This concept encompasses substantial antitumour activity together with a proposed adaptive-constraint component of resistance evolution. Within this same-target architecture, increasing inhibitor diversity is proposed to progressively constrain resistance evolution by restricting the range of accessible adaptive trajectories, thereby delaying detectable resistance emergence; higher-order combinations showed no detectable resistant outgrowth within the experimental timeframe. Consistent with this broader framework, resistance arising under five-drug ULDC treatment exhibited greater reversibility following treatment withdrawal than resistance arising under single-agent treatment, suggesting that therapeutic perturbation architecture may additionally influence the stability of resistant states. This conceptual model integrates the findings of the present study into a unified framework in which same-target collective suppression connects therapeutic design with the evolutionary trajectory of resistance.

Existing approaches to limit resistance evolution have included identifying and targeting acquired resistance mechanisms [29,30]. Mechanism-based combination strategies similarly seek to suppress defined resistance pathways or adaptive vulnerabilities [5]. However, even three-drug cross-target combinations designed to inhibit multiple signalling nodes or adaptive feedback pathways can remain vulnerable to acquired resistance through heterogeneous genetic alterations that reactivate downstream signalling, reflecting the persistence of alternative adaptive trajectories under therapeutic pressure [35,36]. In contrast, ULDCs are not predicated on prior identification of specific resistance mechanisms, but instead rely on increasing same-target inhibitor diversity within a same-target architecture, with the potential to generate substantial collective antitumour activity while progressively constraining the adaptive trajectories available to tumour cells.

Beyond mechanism-based combination approaches, evolutionary treatment strategies have sought to influence resistance dynamics by modulating the selective conditions under which tumour populations evolve. Among these strategies, adaptive therapy seeks to modulate evolutionary dynamics by altering treatment intensity and scheduling over time [37,38]. ULDCs intervene along a different dimension by modifying the architecture and diversity of perturbations applied to a shared oncogenic target. Whereas adaptive therapy alters the temporal pattern of selection pressure, ULDCs distribute multiple individually weak perturbations across different inhibitors directed against the same target, thereby altering the structure and diversity of the perturbations imposed during treatment. By operating through a distinct dimension of evolutionary control, ULDCs may represent a complementary strategy for constraining resistance emergence while maintaining antitumour activity.

An important practical consideration for this strategy is tolerability. In vivo administration of five EGFR-TKIs at ultra-low doses was not associated with overt toxicity under the conditions tested, consistent with our previous observations with ultra-low-dose same-target combinations [18]. By distributing inhibitory pressure across multiple ultra-low-dose agents directed against the same target, this same-target architecture may limit the dose burden contributed by each individual component while retaining collective antitumour activity. Nevertheless, the safety, pharmacological properties, and clinical feasibility of higher-order same-target combinations will require further investigation.

Several limitations of this study should be acknowledged. First, we did not directly compare same-target and cross-target combinations under identical conditions, and thus the specific contribution of targeting architecture to resistance emergence remains to be determined. While cross-target combinations have been widely explored, the present study deliberately focuses on increasing inhibitor diversity within a same-target architecture to define its relationship with resistance emergence. Second, the molecular basis of resistance emergence, including the relative roles of genetic and non-genetic mechanisms, was not fully resolved. Third, while no resistant outgrowth was detected within the experimental timeframe in higher-order same-target combinations, longer-term studies or in vivo studies may reveal additional patterns of resistance emergence. Fourth, single-agent repeated-exposure resistance trajectories were established for the first ten EGFR-TKIs but not for the four additional inhibitors included only in the fourteen-drug ULDC. Accordingly, our data support conclusions regarding the impact of increasing same-target inhibitor diversity at the combination level, but not direct comparison between the fourteen-drug ULDC and each of its individual constituents.

In summary, our results reveal a previously unrecognised relationship between same-target inhibitor diversity within ultra-low-dose same-target combinations (ULDCs) and the emergence of resistance. These findings support a model in which limiting the evolution of resistance may depend not only on inhibitory potency but also on progressively constraining the adaptive trajectories available under therapeutic selection pressure. Taken together, these results support the concept of same-target collective suppression (Fig. 7), whereby individually weak perturbations distributed across multiple inhibitors and directed against the same target collectively maintain substantial antitumour activity and are proposed to impose adaptive constraint on the evolution of resistance. This framework extends current concepts of combination therapy design beyond maximizing inhibitory potency. Interpreted within an evolutionary framework [16,39], this study suggests that adaptive constraint may represent a broader design principle for combination therapies aimed at limiting the evolution of resistance.

## METHODS

### Section 1. Cell lines and cell culture

Human non-small-cell lung cancer (NSCLC) cell lines PC-9, HCC827, and A549, together with the gefitinib-resistant PC-9 derivative PC-9/GR, were obtained from Shanghai Meiyan Biotechnology Co., Ltd. Cell-line identity was authenticated with short tandem repeat (STR) profiling by the supplier. Cells were maintained in RPMI-1640 medium supplemented with 10% fetal bovine serum and 1% penicillin–streptomycin at 37 °C in a humidified atmosphere containing 5% CO₂. Cells in logarithmic growth phase were used for experiments.

### Section 2. EGFR tyrosine kinase inhibitors and drug preparation

Fourteen EGFR tyrosine kinase inhibitors (EGFR-TKIs) were used in this study. Gefitinib (S31076) and erlotinib (S43873) were obtained from Beijing Zhongkai Boya Technology Co., Ltd. Afatinib (HY-10261), neratinib (HY-32721), osimertinib (HY-15772), dacomitinib (HY-13272), olmutinib (HY-19730), icotinib (HY-15164A), almonertinib (HY-112823), lapatinib (HY-50898), furmonertinib (HY-112870), rezivertinib (HY-109189), befotertinib (HY-137433), and oritinib (HY-139920) were obtained from Shanghai Haoyuan Chemexpress Co., Ltd. All compounds had a reported purity of >99%.

Primary stock solutions of all EGFR-TKIs were prepared at 10 mM in dimethyl sulfoxide (DMSO) and stored at 4 °C. Subsequent working solutions were prepared fresh before use by stepwise dilution in complete culture medium, with the DMSO concentration maintained at 0.1% (v/v). For combination treatments containing n inhibitors, each constituent inhibitor was individually prepared as an n-fold concentrated pre-working solution in complete culture medium containing 0.1% DMSO. Equal volumes of the n constituent pre-working solutions were then combined, yielding the intended final concentration of each inhibitor while maintaining a final DMSO concentration of 0.1% (v/v). Vehicle-control medium contained 0.1% DMSO without drug.

### Section 3. Cell viability and concentration–response analyses

Cell viability was assessed using the Cell Counting Kit-8 (CCK-8; Solarbio, Beijing, China; Cat. No. CA1210-500T) assay. Cells in logarithmic growth were dissociated with 0.25% trypsin–EDTA, resuspended as single-cell suspensions and seeded in 96-well plates at 2,000 cells per well in 100 μl complete culture medium. After 24 h, cells were exposed to the indicated concentrations of individual EGFR-TKIs or drug combinations for 96 h. Six technical replicate wells were used for each treatment condition in each independent experiment. Vehicle-treated control wells and cell-free blank wells were included in the assay. Cell-based experiments were independently repeated three times unless otherwise indicated. CCK-8 reagent (10 μl per well) was then added, and plates were incubated for 1 h at 37 °C. Absorbance was measured at 450 nm using a microplate reader. Cell viability and growth inhibition were calculated after subtraction of the corresponding blank absorbance as follows:

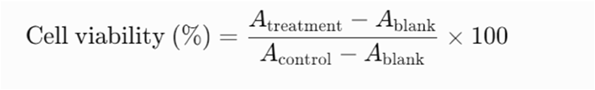

and

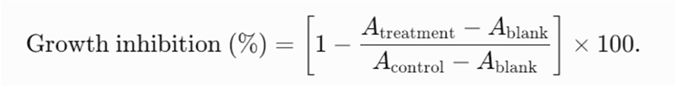

Concentration–response curves for individual EGFR-TKIs (Supplementary Figs. 2, 4, 6, 7–9 and Supplementary Table 1) were generated using GraphPad Prism (version 8.3).

Concentrations corresponding to predefined levels of growth inhibition were derived from the fitted concentration–response curves for each individual inhibitor in the respective cell model. IC₀₅, IC₁₀, IC₁₅ and IC₂₀ concentrations for each inhibitor were used to define the ultra-low-dose conditions for subsequent single-agent and combination experiments. Other effect levels were derived for specific control conditions.

### Section 4. Construction of ultra-low-dose same-target combinations and matched-inhibition conditions

Ultra-low-dose same-target combinations (ULDCs) were constructed by combining multiple EGFR-TKIs, with each constituent inhibitor used at a predefined low-effect concentration derived from its individual concentration–response curve. For the five-drug ULDC comprising gefitinib, erlotinib, afatinib, neratinib, and osimertinib, each inhibitor was used at its respective IC₀₅, IC₁₀, IC₁₅, or IC₂₀ concentration, corresponding to the concentration that produced approximately 5%, 10%, 15%, or 20% growth inhibition when administered individually. Thus, an IC₁₅ five-drug ULDC, for example, contained all five inhibitors simultaneously, with each inhibitor present at its own IC₁₅ concentration. The specific concentrations used for individual inhibitors and combinations are provided in Supplementary Table 2.

Combination working solutions were prepared so that each constituent inhibitor reached its designated final concentration after mixing. For the five-drug combinations, equal volumes of fivefold-concentrated solutions of the individual inhibitors were mixed, yielding the specified final concentration of each component in the combination. The same principle was applied when constructing combinations at other predefined IC levels.

For experiments requiring comparison of treatments at approximately matched levels of growth inhibition, treatment concentrations were selected on the basis of experimentally measured growth inhibition such that the compared single-agent and combination conditions produced approximately 50% growth inhibition.

In repeated cyclic exposure experiments, single-agent and combination treatments were adjusted to produce approximately 50% growth inhibition before repeated exposure.

Similarly, for transcriptomic profiling, individual EGFR-TKIs were used at concentrations producing approximately 50% growth inhibition, whereas the five-drug ULDC was formulated with each EGFR-TKI at its respective IC₁₅ concentration to achieve a comparable overall inhibitory effect.

These approximately matched levels of growth inhibition were used to minimize differences attributable simply to growth-inhibitory strength when comparing biological responses across treatment conditions.

### Section 5. Assessment of same-target collective growth-inhibitory activity

To assess the collective growth-inhibitory activity produced by combining individually weak same-target perturbations in the primary cell model, PC-9 cells were treated with gefitinib, erlotinib, afatinib, neratinib, or osimertinib individually, or with the corresponding five-drug ULDC. Individual inhibitors were tested at their respective IC₀₅, IC₁₀, IC₁₅, and IC₂₀ concentrations. For each ULDC condition, all five inhibitors were administered simultaneously, with each constituent present at the same predefined IC level used for the corresponding single-agent treatment. Thus, the IC₀₅, IC₁₀, IC₁₅, and IC₂₀ ULDC conditions contained five inhibitors, each at its own IC₀₅, IC₁₀, IC₁₅, or IC₂₀ concentration, respectively. The specific concentrations used are provided in Supplementary Table 2. Following 96 h of treatment, cell viability was quantified using the CCK-8 assay. Growth inhibition induced by each five-drug ULDC was compared with the growth inhibition induced by each constituent inhibitor administered individually at the corresponding IC level.

The collective growth-inhibitory activity of the five-drug ULDC was further evaluated in HCC827 and A549 cells at IC₁₀ and IC₂₀ dose levels (Supplementary Tables 8 and 9), with the respective concentrations determined independently in each cell model.

### Section 6. Drop-one analysis of individual inhibitor contributions

To assess the contribution of individual inhibitors to the activity of the five-drug ULDC, a series of drop-one combinations was generated in which one constituent EGFR-TKI was omitted at a time. Drop-one combinations were prepared for gefitinib, erlotinib, afatinib, neratinib, and osimertinib at the predefined IC₀₅, IC₁₀, IC₁₅, and IC₂₀ levels. For each drop-one condition, the concentration of each of the four remaining inhibitors was maintained at the same level as in the corresponding five-drug ULDC. The solution containing the omitted inhibitor was replaced with an equal volume of complete culture medium, thereby maintaining the same final treatment volume without altering the concentrations of the remaining constituents. Thus, differences between the complete five-drug ULDC and each four-drug subset reflected removal of a single inhibitor rather than dilution of the remaining components. PC-9 cells were treated with the complete five-drug ULDC or the corresponding drop-one combinations for 96 h, and cell viability and growth inhibition were determined using the CCK-8 assay as described above.

Growth inhibition induced by each four-drug subset was compared with the growth inhibition induced by the complete five-drug ULDC at the corresponding IC level.

### Section 7. Combination index analysis

Drug interactions within the ULDC were additionally evaluated using the combination index (CI) framework of Chou and Talalay [19,20]. For each combination condition, the experimentally observed growth inhibition was used to determine the concentration of each inhibitor required to produce the same effect when administered alone, based on the concentration–response curve of the individual EGFR-TKI. These effect-equivalent single-agent concentrations were compared with the corresponding concentrations present in the combination to calculate the CI (Supplementary Tables 5, 6, 12, 16 and 18 and Supplementary Figs. 5 and 17). CI values were interpreted according to the conventional Chou–Talalay framework, with CI values below, approximately equal to, or above 1 interpreted as synergistic, additive or antagonistic interactions, respectively. CI analyses were performed for the predefined ultra-low-dose combination conditions within the IC_05_–IC_20_ range.

### Section 8. Scaling same-target inhibitor diversity

To examine the effect of increasing same-target inhibitor diversity, EGFR-TKI combinations containing 5, 8, 10, or 14 inhibitors were evaluated in PC-9 cells. The five-drug ULDC comprised gefitinib, erlotinib, afatinib, neratinib, and osimertinib (inhibitors 1–5). Higher-order combinations were generated by sequentially extending this inhibitor set: the eight-drug ULDC comprised inhibitors 1–8, the ten-drug ULDC comprised inhibitors 1–10, and the fourteen-drug ULDC comprised inhibitors 1–14. The identity of all inhibitors and the corresponding constituent concentrations are provided in Supplementary Table 11. For the higher-order ULDC experiments, constituent inhibitors were used at their respective IC₀₅ or IC₁₀ concentrations, as specified in Supplementary Table 11.

### Section 9. Repeated cyclic drug exposure and assessment of the emergence of resistance

Repeated cyclic drug exposure experiments were performed in PC-9 cells treated with ten individual EGFR-TKIs, the five-drug ULDC, or higher-order ULDCs containing 8, 10, or 14 inhibitors to evaluate the emergence of resistance. Before repeated cyclic drug exposure, treatment concentrations were optimized to produce approximately matched levels of growth inhibition (∼50%) in parental PC-9 cells. For the five-drug ULDC, each constituent EGFR-TKI was administered at approximately its IC₁₅ concentration, whereas the concentrations of the 8-, 10-, and 14-drug ULDCs were proportionally adjusted from their initial IC₀₅- or IC₁₀-based combinations to achieve comparable growth inhibition (Supplementary Fig. 14). The optimized treatment concentrations are summarized in Supplementary Tables 13 and 14. These approximately matched growth-inhibition conditions were designed to minimize potential confounding arising from differences in growth-inhibitory strength, thereby enabling resistance outcomes to be compared across combinations containing different numbers of inhibitors.

Each treatment cycle consisted of 48 h of drug exposure followed by 48 h of drug-free recovery. This four-day cycle was repeated under identical culture conditions. Cell viability was assessed after 5, 10, 15, 20, 25, 30, 35, and 40 exposure cycles using the CCK-8 assay. No measurements were performed during the intervening exposure cycles. At each predefined assessment point, cell viability of repeatedly exposed cells was compared with that of parental PC-9 cells evaluated in parallel under identical assay conditions. Resistance was considered to have emerged when cell viability was significantly higher than that of the parental PC-9 cells (P < 0.05).

### Section 10. Assessment of ULDC activity in gefitinib-resistant PC-9/GR cells

The activity of ULDCs against resistant cells was evaluated in gefitinib-resistant PC-9/GR cells. Gefitinib, erlotinib, afatinib, neratinib, and osimertinib were tested individually at the same predefined IC₀₅, IC₁₀, IC₁₅, and IC₂₀ concentrations established in parental PC-9 cells. The corresponding five-drug ULDCs were constructed at each concentration level, with each constituent inhibitor present at the same concentration used for the corresponding single-agent treatment. The specific treatment concentrations are provided in Supplementary Table 2. PC-9/GR cells were exposed to the indicated treatments for 96 h, after which cell viability and growth inhibition were determined using the CCK-8 assay. The activity of the five-drug ULDC was evaluated relative to the activity of the corresponding individual EGFR-TKIs at each predefined concentration level.

### Section 11. Assessment of the reversibility of resistance

To assess the reversibility of resistance states arising during repeated drug exposure, resistant PC-9 cell populations derived from single-agent EGFR-TKI treatment or the five-drug ULDC were cultured in drug-free complete medium. For the single-agent groups, resistant populations were generated after 15 cycles of exposure to gefitinib, erlotinib, afatinib, neratinib, or osimertinib. The ULDC-selected resistant populations were obtained after 40 cycles of exposure to the five-drug ULDC.

Cells were maintained in drug-free medium for five consecutive recovery periods, with each recovery period corresponding to two cell passages. Drug sensitivity was reassessed after each recovery period (passages 2, 4, 6, 8, and 10) by re-exposing cells to the same treatment conditions used during the original repeated cyclic exposure experiments, corresponding to approximately matched levels of growth inhibition (∼50%) in parental PC-9 cells. Cell viability was then determined using the CCK-8 assay. For each treatment group, changes in drug sensitivity during recovery were evaluated relative to the corresponding resistant population before drug-free recovery and with reference to the response of parental PC-9 cells under the same assay conditions. Recovery of drug sensitivity toward parental PC-9 levels was used to assess the reversibility of the resistant state.

### Section 12. RNA sequencing and transcriptomic profiling

PC-9 cells were treated with gefitinib, erlotinib, afatinib, neratinib, osimertinib, or the five-drug ULDC under approximately matched levels of growth inhibition (∼50%). Untreated PC-9 cells (in drug-free complete medium) served as controls. Three independent biological replicates were analysed for each condition. Total RNA was extracted using TRIzol reagent according to the manufacturer’s instructions. RNA concentration and integrity were assessed using a NanoDrop spectrophotometer (Thermo Fisher Scientific) and an Agilent 5400 Bioanalyzer (Agilent Technologies), respectively. Samples meeting RNA quality requirements were used for library preparation and transcriptome sequencing, which were performed by BGI Genomics Co., Ltd. (Shenzhen, China). Raw sequencing reads were filtered using SOAPnuke to remove adapter-contaminated reads, reads containing >5% unknown bases, and low-quality reads, defined as reads in which >20% of bases had quality scores below 15. Clean reads were aligned to the human reference genome (NCBI, GCF_000001405.39_GRCh38.p13) using HISAT2. Subsequent transcriptomic analyses were performed as described below.

### Section 13. Differential expression and functional enrichment analyses

Gene expression levels were quantified by aligning clean reads to the reference gene set using Bowtie2, followed by transcript abundance estimation using RSEM. Differential expression analysis between each treatment condition and the untreated control was performed using DESeq2. Genes with an absolute log2 fold change >1 and a Q value <0.05 were considered differentially expressed genes (DEGs). Principal component analysis (PCA) was performed to assess the overall similarity and separation of transcriptomic profiles across treatment groups. Intersections among DEG sets identified for the five individual EGFR-TKIs and the five-drug ULDC were visualized using UpSet analysis to identify shared and treatment-associated transcriptional responses. KEGG pathway enrichment analysis was performed on the combination-associated DEGs identified for the five-drug ULDC. Enrichment significance was evaluated using the hypergeometric test, and pathways with a Q value <0.05 were considered significantly enriched.

### Section 14. RT–qPCR validation of RNA-seq findings

Representative differentially expressed genes identified by RNA sequencing were selected for validation by reverse transcription quantitative PCR (RT–qPCR). Total RNA was extracted from PC-9 cells using TRIzol reagent and reverse-transcribed into cDNA using a first-strand cDNA synthesis kit.

RT–qPCR reactions were performed in a total volume of 20 μL containing 10.0 μL of 2× SYBR Green PCR mix, 0.4 μL each of forward and reverse primers, 1.0 μL of cDNA template, and 8.2 μL of ddH₂O. Amplification was performed using the following cycling conditions: 50 °C for 2 min, 94 °C for 10 min, 95 °C for 3 min, followed by 40 cycles of 95 °C for 5 s and 60 °C for 30 s. GAPDH was used as the internal reference gene.

Relative gene expression levels were calculated using the 2^⁻ΔΔCt^ method. Primer sequences used for RT–qPCR validation are provided in Supplementary Table 19.

### Section 15. In vivo antitumour efficacy study

Female and male SPF-grade BALB/c nude mice (4 weeks old, 15–18 g) were obtained from the Laboratory Animal Center of Peking University Health Science Center and maintained under specific pathogen-free (SPF) conditions. The animal experiments reported in this study were conducted without prior approval from an institutional animal ethics committee, as an ethics review application had not been submitted before the experiments were initiated.

Subcutaneous xenograft tumours were established by injecting 1 × 10⁶ PC-9 cells suspended in 0.1 mL PBS into the right axillary region of each mouse. When tumour volumes reached approximately 50–100 mm³, mice were randomly assigned to nine experimental groups, including a vehicle control group, five individual ultra-low-dose EGFR-TKI groups, a five-drug ULDC group, a positive-control gefitinib group and a non-tumour-bearing control group. Each group contained ten mice (five females and five males).

Ultra-low doses were defined with reference to the World Health Organization (WHO) defined daily dose (DDD) for each EGFR-TKI. Mouse doses were converted to human-equivalent doses (HEDs) using body-surface-area scaling based on standard Km factors (mouse, 3; adult human, 37) [40]. Based on a reference human body weight of 60 kg, the resulting HEDs corresponded to 9.7%, 9.7%, 9.7%, 9.7%, and 4.9% of the respective WHO-defined DDDs for gefitinib, erlotinib, afatinib, neratinib, and osimertinib, respectively [41]. Individual ultra-low-dose EGFR-TKIs were administered by oral gavage at 5 mg/kg gefitinib, 3 mg/kg erlotinib, 0.8 mg/kg afatinib, 4.8 mg/kg neratinib, and 0.8 mg/kg osimertinib. The five-drug ULDC consisted of these five inhibitors administered simultaneously at the same constituent doses. A positive-control group received gefitinib at 25 mg/kg. Treatments were administered for three consecutive weeks according to a schedule of five treatment days followed by two drug-free days per week.

Body weights were recorded daily throughout the treatment period. Tumour dimensions were measured every two days using digital calipers. Tumour volume was calculated as V = L × W²/2, where L and W represent the longest and shortest tumour diameters, respectively. Tumour volume ratios were calculated relative to tumour volume measured on the first day of treatment and were used to generate tumour growth curves (Fig. 5). At the completion of the study, mice were euthanized, and tumours were excised, rinsed with physiological saline, blotted dry, and weighed. Major organs together with serum samples were collected for subsequent histopathological, immunohistochemical and biochemical analyses.

### Section 16. Tissue collection and histopathological analysis

Tumour tissues and major organs, including the heart, liver, spleen, lung, and kidneys, were fixed in 4% paraformaldehyde, routinely processed, embedded in paraffin and sectioned at 6–7 μm thickness. For histopathological evaluation, paraffin sections were stained with haematoxylin and eosin (H&E) according to standard procedures and examined by light microscopy for treatment-associated histopathological alterations.

Histopathological evaluation was performed on tumour tissues and major organs, including the heart, liver, spleen, lung, and kidneys. Representative H&E-stained histological images are presented in Supplementary Figs. 24 and 25.

### Section 17. Immunohistochemistry

Paraffin-embedded tumour sections prepared as described above were deparaffinized and rehydrated, followed by heat-induced antigen retrieval in EDTA antigen-retrieval buffer (pH 9.0) using microwave heating (medium power for 8 min, an 8-min interval without heating, followed by medium-low power for 7 min). Sections were cooled to room temperature and washed with PBS. Endogenous peroxidase activity was blocked with 3% H₂O₂ for 25 min at room temperature in the dark, followed by blocking with 3% bovine serum albumin (BSA) for 30 min at room temperature.

Sections were incubated overnight at 4 °C with rabbit anti-p-EGFR antibody (Zenbio, Cat. No. 310055; 1:100) or mouse anti-Ki-67 antibody (Servicebio, Cat. No. GB121141; 1:600). After washing with PBS, sections were incubated for 50 min at room temperature with HRP-conjugated goat anti-rabbit IgG (Servicebio, Cat. No. GB23303; 1:200) or HRP-conjugated goat anti-mouse IgG (Servicebio, Cat. No. GB23301; 1:200), as appropriate. Immunoreactivity was visualized using a DAB detection kit (Servicebio, Cat. No. G1212), followed by haematoxylin counterstaining. Sections were subsequently dehydrated, cleared, mounted, and examined using a Nikon E100 light microscope.

Positive immunoreactivity was visualized as brown staining, with nuclei counterstained blue. Representative p-EGFR and Ki-67 immunohistochemical images are shown in Fig. 5.

### Section 18. Serum biochemical analyses

At the completion of the treatment period, mice were anesthetized with isoflurane, and whole blood was collected by ocular enucleation. Mice were euthanized immediately after blood collection. Blood samples were allowed to clot at room temperature, and serum was obtained by centrifugation at 3,500 rpm for 10 min and stored at −80℃ until analysis.

Serum biochemical parameters, including alanine aminotransferase (ALT), aspartate aminotransferase (AST), albumin (ALB), total protein (TP), creatinine (CREA), urea, uric acid (UA), creatine kinase (CK), lactate dehydrogenase (LDH), and glucose (GLU), were measured using an automated biochemical analyzer according to the manufacturer’s instructions.

### Section 19. Statistical analysis

Data are presented as mean ± standard deviation (SD), unless otherwise indicated. For cell-based experiments, n represents the number of independent experiments rather than the number of technical replicate wells. Unless otherwise indicated, cell-based experiments were independently repeated three times, with six technical replicate wells per treatment condition for each independent experiment. Statistical analyses were performed using GraphPad Prism 8.3. Comparisons among multiple groups were performed using analysis of variance (ANOVA), followed by appropriate multiple-comparison testing where applicable. Statistical tests used for individual experiments are specified in the corresponding figure legends. A two-sided P value <0.05 was considered statistically significant. For repeated cyclic drug exposure experiments, the emergence of resistance was assessed at predefined time points by comparing cell viability of repeatedly exposed cells with the viability of parental PC-9 cells under identical assay conditions. Resistance was defined as significantly higher cell viability compared with the parental PC-9 cells (P < 0.05), indicating reduced drug sensitivity following repeated cyclic exposure.

## Data Availability and Code Availability statements

The data presented in the main manuscript and Supplementary Information support the findings reported in this study.

## Acknowledgements

The conceptual foundations of this study developed from over three decades of research by our group on the pharmacologically active substances and mechanisms of action of traditional Chinese medicines. We gratefully acknowledge the long-term support of the research programmes from which these concepts and the experimental framework of the present study emerged. These research programmes received support from the National Outstanding Youth Science Foundation Project of the National Natural Science Foundation of China (30425018 to S.-Q.C.); Key Projects of the National Natural Science Foundation of China (30330710 to X.W. and S.-Q.C.; 30830120 to S.-Q.C. and F.X.); National Natural Science Foundation of China projects (30371721 to S.-Q.C. and F.X.; 81274073 to S.-Q.C. and F.X.; 81673595 to S.-Q.C. and F.X.; 81573593 to F.X.; and 81973472 to F.X.); National 863 Plan Project of the Ministry of Science and Technology of China (2004AA2Z3730-05 to S.-Q.C.); National 973 Project of the Ministry of Science and Technology of China (2006CB504707 to S.-Q.C. and X.W.); Major National Science and Technology Project for the Creation of Major New Drugs of the Ministry of Science and Technology of China (2009ZX09301010-P08 to S.-Q.C., X.W., and F.X.; 2009ZX09502-023 to S.-Q.C., F.X., and X.W.; 2013ZX09508104 to S.-Q.C.; and 2019ZX09201004 to F.X.); National TCM Standardization Project of the National Development and Reform Commission of China (ZYBZH-C-JS-35 to X.W.; ZYBZH-Y-SC-41 to R.T. and S.-Q.C.; and ZYBZH-Y-TJ-43 to X.W.); Beijing TCM Science and Technology Development Fund Project (JJ2016-78 to S.-Q.C. and F.X.); and Peking University Health Science Center 985 Discipline Construction Fund (BMU20130358 to S.-Q.C., X.W., and F.X.).

## Author Contributions

**Conceptualization**: S.-Q.C., R.T., and F.X.; **Funding acquisition**: S.-Q.C., F.X., X.W., and R.T.; **Supervision**: R.T., S.-Q.C., and F.X.; **Project administration**: S.-Q.C., R.T., F.X., and Z.-G.Z.; **Investigation**: Z.-G.Z., F.X., and Q.Z.; **Resources**: Z.-G.Z., W.C., Q.Z., and X.W.; **Methodology**: Z.-G.Z., S.-Q.C., and F.X.; **Visualization**: Z.-G.Z., Q.Z., W.C., and F.X.; **Data curation**: Z.-G.Z., F.X., W.C., R.T., and S.-Q.C.; **Formal analysis**: Z.-G.Z., W.C., S.-Q.C., and R.T.; **Validation**: Z.-G.Z., F.X., R.T., S.-Q.C., and W.C.; **Writing – original draft**: S.-Q.C., Z.-G.Z., and W.C.; **Writing – review & editing**: S.-Q.C., Z.-G.Z., F.X., and R.T.

## Competing interests

The authors declare the following competing interests: S.-Q.C. is the applicant, and S.-Q.C., Z.-G.Z., F.X., R.T., and X.W. are inventors on Chinese patent application no. 202511888023.5 (publication no. CN 121445747 A; published on 3 February 2026), relating to ultra-low-dose combinations of multiple EGFR inhibitors for the treatment of non-small-cell lung cancer, including aspects of their antitumour activity and resistance-related effects reported in this manuscript. The remaining authors declare no competing interests.

**Supplementary Information** is available for this paper.

## REFERENCES

1. Li, J., Hu, J., Yang, Y., et al. Drug resistance in cancer: molecular mechanisms and emerging treatment strategies. Mol. Biomed. 6, 111 (2025).

2. Dagogo-Jack, I. & Shaw, A. T. Tumour heterogeneity and resistance to cancer therapies. Nat. Rev. Clin. Oncol. 15 (2), 81–94 (2018).

3. Marine, J.-C., Dawson, S.-J., & Dawson, M. A. Non-genetic mechanisms of therapeutic resistance in cancer. Nat. Rev. Cancer 20 (12), 743–756 (2020).

4. Jin, H., Wang, L., & Bernards, R. Rational combinations of targeted cancer therapies: background, advances and challenges. Nat. Rev. Drug Discov. 22 (3), 213–234 (2023).

5. Obenauf, A. C. Mechanism-based combination therapies for metastatic cancer. Sci. Transl. Med. 14 (655), eadd0887 (2022).

6. Khan, S.U., Fatima, K., Aisha, S., & Malik, F. Unveiling the mechanisms and challenges of cancer drug resistance. Cell Commun. Signal. 22, 109 (2024).

7. Rezayatmand, H., Razmkhah, M., & Razeghian-Jahromi, I. Drug resistance in cancer therapy: the Pandora’s Box of cancer stem cells. Stem Cell Res. Ther. 13, 181 (2022).

8. Herbst, R. S., Morgensztern, D., & Boshoff, C. The biology and management of non-small cell lung cancer. Nature 553 (7689), 446–454 (2018).

9. Wang, M., Herbst, R. S., & Boshoff, C. Toward personalized treatment approaches for non-small-cell lung cancer. Nat. Med. 27 (8), 1345–1356 (2021).

10. Soria, J.-C., Ohe, Y., Vansteenkiste, J. et al. Osimertinib in untreated EGFR-mutated advanced non-small-cell lung cancer. N. Engl. J. Med. 378 (2), 113–125 (2018).

11. Ramalingam, S. S., Vansteenkiste, J., Planchard, D. et al. Overall survival with osimertinib in untreated EGFR-mutated NSCLC. N. Engl. J. Med. 382 (1), 41–50 (2020).

12. Mok, T. S., Wu, Y.-L., Ahn, M.-J. et al. Osimertinib or platinum-pemetrexed in EGFR T790M-positive lung cancer. N. Engl. J. Med. 376 (7), 629–640 (2017).

13. Leonetti, A., Sharma, S., Minari, R. et al. Resistance mechanisms to osimertinib in EGFR-mutated non-small cell lung cancer. Br. J. Cancer 121 (9), 725–737 (2019).

14. Choudhury, N. J., Marra, A., Sui, J. et al. Molecular biomarkers of disease outcomes and mechanisms of acquired resistance to first-line osimertinib in advanced EGFR-mutant lung cancers. J. Thorac. Oncol. 18 (4), 463–475 (2023).

15. Wu, Y.-L., Giaccone, G., Hayashi, H. et al. Tepotinib plus osimertinib in patients with EGFR-mutated non-small-cell lung cancer with MET amplification following progression on first-line osimertinib (INSIGHT 2): a multicentre, open-label, phase 2 trial. Lancet Oncol. 25 (8), 989–1002 (2024).

16. Greaves, M. & Maley, C. C. Clonal evolution in cancer. Nature 481 (7381), 306– 313 (2012).

17. Laplane, L. & Maley, C.C. The evolutionary theory of cancer: challenges and potential solutions. Nat. Rev. Cancer 24, 718–733 (2024).

18. Wang, Y., Xu, F., Zhao, Q. et al. Combining same-target drugs exhibits additive efficacy with minimal toxicity [preprint]. Research Square (2024): 10.21203/rs.3.rs-4822017/v1

19. Chou, T. C. Drug combination studies and their synergy quantification using the Chou-Talalay method. Cancer Res. 70 (2), 440–446 (2010).

20. Chou, T. C. Theoretical basis, experimental design, and computerized simulation of synergism and antagonism in drug combination studies. Pharmacol. Rev. 58 (3), 621–681 (2006).

21. Liu, B., Chen, D., Chen S., et al. Transcriptional activation of cyclin D1 via HER2/HER3 contributes to EGFR-TKI resistance in lung cancer. Biochem. Pharmacol. 178, 114095 (2020).

22. Fu, M., Zhao, J., Zhang, L., et al. Overcoming tyrosine kinase inhibitor resistance in lung cancer brain metastasis with CTLA4 blockade. Cancer Cell 42 (11), 1882–1897 (2024).

23. Ma, Y., Feng, Q., Han, B., et al. Elevated HMGB1 promotes the malignant progression and contributes to cisplatin resistance of non-small cell lung cancer. Hereditas 160 (1), 33 (2023).

24. Castro, M.A.A., Dal-Pizzol, F., Zdanov, S., et al. CFL1 expression levels as a prognostic and drug resistance marker in non-small cell lung cancer. Cancer 116 (15), 3645–3655 (2010).

25. Eck, M. J. & Yun, C.-H. Structural and mechanistic underpinnings of the differential drug sensitivity of EGFR mutations. Cold Spring Harb. Perspect. Med. 10 (11), a036889 (2020).

26. Yun, C.-H., Boggon, T. J., Li, Y., et al. Structures of lung cancer-derived EGFR mutants and inhibitor complexes: mechanism of activation and insights into differential inhibitor sensitivity. Cancer Cell 11 (3), 217–227 (2007).

27. Zhang, X., Gureasko, J., Shen, K. et al. An allosteric mechanism for activation of the EGFR kinase domain of EGFR. Cell 125 (6), 1137–1149 (2006).

28. Jura, N., Endres, N. F., Engel, K. et al. Mechanism for activation of the EGF receptor catalytic domain by the juxtamembrane segment. Cell 137 (7), 1293–1307 (2009).

29. Thress, K. S., Paweletz, C. P., Felip, E. et al. Acquired EGFR C797S mutation mediates resistance to AZD9291 in non-small cell lung cancer harboring EGFR T790M. Nat. Med. 21 (6), 560–562 (2015).

30. Niederst, M. J., Hu, H., Mulvey, H. E. et al. The allelic context of the C797S mutation acquired upon treatment with third-generation EGFR inhibitors impacts sensitivity to subsequent treatment strategies. Clin. Cancer Res. 21 (17), 3924–3933 (2015).

31. Fernandes Neto, J. M., et al. Multiple low dose therapy as an effective strategy to treat EGFR inhibitor-resistant NSCLC tumours. Nat. Commun. 11, 3157 (2020).

32. Modi, V. & Dunbrack, R. L. Defining a new nomenclature for the structures of active and inactive kinases. Proc. Natl. Acad. Sci. USA 116 (14), 6818–6827 (2019).

33. Roskoski, R. Classification of small molecule protein kinase inhibitors based upon the structures of their drug–enzyme complexes. Pharmacol. Res. 103, 26–48 (2016).

34. Hata, A. N., Niederst, M. J., Archibald, H. L. et al. Tumor cells can follow distinct evolutionary paths to become resistant to EGFR inhibition. Nat. Med. 22 (3), 262–269 (2016).

35. Corcoran, R. B., et al. Combined BRAF, EGFR, and MEK inhibition in patients with *BRAF*^V600E^-mutant colorectal cancer. Cancer Discov. 8, 428–443 (2018).

36. Hazar-Rethinam, M., et al. Convergent therapeutic strategies to overcome the heterogeneity of acquired resistance in *BRAF*^V600E^ colorectal cancer. Cancer Discov. 8, 417–427 (2018).

37. West, J., You, L., Zhang, J. et al. Towards multidrug adaptive therapy. Cancer Res. 80 (7), 1578–1589 (2020).

38. West, J., Adler, F., Gallaher, J. et al. A survey of open questions in adaptive therapy: bridging mathematics and clinical translation. eLife 12, e84263 (2023).

39. Nowell, P. C. The clonal evolution of tumor cell populations: Acquired genetic lability permits stepwise selection of variant sublines and underlies tumor progression. Science 194 (4260), 23–28 (1976).

40. U.S. Food and Drug Administration. Guidance for Industry: Estimating the Maximum Safe Starting Dose in Initial Clinical Trials for Therapeutics in Adult Healthy Volunteers (2005). https://www.fda.gov/files/drugs/published/Estimating-the-Maximum-Safe-Starting-Dose-in-Initial-Clinical-Trials-for-Therapeutics-in-Adult-Healthy-Volunteers.pdf

41. WHO Collaborating Centre for Drug Statistics Methodology. ATC classification index with DDDs, 2026. Oslo, Norway (2026). https://atcddd.fhi.no/atc_ddd_index/

