## Supplementary Information for "Ultra-low-dose same-target combinations control drug resistance across stages"

### ABSTRACT

The emergence of drug resistance continues to limit the long-term effectiveness of cancer therapies [1], as tumour cells adapt to therapeutic selective pressure through diverse genetic and non-genetic routes [2,3]. Despite advances in more potent and next-generation inhibitors and mechanism-based combination therapies [4,5], durable disease control remains difficult to achieve. Here we tested an alternative strategy: combining multiple inhibitors directed against the same oncogenic driver, each at ultra-low doses with minimal individual activity, to delay resistance emergence. In epidermal growth factor receptor (EGFR)-driven cancer models, such ultra-low-dose same-target combinations (ULDCs) distributed individually weak perturbations across multiple inhibitors directed against the same target and achieved substantial growth inhibition in vitro and antitumour activity in vivo despite weak effects of the corresponding single agents. Under similar levels of growth inhibition (~50%), single-agent EGFR inhibitors generated resistant outgrowth, whereas a five-drug ULDC delayed resistance emergence. Increasing the number of inhibitors further delayed resistance emergence, with no detectable resistant outgrowth observed for combinations containing 8–14 inhibitors within the experimental timeframe. The five-drug ULDC also retained measurable growth-inhibitory activity against gefitinib-resistant PC-9/GR cells, and resistant cells arising under five-drug ULDC treatment exhibited markedly greater reversibility following drug withdrawal than those arising under single-agent treatment. Although pharmacological additivity or synergy may contribute to the collective growth-inhibitory effects of ULDCs, the divergent long-term resistance outcomes observed under similar levels of growth inhibition indicate that resistance behaviour is not determined by inhibitory strength alone. Instead, our findings support a model in which increasing same-target inhibitor diversity progressively constrains the adaptive trajectories available during resistance evolution, while therapeutic perturbation architecture may also influence the stability of resistant states that emerge.

More broadly, these findings support the concept of same-target collective suppression, in which multiple individually weak perturbations directed against the same target can collectively produce substantial antitumour activity and constrain resistance evolution across multiple stages. We propose that increasing same-target inhibitor diversity may restrict accessible adaptive trajectories, while therapeutic perturbation architecture may also influence the stability of resistant states.

### CONTENT


#### Supplementary Note 1. Experimental characterization of ultra-low-dose same-target combinations

1.1 Collective growth inhibition by five-drug ULDC

1.1.1 Concentration–response characterization of EGFR-TKIs in PC-9 cells

1.1.2 Collective growth inhibition by five-drug ULDCs in PC-9 cells

1.1.3 Combination index analysis of five-drug ULDCs in PC-9 cells

1.1.4 Distributed contribution within five-drug ULDC

1.1.5 Validation across additional cellular contexts

1.1.5.1 Validation in HCC827 cells

1.1.5.2 Collective growth inhibition in non-EGFR-driven cellular contexts

1.2 Scaling of collective growth inhibition with increasing same-target inhibitor diversity

1.2.1 Expanded EGFR-TKI panel

1.2.2 Increasing same-target diversity progressively enhances collective growth inhibition

1.2.3 Combination index analysis across increasing inhibitor numbers

#### Supplementary Note 2. Resistance emergence and resistant-state properties under ULDCs

2.1 Resistance emergence under repeated drug exposure

2.1.1 Approximately matched growth-inhibition conditions for repeated cyclic drug exposure

2.1.2 Five-drug ULDC delays resistance emergence

2.1.3 Increasing same-target inhibitor diversity further delays detectable resistance emergence

2.1.4 Combination index analysis across higher-order ULDCs

2.2 Five-drug ULDC activity in resistant PC-9/GR cells

2.3 Reversibility of resistance states

**Supplementary Note 3. Transcriptomic profiling reveals distinct biological responses induced by the five-drug ULDC**

3.1 The five-drug ULDC induces a distinct transcriptomic profile compared with individual EGFR-TKIs

3.2 The five-drug ULDC produces broader transcriptional changes than individual inhibitors

3.3 ULDC-associated transcriptional responses

3.4 ULDC-induced transcriptional changes are associated with tumour-relevant biological processes

3.5 Validation of RNA-seq findings by RT-qPCR

**Supplementary Note 4. In vivo antitumour activity and tolerability of the five-drug ULDC**

4.1 In vivo collective antitumour activity of the five-drug ULDC

4.2 Reduced EGFR phosphorylation and tumour-cell proliferation in vivo

4.3 In vivo tolerability assessment

**Supplementary Discussion. Conceptual origin of the same-target combination strategy**

### <u>Supplementary Note 1.</u> Experimental characterization of ultra-low-dose same-target combinations

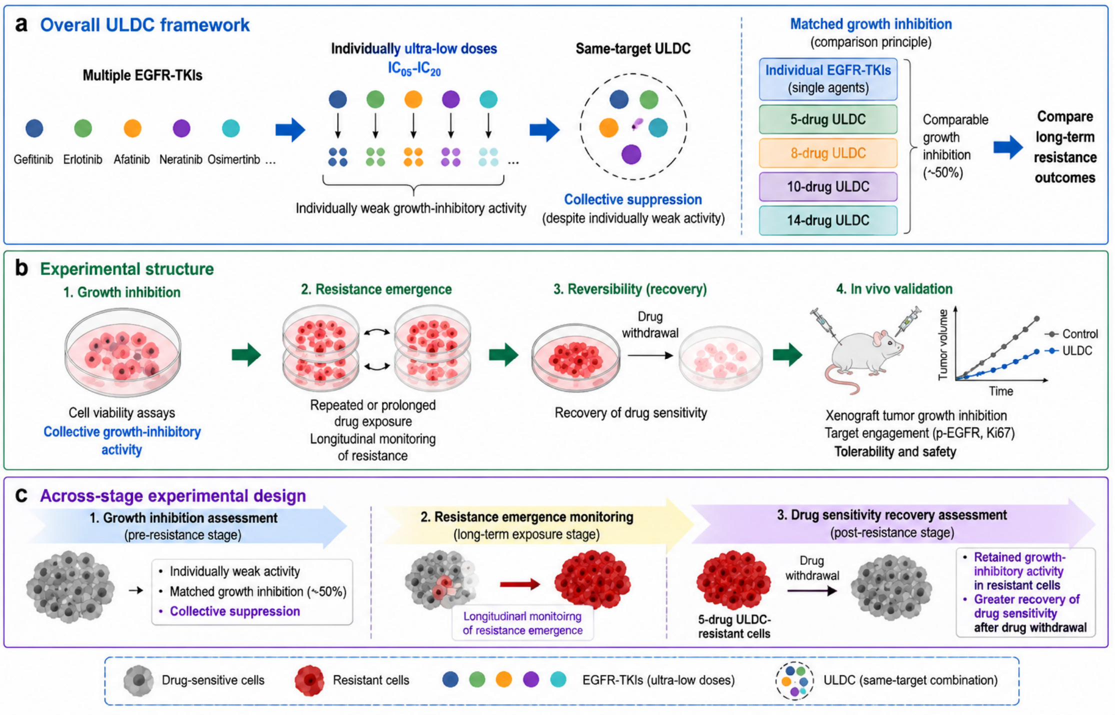


**Supplementary Fig. 1. Overall experimental workflow of the study.** **a**, Schematic of the ultra-low-dose same-target combination (ULDC) framework. Multiple EGFR-TKIs (epidermal growth factor receptor tyrosine kinase inhibitors) with individually weak activity at ultra-low doses (typically $IC_{05}$–$IC_{20}$ concentrations) were combined to generate collective growth inhibition. Approximately matched level of growth inhibition (~50%) was used as a comparison principle to evaluate long-term resistance outcomes across treatment conditions with increasing same-target inhibitor diversity. **b**, Overview of the experimental strategy, including growth inhibition, resistance emergence during repeated or prolonged drug exposure, recovery of drug sensitivity following drug withdrawal, and *in vivo* evaluation of antitumour activity, target inhibition and tolerability. **c**, Across-stage experimental design for evaluating ULDC activity before resistance, during resistance emergence and after resistance establishment, encompassing growth inhibition assessment, longitudinal monitoring of resistance emergence and assessment of drug sensitivity recovery following drug withdrawal.

#### 1.1 Collective growth inhibition by five-drug ULDC

##### 1.1.1 Concentration–response characterization of EGFR-TKIs in PC-9 cells

Concentration–response relationships of individual EGFR-TKIs (gefitinib, erlotinib, afatinib, neratinib, and osimertinib) were first determined in human EGFR-mutant non-small-cell lung cancer (NSCLC) PC-9 cells using CCK-8 assays. Concentrations corresponding to $IC_{05}$, $IC_{10}$, $IC_{15}$, and $IC_{20}$ values were calculated by interpolation from fitted concentration-response curves (Supplementary Fig. 2, Supplementary Tables 1 and 2). These concentrations were subsequently used to construct ultra-low-dose same-target combinations (ULDCs).

Consistent with the concentration–response curves, individual EGFR-TKIs produced only minimal growth inhibition at $IC_{05}$–$IC_{20}$ concentrations. However, when the five inhibitors were combined with each component maintained at the same concentration as in the corresponding single-agent condition, the resulting ULDCs produced substantially greater growth inhibition than any individual inhibitor alone (Fig. 2a).

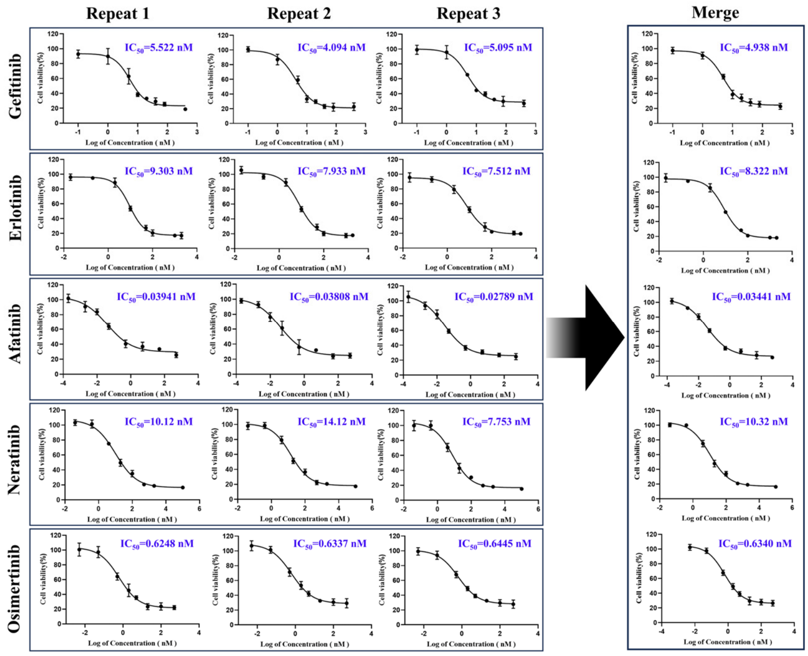


**Supplementary Fig. 2. Concentration–response curves of five EGFR-TKIs in PC-9 cells.** Data are presented as mean ± SD (n = 3).

**Supplementary Table 1. Concentration–response concentration ranges and $IC_{50}$ values of EGFR-TKIs in PC-9 cells**

| Inhibitor | Concentration range (nM) | $IC_{50}$ replicate 1 (nM) | $IC_{50}$ replicate 2 (nM) | $IC_{50}$ replicate 3 (nM) | Fitted $IC_{50}$ (nM) |
|---|---|---|---|---|---|
| Gefitinib | 0.1, 1, 5, 10, 20, 40, 80, 400 | 5.522 | 4.094 | 5.095 | 4.938 |
| Erlotinib | 0.02, 0.2, 2, 10, 50, 100, 1000, 2000 | 9.303 | 7.933 | 7.512 | 8.322 |
| Afatinib | 0.0002, 0.002, 0.01, 0.05, 0.5, 5, 50, 500 | 0.03941 | 0.03808 | 0.02789 | 0.03441 |
| Neratinib | 0.04, 0.4, 4, 20, 100, 500, 2000, 10000 | 10.12 | 7.864 | 14.12 | 10.32 |
| Osimertinib | 0.005, 0.05, 0.5, 2, 5, 20, 100, 500 | 0.6248 | 0.6445 | 0.6334 | 0.6340 |

**Supplementary Table 2. Concentrations of individual EGFR-TKIs and their combinations used in PC-9 and PC-9/GR cell experiments**

| Groups | Concentration（nM） | | | |
|---|---|---|---|---|
| | $IC_{05}$ | $IC_{10}$ | $IC_{15}$ | $IC_{20}$ |
| Gefitinib | 0.5463 | 1.214 | 1.785 | 2.345 |
| Erlotinib | 0.4766 | 1.219 | 1.991 | 2.832 |
| Afatinib | 0.001438 | 0.003210 | 0.005906 | 0.009869 |
| Neratinib | 0.5761 | 1.134 | 1.877 | 2.845 |
| Osimertinib | 0.05841 | 0.1062 | 0.1676 | 0.2462 |
| 5-drug Combination | $IC_{05}$ Combo （1.659） | $IC_{10}$ Combo （3.676） | $IC_{15}$ Combo （5.827） | $IC_{20}$ Combo （8.278） |

Note: The $IC_{05}$, $IC_{10}$, $IC_{15}$, and $IC_{20}$ concentrations of the five EGFR-TKIs were determined individually in PC-9 cells. $IC_{05}$ Combo to $IC_{20}$ Combo represent the summed concentrations of all individual EGFR-TKIs in the corresponding column at the indicated concentration level.

##### 1.1.2 Collective growth inhibition by five-drug ULDCs in PC-9 cells

We next examined whether five EGFR-TKIs produced same-target collective growth inhibition when combined at ultra-low concentrations. At the $IC_{05}$ level, individual inhibitors produced only minimal growth inhibition, ranging from 2.67%–8.81%, close to the theoretical value (5%), whereas the five-drug ULDC produced 33.48% inhibition. Similar collective effects were observed across increasing ultra-low-dose levels, with five-drug ULDCs producing 38.78%, 49.53%, and 62.32% inhibition at $IC_{10}$, $IC_{15}$, and $IC_{20}$, respectively (Supplementary Tables 3 and 4). Combination index (CI) analysis yielded CI values below 1 (0.43–0.63) across all ultra-low-dose levels, indicating synergistic interactions according to the CI framework under these conditions (Supplementary Table 5). Together, these results demonstrate that weak individual EGFR-TKI perturbations can generate substantial collective growth inhibition when combined within the same-target ULDC framework.

**Supplementary Table 3. Collective growth inhibition of five-drug ULDCs in PC-9 cells**

| Condition | Expected inhibition (%) | Observed inhibition of single agents (%) | Observed inhibition of five-drug ULDC (%) | P value vs single-agent groups |
|---|---|---|---|---|
| $IC_{05}$ | 5 | 2.67–8.81 | 33.48 | <0.0001 |
| $IC_{10}$ | 10 | ~10 | 38.78 | <0.05 |
| $IC_{15}$ | 15 | ~15 | 49.53 | <0.05 |
| $IC_{20}$ | 20 | ~20 | 62.32 | <0.05 |

**Supplementary Table 4. Relative inhibitor concentrations in $IC_{15}$ ULDCs compared with single-agent $IC_{50}$ values**

| Inhibitor | Relative concentration in $IC_{15}$ ULDC (% of $IC_{50}$ for individual agent) |
|---|---|
| Gefitinib | 36.15 |
| Erlotinib | 23.91 |
| Afatinib | 17.16 |
| Neratinib | 18.19 |
| Osimertinib | 26.44 |

##### 1.1.3 Combination index analysis of five-drug ULDCs in PC-9 cells

To further characterize pharmacological interactions within ULDCs, combination index (CI) analysis was performed using the Chou–Talalay method [19,20].

Across the $IC_{05}$–$IC_{20}$ conditions, five-drug ULDCs exhibited CI values ranging from 0.4287 to 0.6293, consistent with synergistic pharmacological interactions according to the Chou–Talalay framework (Supplementary Table 5). The component concentrations used in the combinations and the corresponding single-agent concentrations theoretically required to produce the same observed level of growth inhibition are provided in Supplementary Table 6.

**Supplementary Table 5. Combination index (CI) analysis of five-drug ULDCs in PC-9 cells**

| Combination condition | Inhibition (%) | CI value | Interaction interpretation |
|---|---|---|---|
| $IC_{05}$ combination | 33.48 | 0.4423 | Synergistic |
| $IC_{10}$ combination | 38.78 | 0.5845 | Synergistic |
| $IC_{15}$ combination | 49.53 | 0.6293 | Synergistic |
| $IC_{20}$ combination | 62.32 | 0.4287 | Synergistic |

Note: CI values below 1 indicate synergistic interactions, CI values approximately equal to 1 indicate additive interactions, and CI values above 1 indicate antagonistic interactions according to the Chou–Talalay framework [19,20].

**Supplementary Table 6. Concentration parameters used for combination index (CI) analysis of the five-EGFR-TKI combinations in PC-9 cells**

| Groups | | Gefitinib | Erlotinib | Afatinib | Neratinib | Osimertinib |
|---|---|---|---|---|---|---|
| $CI_{05}$ | A (Combination dose) | 0.5463 | 0.4766 | 0.001438 | 0.5761 | 0.05836 |
| | B (Estimated single-agent dose producing equivalent inhibition) | 4.040 | 5.696 | 0.03216 | 7.164 | 0.5937 |
| $IC_{10}$ | A | 1.214 | 1.219 | 0.003207 | 1.134 | 0.1062 |
| | B | 5.658 | 8.682 | 0.07234 | 13.01 | 1.085 |
| $IC_{15}$ | A | 1.785 | 1.990 | 0.005906 | 1.877 | 0.1676 |
| | B | 7.421 | 12.03 | 0.1443 | 20.90 | 1.808 |

| | | | | | | |
|---|---|---|---|---|---|---|
| $IC_{20}$ | A | 2.345 | 2.830 | 0.009869 | 2.845 | 0.2460 |
| | B | 12.83 | 21.89 | 0.6537 | 50.41 | 5.448 |

Note: Rows labelled "A" represent the concentrations of the five EGFR-TKIs used in the combination treatment. Rows labelled "B" represent the theoretical concentrations of each individual EGFR-TKI required to achieve the same experimentally observed growth inhibitory effect produced by the corresponding five-EGFR-TKI combination. These concentrations were calculated by interpolation from individual concentration–response curves.

##### 1.1.4 Distributed contribution within five-drug ULDC

To assess whether collective growth inhibition depended predominantly on any single inhibitor, drop-one analyses (Supplementary Fig. 3) were performed by individually removing each component from the five-drug ULDC.

Within the ultra-low-dose range ($IC_{05}$–$IC_{20}$), individual removal of each inhibitor from the five-drug ULDCs resulted in slight reductions in overall growth inhibition, whereas no significant differences were observed among the corresponding four-drug combinations (Supplementary Fig. 3 and Supplementary Table 7). These findings support a distributed contribution model in which collective growth inhibition emerges from coordinated weak perturbations directed against a shared target rather than dependence on a single dominant inhibitor.

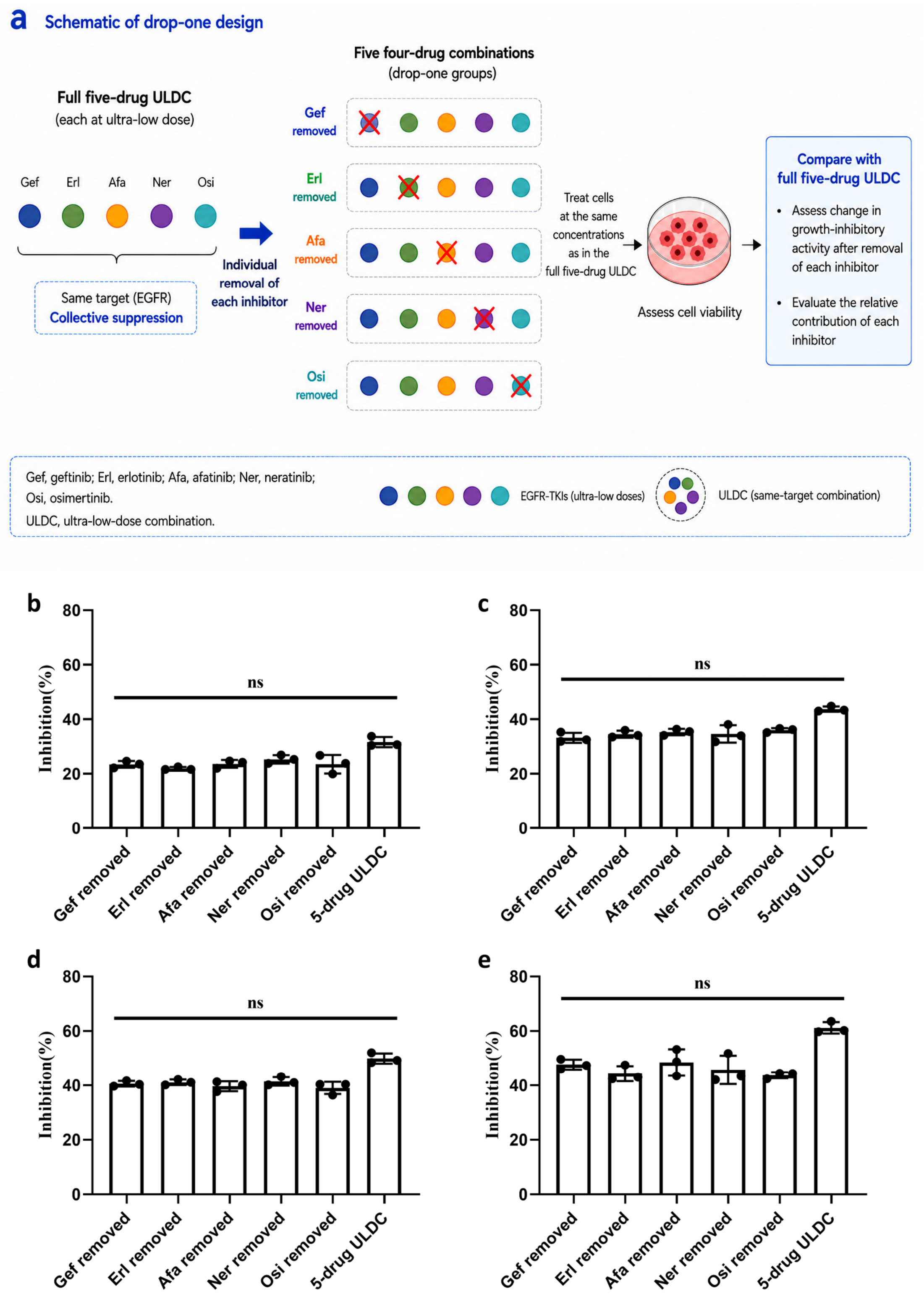


**Supplementary Fig. 3. Drop-one analysis of distributed contributions within the five-drug ULDC. a,** Schematic of the drop-one design. Each EGFR-TKI was individually removed from the complete five-drug ULDC, generating five corresponding four-drug combinations. In each drop-one condition, the remaining four inhibitors were maintained at the same concentrations as in the complete five-drug ULDC, without dose adjustment. Cell viability was compared between the four-drug and complete five-drug ULDCs. **b–e,** Drop-one analyses performed at $IC_{05}$, $IC_{10}$, $IC_{15}$, and $IC_{20}$ conditions, respectively. Each drop-one group was compared with the corresponding complete five-drug ULDC at the same concentration conditions. Data are presented as mean ± SD (n = 3); ns, not significant.

**Supplementary Table 7. Drop-one analysis of five-drug ULDCs in PC-9 cells**

| Combination condition | Full five-drug ULDC inhibition (%) | Four-drug combinations after removal of one inhibitor (%) | Statistical comparison among four-drug groups |
|---|---|---|---|
| $IC_{05}$ | 33.48 | 21.95–25.33 | ns |
| $IC_{10}$ | 38.78 | 33.21–36.01 | ns |
| $IC_{15}$ | 49.53 | 39.06–41.46 | ns |
| $IC_{20}$ | 62.32 | 43.74–48.46 | ns |

Note：ns, not significant.

##### 1.1.5 Validation across additional cellular contexts

###### 1.1.5.1 Validation in HCC827 cells

To evaluate whether collective growth inhibition extended beyond PC-9 cells, additional experiments were performed in human EGFR-mutant NSCLC HCC827 cells. Five-drug ULDCs produced substantially greater growth inhibition than individual EGFR-TKIs administered at corresponding $IC_{10}$ or $IC_{20}$ concentrations (Fig. 2c, Supplementary Fig. 4, and Supplementary Tables 8 and 9). Combination treatment achieved inhibition rates of 54.78% and 64.50% under $IC_{10}$ and $IC_{20}$ conditions, respectively, whereas individual inhibitors produced only minimal inhibition (e.g. 7.93%–11.48% under $IC_{10}$). CI analysis yielded values of 0.8716 and 0.9475 under $IC_{10}$ and $IC_{20}$ conditions, respectively, consistent with mild synergistic pharmacological interactions (Supplementary Fig. 5).

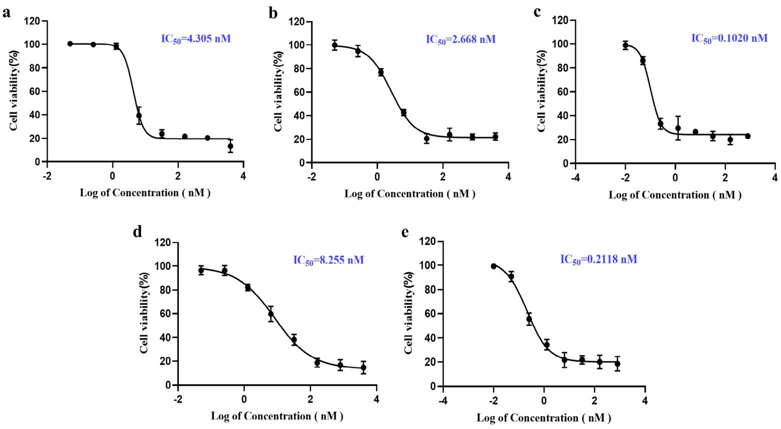


**Supplementary Fig. 4. Concentration–response curves and $IC_{50}$ values of five EGFR-TKIs for inhibition of cell viability in HCC827 cells.** **a**, Gefitinib; **b**, Erlotinib; **c**, Afatinib; **d**, Neratinib; **e**, Osimertinib. Data are presented as mean ± SD (n=3).

**Supplementary Table 8. $IC_{50}$ values of EGFR-TKIs in HCC827 and A549 cells**

| Cell line | Gefitinib | Erlotinib | Afatinib | Neratinib | Osimertinib |
|---|---|---|---|---|---|
| HCC827 (nM) | 4.305 | 2.668 | 0.1020 | 8.255 | 0.2118 |
| A549 (μM) | 24.22 | 40.04 | 2.463 | 13.36 | 1.244 |

**Supplementary Table 9. $IC_{10}$ and $IC_{20}$ concentrations used for ULDC validation experiments in HCC827 and A549 cells**

| Cell line | Condition | Gefitinib | Erlotinib | Afatinib | Neratinib | Osimertinib |
|---|---|---|---|---|---|---|
| HCC827 | $IC_{10}$ | 2.19 | 0.556 | 0.0409 | 0.548 | 0.0438 |
| HCC827 | $IC_{20}$ | 2.94 | 1.12 | 0.0621 | 1.68 | 0.0864 |
| A549 | $IC_{10}$ | 6.53 | 1.61 | 0.274 | 7.84 | 0.241 |
| A549 | $IC_{20}$ | 11.0 | 4.88 | 0.681 | 9.86 | 0.474 |

Note：Concentrations for HCC827 cells are expressed in nM, whereas concentrations for A549 cells are expressed in μM.

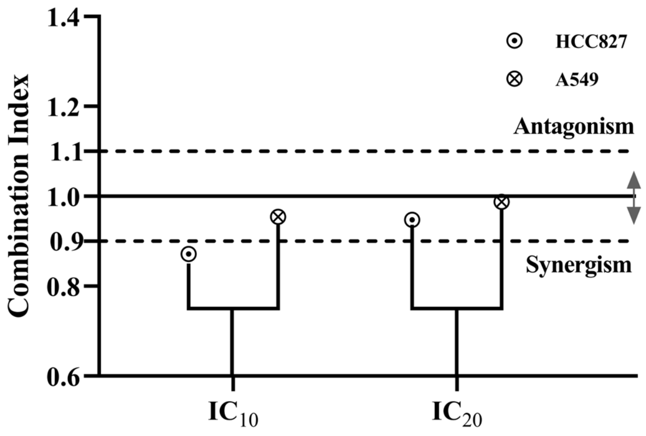


**Supplementary Fig. 5. Combination index (CI) values of five-EGFR-TKI combinations in HCC827 and A549 cell lines.** CI values below 1 indicate synergistic interactions, CI values approximately equal to 1 indicate additive interactions, and CI values above 1 indicate antagonistic interactions according to the Chou–Talalay framework. The horizontal dashed lines indicate the corresponding CI levels on the y-axis.

##### 1.1.5.2 Collective growth inhibition in non-EGFR-driven cellular contexts

To explore whether aspects of the ULDC phenomenon extended beyond highly EGFR-dependent systems, similar experiments were performed in KRAS-mutant human NSCLC A549 cells. Although A549 cells were substantially less sensitive to EGFR-TKIs than EGFR-mutant models, five-drug ULDCs still produced measurable collective growth-inhibitory effects, achieving inhibition rates of 52.47% and 71.82% under $IC_{10}$ and $IC_{20}$ conditions, respectively (Fig. 2d, Supplementary Fig. 6, and Supplementary Tables 8 and 9). CI values in A549 cells yielded values of 0.9546 and 0.9876 under $IC_{10}$ and $IC_{20}$ conditions, and were closer to 1, consistent with near-additive pharmacological interactions (Supplementary Fig. 5). Together, these findings suggest that five-drug ULDCs can retain measurable collective growth-inhibitory activity beyond highly EGFR-dependent models, although the mechanistic basis and biological relevance in non-EGFR-driven systems remain to be established.

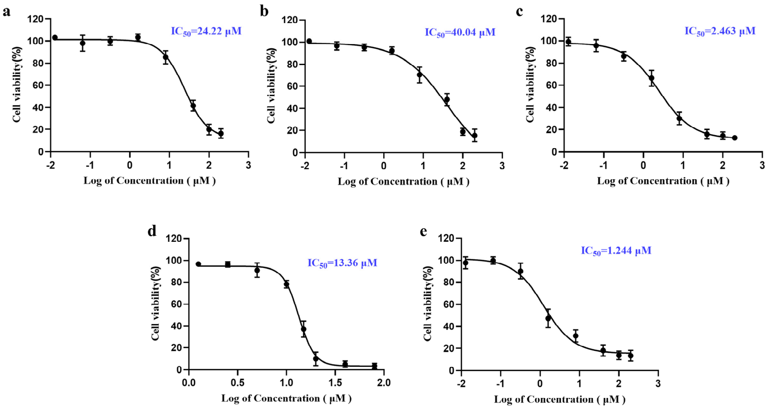


**Supplementary Fig. 6. Concentration–response curves and $IC_{50}$ values for five EGFR-TKIs for inhibition of cell viability in A549 cells.** **a**, Gefitinib; **b**, Erlotinib; **c**, Afatinib; **d**, Neratinib; **e**, Osimertinib. Data are presented as mean ± SD (n=3).

#### 1.2 Scaling of collective growth inhibition with increasing same-target inhibitor diversity

##### 1.2.1 Expanded EGFR-TKI panel

To investigate how collective growth inhibition scaled with increasing same-target inhibitor diversity, the number of distinct EGFR-TKIs included within the ULDC architecture was expanded from five to eight, ten, and fourteen inhibitors. Additional EGFR-TKIs included dacomitinib, olmutinib, icotinib, almonertinib, lapatinib, furmonertinib, rezivertinib, befotertinib, and oritinib. Concentration–response characterization of the additional EGFR-TKIs in PC-9 cells is shown in Supplementary Figs. 7–9, and corresponding $IC_{50}$ values are summarized in Supplementary Table 10. $IC_{05}$ and $IC_{10}$ concentrations used for higher-order ULDC experiments are summarized in Supplementary Table 11.

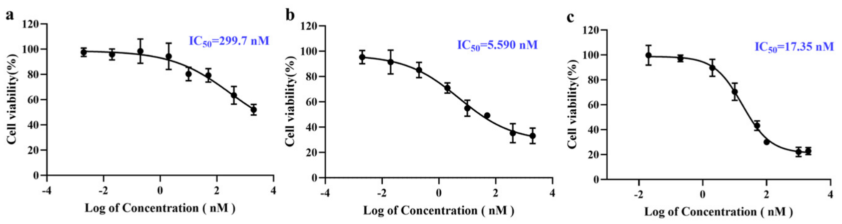


**Supplementary Fig. 7. Concentration–response curves for three additional EGFR-TKIs used to expand the five-drug ULDC to an eight-drug combination in PC-9 cells.**

**a**, Dacomitinib; **b**, Olmutinib; **c**, Icotinib. Data are presented as mean ± SD (n=3).

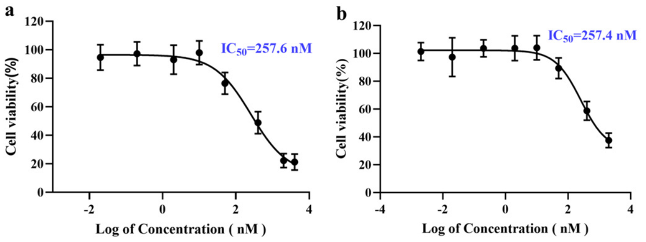


**Supplementary Fig. 8. Concentration–response curves for two additional EGFR-TKIs used to expand the eight-drug ULDC to a ten-drug combination in PC-9 cells.** **a**, Almonertinib; **b**, Lapatinib. Data are presented as mean ± SD (n=3).

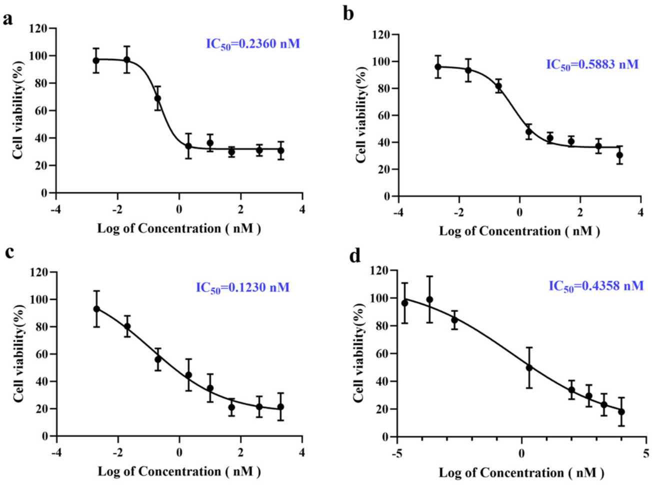


**Supplementary Fig. 9. Concentration–response curves for four additional EGFR-TKIs used to expand the ten-drug ULDC to a 14-drug combination in PC-9 cells.** **a**, Furmonertinib; **b**, Rezivertinib; **c**, Befotertinib; **d**, Oritinib. Data are presented as mean ± SD (n=3).

**Supplementary Table 10. $IC_{50}$ values of additional EGFR-TKIs in PC-9 cells**

| Inhibitor | $IC_{50}$ |
|---|---|
| Dacomitinib | 299.7 nM |
| Olmutinib | 5.590 nM |
| Icotinib | 17.35 nM |
| Almonertinib | 257.6 nM |
| Lapatinib | 257.4 nM |

| Inhibitor | $IC_{50}$ |
|---|---|
| Furmonertinib | 0.2360 nM |
| Rezivertinib | 0.5883 nM |
| Befotertinib | 0.1230 nM |
| Oritinib | 0.4358 nM |

**Supplementary Table 11.  $IC_{05}$ and $IC_{10}$ concentrations of individual EGFR-TKIs for constructing higher-order ULDCs in PC-9 cells**

| No. | Drug | $IC_{05}$ (nM) | $IC_{10}$ (nM) |
|---|---|---|---|
| 1 | Gefitinib | 0.5463 | 1.214 |
| 2 | Erlotinib | 0.4766 | 1.219 |
| 3 | Afatinib | 0.0014 | 0.0032 |
| 4 | Neratinib | 0.5761 | 1.134 |
| 5 | Osimertinib | 0.0584 | 0.1062 |
| 6 | Dacomitinib | 0.3483 | 3.224 |
| 7 | Olmutinib | 0.0025 | 0.0463 |
| 8 | Icotinib | 0.8478 | 2.144 |
| 9 | Almonertinib | 2.627 | 15.08 |
| 10 | Lapatinib | 32.58 | 57.72 |
| 11 | Furmonertinib | 0.0309 | 0.0654 |
| 12 | Rezivertinib | 0.0110 | 0.0630 |
| 13 | Befotertinib | 0.0015 | 0.0038 |
| 14 | Oritinib | 0.0010 | 0.0071 |

Note：$IC_{05}$ and $IC_{10}$ concentrations for each EGFR-TKI were derived by interpolation from the corresponding concentration–response curves in PC-9 cells. Five-, eight-, ten-, and fourteen-drug ULDCs, including inhibitors 1–5, 1–8, 1–10, and 1–14, respectively, represent increasing same-target inhibitor diversity.

##### 1.2.2 Increasing same-target diversity progressively enhances collective growth inhibition

To determine whether increasing same-target inhibitor diversity further enhanced collective growth inhibition, the number of EGFR-TKIs included in ULDCs was progressively expanded from five to eight, ten, and fourteen inhibitors. Importantly, each component inhibitor remained at its predefined ultra-low concentration, such that expansion of the ULDCs increased the diversity of weak perturbations directed against the same target rather than increasing the dose of any individual inhibitor.

In the inhibitor-diversity scaling experiments reported here, increasing same-target inhibitor diversity progressively increased growth inhibition in PC-9 cells under $IC_{05}$ conditions (Supplementary Fig. 10). The five-drug ULDC produced 31.37% growth inhibition, whereas the eight-, ten-, and fourteen-drug ULDCs produced 40.41%, 47.07%, and 54.61% inhibition, respectively. A similar scaling pattern was observed under $IC_{10}$ conditions (Supplementary Fig. 11), with growth inhibition increasing from 41.56% for the five-drug ULDC to 54.96%, 64.99%, and 81.15% for the eight-, ten-, and fourteen-drug ULDCs, respectively. Direct comparison across increasing inhibitor numbers demonstrated a progressive increase in collective growth inhibition with increasing same-target inhibitor diversity (Supplementary Fig. 12).

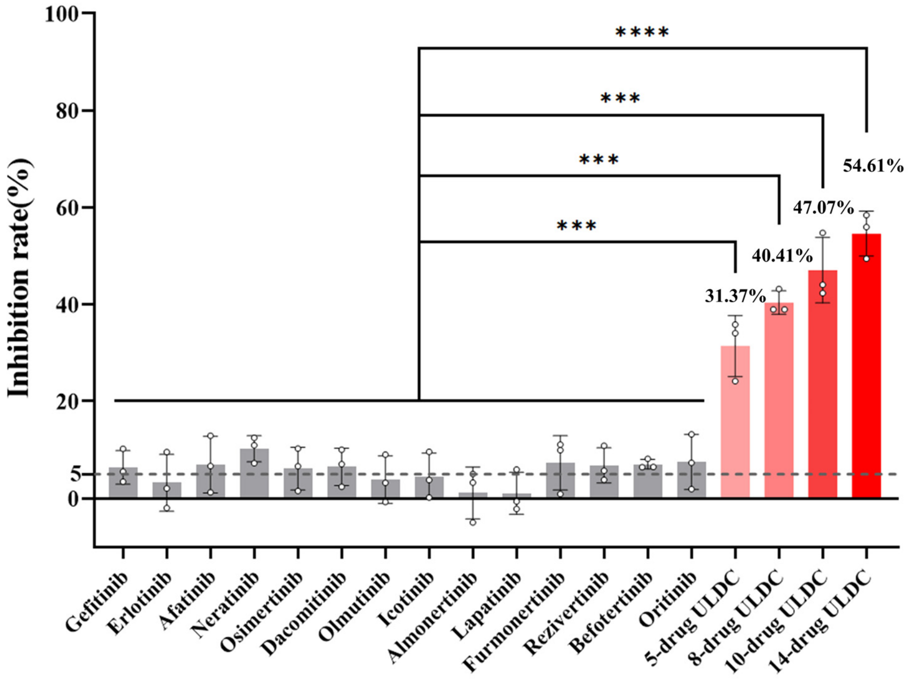


**Supplementary Fig. 10. Increasing same-target diversity progressively enhances collective growth inhibition under $IC_{05}$ conditions.** Increasing the number of EGFR-TKIs included in the

ULDCs progressively enhanced inhibition of PC-9 cell growth under $IC_{05}$ conditions. All higher-order ULDCs (8-drug, 10-drug, and 14-drug ULDCs) produced substantially greater inhibition than the corresponding single-agent at ultra-low-dose conditions. The horizontal dashed lines indicate the corresponding growth inhibition levels on the y-axis. Statistical comparisons were performed using one-way ANOVA with the corresponding single-agent group as the comparator; ****$P < 0.0001$, ***$P < 0.001$. Data are presented as mean ± SD (n = 3).

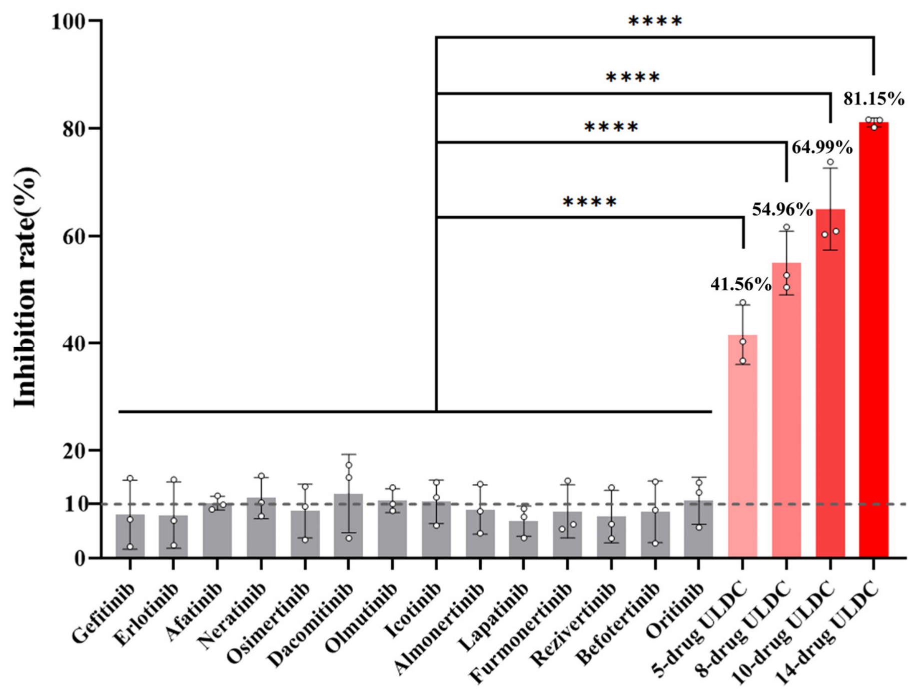


**Supplementary Fig. 11. Increasing same-target diversity progressively enhances collective growth inhibition under $IC_{10}$ conditions.** Progressive enhancement of collective growth inhibition was observed under $IC_{10}$ conditions as inhibitor number increased from five to fourteen EGFR-TKIs. The horizontal dashed lines indicate the corresponding growth inhibition levels on the y-axis. Statistical comparisons were performed using one-way ANOVA with the corresponding single-agent group as the comparator; ****$P < 0.0001$. Data are presented as mean ± SD (n = 3).

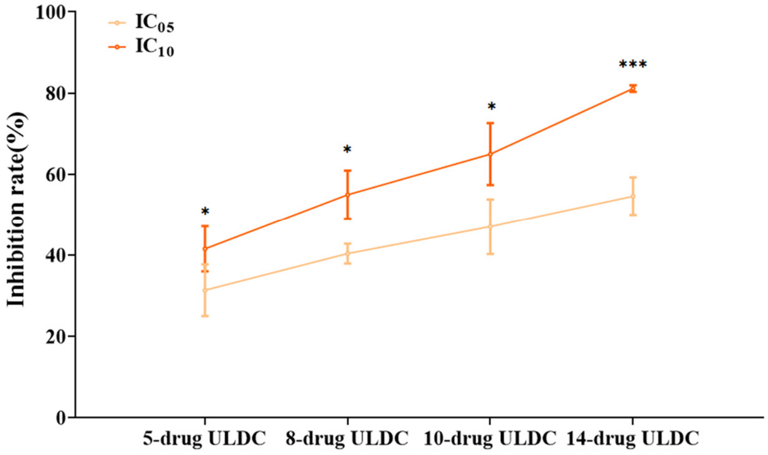


**Supplementary Fig. 12. Scaling relationship between same-target diversity and collective growth inhibition.** Collective growth inhibition progressively increased with increasing same-target diversity under both $IC_{05}$ and $IC_{10}$ conditions, revealing a scaling relationship between inhibitor diversity and collective growth inhibition. Statistical comparisons were performed between the $IC_{05}$ and $IC_{10}$ conditions at each inhibitor number; ***P < 0.001, *P < 0.05. Data are presented as mean ± SD (n = 3).

##### 1.2.3. Combination index analysis across increasing inhibitor numbers

Combination index (CI) analysis demonstrated progressively decreasing CI values with increasing same-target inhibitor diversity under both $IC_{05}$ and $IC_{10}$ conditions (Supplementary Table 12), indicating stronger synergistic pharmacological interactions among higher-order ULDCs according to the Chou-Talalay framework.

**Supplementary Table 12. CI analysis across increasing same-target diversity conditions**

| Condition | 5-drug | 8-drug | 10-drug | 14-drug |
|---|---|---|---|---|
| $IC_{05}$ | 0.4423 | 0.3575 | 0.3228 | 0.2698 |
| $IC_{10}$ | 0.5845 | 0.3688 | 0.2238 | 0.0402 |

Note: Lower CI values indicate stronger pharmacological interactions according to the Chou–Talalay framework.

#### Supplementary Note 2. Resistance emergence and resistant-state properties under ULDCs

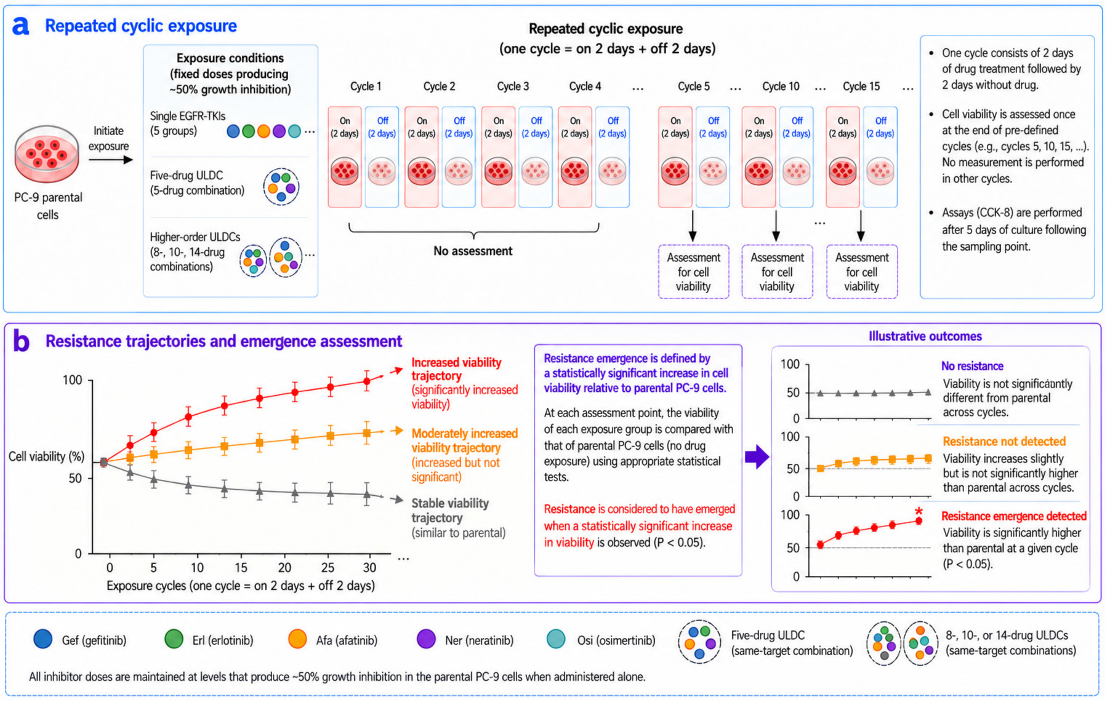


**Supplementary Fig. 13. Repeated drug exposure framework and assessment of resistance emergence.** **a**, Schematic of the repeated cyclic exposure protocol. Parental PC-9 cells were assigned to individual EGFR-TKI groups, the five-drug ULDC, or higher-order ULDCs comprising 8, 10, or 14 inhibitors. Treatment conditions were selected to produce approximately matched levels of growth inhibition (~50%) in parental PC-9 cells. Each exposure cycle consisted of 2 days of drug treatment followed by 2 days without drug. Cell viability was assessed only at the end of pre-defined exposure cycles (cycles 5, 10, 15, and subsequent indicated time points) by CCK-8 assay after 5 days of culture following sampling. No measurements were performed during the intervening cycles. **b**, Schematic representation of longitudinal viability trajectories and assessment of resistance emergence. At each pre-defined assessment point, cell viability of repeatedly exposed cells was compared with the viability of the parental PC-9 cells under identical assay conditions to evaluate the emergence of resistance. Resistance was considered to have emerged when cell viability was significantly higher than that of the parental PC-9 cells ($P < 0.05$). Illustrative trajectories show stable viability, moderately increased viability without detected resistance, and increased viability associated with detected resistance emergence during repeated cyclic exposure.

#### 2.1 Resistance emergence under repeated drug exposure

##### 2.1.1 Approximately matched growth-inhibition conditions for repeated cyclic drug exposure

To compare resistance emergence under approximately matched levels of growth inhibition, repeated cyclic drug exposure experiments (Supplementary Fig. 13) were performed using treatment conditions producing approximately 50% growth inhibition in PC-9 cells. Single-agent groups were treated at concentrations producing approximately 50% growth inhibition, whereas ULDCs were assembled from substantially lower concentrations of the constituent EGFR-TKIs.

For the five-drug ULDC, each EGFR-TKI was administered at approximately its $IC_{15}$ concentration, yielding a level of growth inhibition comparable to that of the single-agent groups (Supplementary Table 13).

For the 8-, 10-, and 14-drug ULDCs, combination concentrations were further adjusted to achieve approximately matched levels of growth inhibition (~50%) before repeated cyclic drug exposure experiments were initiated (Supplementary Table 14 and Supplementary Fig. 14).

These approximately matched growth-inhibition conditions were used to minimize differences attributable simply to inhibitory strength when comparing resistance emergence across treatment conditions.

**Supplementary Table 13. Concentrations used in repeated cyclic drug exposure experiments for five EGFR-TKIs and the five-drug ULDC**

| Group | Treatment condition | Approximate growth inhibition |
|---|---|---|
| Gefitinib | $IC_{50}$ | 50% |
| Erlotinib | $IC_{50}$ | 50% |
| Afatinib | $IC_{50}$ | 50% |
| Neratinib | $IC_{50}$ | 50% |
| Osimertinib | $IC_{50}$ | 50% |
| Five-drug ULDC | Each agent at approximately $IC_{15}$ | 50% |

Note: Single-agent groups and the five-drug ULDC group were treated at concentrations producing approximately 50% growth inhibition in PC-9 cells. The five-drug ULDC consisted of five EGFR-

TKIs administered simultaneously at individually weak inhibitory concentrations (~$IC_{15}$ each), while maintaining comparable overall inhibition.

**Supplementary Table 14. Optimization of matched-inhibition conditions for higher-order ULDCs**

| Group | Initial combination condition | Final concentration used for repeated cyclic drug exposure | Approximate growth inhibition |
|---|---|---|---|
| 8-drug ULDC | Modified $IC_{10}$-based combination | 95% of $IC_{10}$ combination | 50% |
| 10-drug ULDC | Modified $IC_{05}$-based combination | 105% of $IC_{05}$ combination | 50% |
| 14-drug ULDC | Modified $IC_{05}$-based combination | 90% of $IC_{05}$ combination | 50% |

Note: Combination concentrations were adjusted to achieve approximately matched growth inhibition prior to repeated cyclic drug exposure experiments.

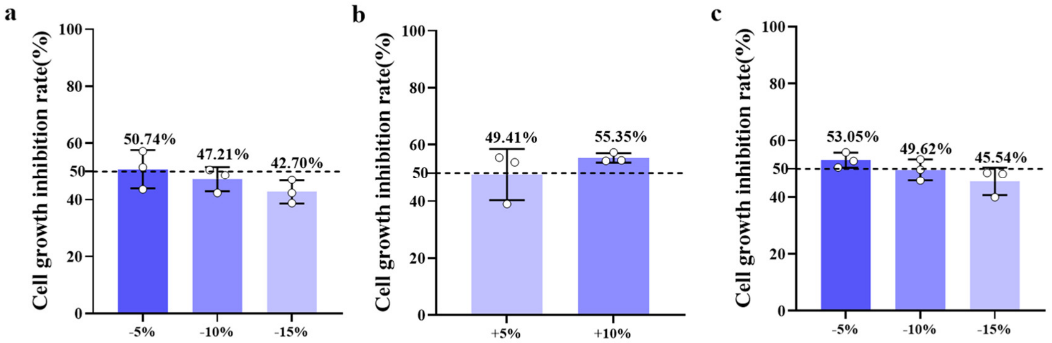


**Supplementary Fig. 14. Optimization of concentrations for higher-order ULDCs under matched levels of growth inhibition in PC-9 cells.** **a**, Concentration adjustment for the 8-drug ULDC. The concentrations of the eight constituent EGFR-TKIs in the $IC_{10}$-based combination were proportionally reduced by 5%, 10%, or 15%. **b**, Concentration adjustment for the 10-drug ULDC. The concentrations of the ten constituent EGFR-TKIs in the $IC_{05}$-based combination were proportionally increased by 5% or 10%. **c**, Concentration adjustment for the 14-drug ULDC. The concentrations of the fourteen constituent EGFR-TKIs in the $IC_{05}$-based combination were proportionally reduced by 5%, 10%, or 15%. Combination concentrations were adjusted to achieve approximately matched levels of growth inhibition (~50%) prior to repeated cyclic drug exposure experiments. The horizontal dashed lines indicate the corresponding growth inhibition rate levels on the y-axis. Data are presented as mean ± SD (n=3).

##### 2.1.2 Five-drug ULDC delays resistance emergence

Under approximately matched levels of growth inhibition, repeated cyclic drug exposure revealed distinct patterns of resistance emergence between single-agent treatment and the five-drug ULDC. All single-agent EGFR-TKI groups progressively developed resistance during prolonged exposure. Increased cell viability became detectable after repeated drug exposure cycles, although the timing (between 5 and 15 exposure cycles, corresponding to approximately 20 and 60 days) differed among individual inhibitors (Supplementary Fig. 15).

By contrast, detectable resistance emergence was substantially delayed in the five-drug ULDC group. No significant resistance emergence was detected during the early and intermediate stages of repeated drug exposure cycles (Supplementary Fig. 15). However, a gradual upward trend in cell viability became apparent over extended treatment, and significant resistance emergence was eventually observed after prolonged exposure (30 exposure cycles, corresponding to approximately 120 days) (Fig. 3). Consistent with this delay, cell viability was already significantly elevated in all five single-agent groups after 15 exposure cycles, whereas no significant increase was detected in the five-drug ULDC group through 25 cycles; a significant increase in cell viability first became apparent after 30 cycles and remained detectable at subsequent assessment points (Supplementary Table 15).

These findings indicate that the five-drug ULDC markedly delayed resistance emergence relative to single-agent treatments under approximately matched levels of growth inhibition, although it did not prevent resistance emergence during prolonged observation.

**Supplementary Table 15. Cell viability at different stages of resistance emergence under single-agent and five-drug ULDC treatment**

| Group | Time point of assessment | Cell viability (%) | P value vs parental PC-9 cells |
|---|---|---|---|
| Gefitinib | 15 exposure cycles (corresponding to approximately 60 days) | 84.61 | ***P<0.001 |
| Erlotinib | | 85.08 | ***P<0.001 |
| Afatinib | | 72.42 | *P<0.05 |
| Neratinib | | 75.38 | **P<0.01 |

| | | | |
|---|---|---|---|
| Osimertinib | | 67.55 | **P<0.01 |
| 5-drug ULDC | 15 exposure cycles | 55.12 | ns |
| | 20 exposure cycles | 55.85 | ns |
| | 25 exposure cycles | 57.72 | ns |
| | 30 exposure cycles | 64.11 | **P<0.01 |
| | 35 exposure cycles | 63.27 | **P<0.01 |
| | 40 exposure cycles | 70.68 | **P<0.01 |

Note: Statistical comparisons were performed relative to parental PC-9 cells cultured in drug-free complete medium. ***P < 0.001, **P < 0.01 and *P < 0.05; ns, not significant. Data are presented as mean ± SD. (n = 3).

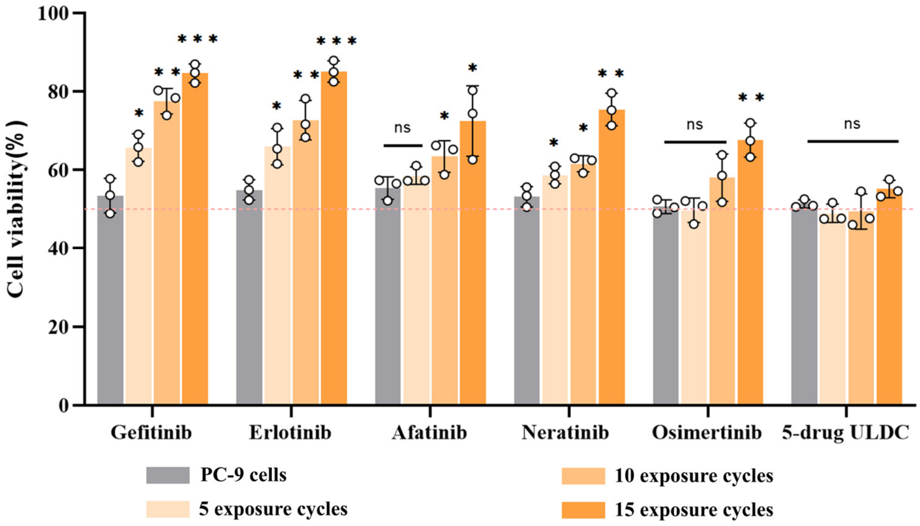


**Supplementary Fig. 15. Repeated cyclic drug exposure experiments with single-agent EGFR-TKIs and the five-drug ULDC.** Representative intermediate assessment points from repeated cyclic drug exposure experiments comparing single-agent EGFR-TKIs and the five-drug ULDC in PC-9 cells are shown. Single-agent groups progressively developed resistance during repeated cyclic drug exposure. The five-drug ULDC substantially delayed resistance emergence but eventually showed increased cell viability during prolonged exposure. All treatment conditions were selected to produce approximately matched levels of growth inhibition (~50%). Repeated cyclic drug exposure was performed for 5, 10, or 15 cycles, with each cycle lasting 4 days. The orange dashed line indicates 50% cell viability. Compared with untreated control PC-9 cells cultured in complete medium without drugs, ***P ＜ 0.001, **P ＜ 0.01, *P ＜ 0.05; ns, not significant. Data are presented as mean ± SD (n = 3).

##### 2.1.3 Increasing same-target inhibitor diversity further delays detectable resistance emergence

To determine whether increasing same-target inhibitor diversity further influenced the timing of resistance emergence, repeated cyclic drug exposure experiments were extended to ULDCs containing 8, 10, and 14 EGFR-TKIs. Whereas the five-drug ULDC eventually showed evidence of resistance emergence during prolonged cyclic drug exposure, no detectable resistance emergence was observed in the 8-, 10-, and 14-drug ULDC groups throughout the observation window, including after extended repeated drug exposure cycles (~160 days) (Fig. 3b).

Although higher-order ULDCs contained a greater number of drug components, all repeated cyclic exposure experiments were initiated under conditions producing approximately matched levels of growth inhibition. Thus, the observed differences in resistance emergence cannot be attributed simply to differences in initial inhibitory strength, although the contribution of differences in total drug exposure across treatment conditions cannot be excluded. Together, these findings indicate that increasing same-target inhibitor diversity was associated with further delay in detectable resistance emergence, with no detectable resistance observed in the 8-, 10-, or 14-drug ULDC groups within the observation window. This diversity-dependent pattern is consistent with a model in which increasing same-target inhibitor diversity may impose greater adaptive constraint during resistance evolution, although the underlying adaptive trajectories were not directly measured.

##### 2.1.4 Combination index analysis across higher-order ULDCs

Combination index (CI) analysis showed lower CI values in the 8-, 10-, and 14-drug ULDCs than in the five-drug ULDC (Supplementary Table 16), consistent with stronger synergistic pharmacological interactions according to the Chou–Talalay framework. However, because all repeated cyclic drug exposure experiments were performed under approximately matched levels of growth inhibition, the differences in resistance emergence could not be explained solely by increased pharmacological synergy.

**Supplementary Table 16. Combination index analysis of higher-order ULDCs**

| Group | CI value |
|---|---|
| 5-drug ULDC | 0.6293 |

| Group | CI value |
|---|---|
| 8-drug ULDC | 0.4799 |
| 10-drug ULDC | 0.2790 |
| 14-drug ULDC | 0.3116 |

Note: Lower CI values indicate stronger synergistic pharmacological interactions according to the Chou–Talalay framework.

#### 2.2 Five-drug ULDC activity in resistant PC-9/GR cells

To evaluate whether five-drug ULDCs retained inhibitory activity in a resistant cellular context, experiments were performed using gefitinib-resistant PC-9/GR cells.

Resistance to gefitinib markedly reduced the inhibitory activity of most individual EGFR-TKIs when administered at the ultra-low concentrations defined in parental PC-9 cells (Supplementary Table 2). At the $IC_{05}$–$IC_{20}$ concentrations, individual inhibitors generally produced minimal or no measurable growth inhibition in PC-9/GR cells, with only afatinib and particularly osimertinib retaining partial activity, indicating incomplete cross-resistance across the tested EGFR-TKIs.

Despite the markedly reduced activity of individual inhibitors, five-drug ULDCs consistently produced substantially greater growth inhibition than any corresponding single-agent treatment across all ultra-low-dose combinations (Fig. 4a and Supplementary Table17). Importantly, this enhanced inhibitory activity was achieved using the same ultra-low-dose concentrations defined in parental PC-9 cells, rather than concentrations individually optimized for resistant PC-9/GR cells. Thus, five-drug ULDCs retained substantial collective growth-inhibitory activity despite substantial loss of sensitivity to individual EGFR-TKIs in resistant cells.

Although the overall inhibitory effects of the five-drug ULDCs were reduced in PC-9/GR cells compared with parental PC-9 cells (Supplementary Table 17 and Supplementary Fig. 16), substantial collective growth-inhibitory activity remained evident across all ultra-low-dose conditions.

Combination index analysis further showed that pharmacological interactions remained within the synergistic range according to the Chou–Talalay framework in the resistant cellular context, with all ultra-low-dose combinations exhibiting CI values below 1, namely 0.4225, 0.2841, 0.2745, and 0.1620 for $IC_{05}$ Combo, $IC_{10}$ Combo, $IC_{15}$ Combo, and $IC_{20}$ Combo, respectively (Supplementary Table 18 and Supplementary Fig. 17).

Together, these findings demonstrate that the collective growth-inhibitory activity of the five-drug ULDC is retained in a gefitinib-resistant cellular context despite markedly reduced sensitivity to individual EGFR-TKIs.

**Supplementary Table 17. Inhibition rates (%) of the five-EGFR-TKI combination at different ultra-low-dose conditions in PC-9 and PC-9/GR cells (n = 3).**

| Groups | PC-9 cells | PC-9/GR cells | *P* value |
|---|---|---|---|
| $IC_{05}$ Combo | 28.85±4.372 | 13.88±2.931 | 0.0048 |
| $IC_{10}$ Combo | 38.82±4.227 | 24.40±2.153 | 0.0026 |
| $IC_{15}$ Combo | 51.39±4.211 | 32.10±2.822 | 0.0015 |
| $IC_{20}$ Combo | 62.32±8.187 | 46.37±5.740 | 0.0205 |

Note: The detailed concentrations of individual EGFR-TKIs in $IC_{05}$ Combo to $IC_{20}$ Combo are provided in Supplementary Table 2. Comparisons of inhibition rates between PC-9 and PC-9/GR cells were performed, and P < 0.05 was considered statistically significant.

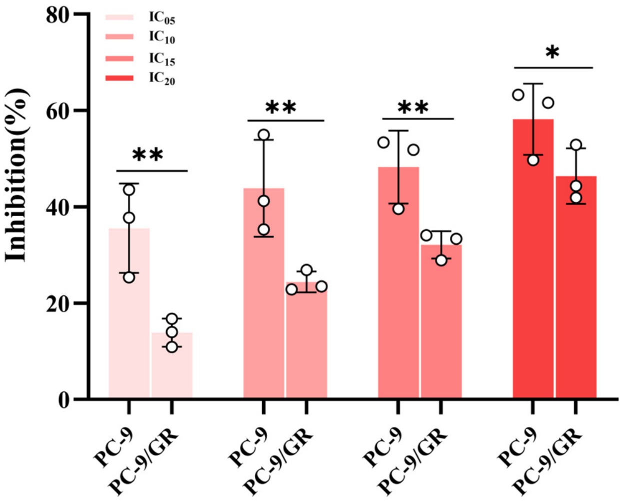


**Supplementary Fig. 16. Comparison of the growth-inhibitory effects of the five EGFR-TKI combinations at ultra-low-dose concentrations in parental PC-9 and gefitinib-resistant PC-9/GR cells.** Growth inhibition (%) was compared between PC-9 and PC-9/GR cells treated with the five-EGFR-TKI combination at four ultra-low-dose concentration levels ($IC_{05}$, $IC_{10}$, $IC_{15}$, and $IC_{20}$). Bars represent mean ± SD (n = 3), with individual data points shown. Statistical comparisons between PC-9 and PC-9/GR cells at each concentration level were performed using two-tailed independent-samples t-tests. *P < 0.05; **P < 0.01.

**Supplementary Table 18. Concentrations used for combination index (CI) analysis of the five-EGFR-TKI combinations in PC-9/GR cells （nM）**

| Combination condition | | Gefitinib | Erlotinib | Afatinib | Neratinib | Osimertinib |
|---|---|---|---|---|---|---|
| $IC_{05}$ Combo | A | 0.5463 | 0.4766 | 0.001438 | 0.5761 | 0.05836 |
| | B | 825.8 | 54.54 | 0.01877 | 22.31 | 0.1878 |
| $IC_{10}$ Combo | A | 1.214 | 1.219 | 0.003207 | 1.134 | 0.1062 |
| | B | 1639.5 | 216.1 | 0.1212 | 62.22 | 0.4558 |
| $IC_{15}$ Combo | A | 1.785 | 1.990 | 0.005906 | 1.877 | 0.1676 |
| | B | 2218.2 | 416.9 | 0.3513 | 98.77 | 0.7184 |
| $IC_{20}$ Combo | A | 2.345 | 2.830 | 0.009869 | 2.845 | 0.2460 |
| | B | 3611 | 1271 | 2.874 | 209.1 | 1.731 |

Note: Rows labelled "A" represent the concentrations of the five EGFR-TKIs used in the combination treatment. Rows labelled "B" represent the theoretical concentrations of each individual EGFR-TKI required to achieve the same experimentally observed growth inhibitory effect produced by the corresponding five-EGFR-TKI combination. The detailed concentrations of individual EGFR-TKIs in $IC_{05}$ Combo to $IC_{20}$ Combo are provided in Supplementary Table 2.

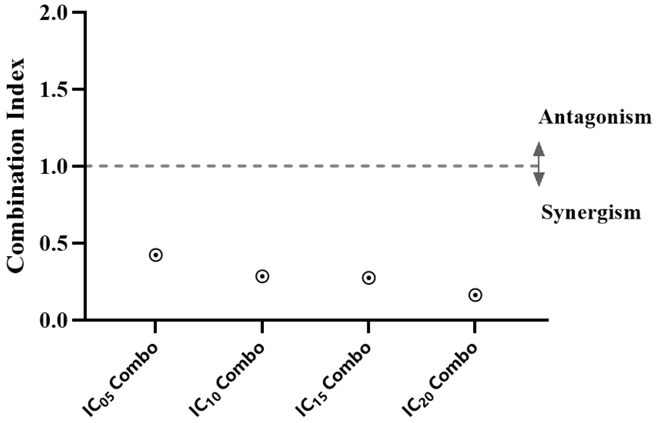


**Supplementary Fig. 17. Combination index (CI) values of the five-EGFR-TKI combination across $IC_{05}$ Combo to $IC_{20}$ Combo conditions in PC-9/GR cells.** $IC_{05}$ Combo, $IC_{10}$ Combo, $IC_{15}$ Combo, or $IC_{20}$ Combo represent four combinations generated by combining the five EGFR-TKIs at their respective $IC_{05}$, $IC_{10}$, $IC_{15}$, or $IC_{20}$ concentrations determined in PC-9 cells. CI values were interpreted according to the Chou–Talalay framework, with values below, approximately equal to, or above 1 corresponding to synergistic, additive, or antagonistic interactions, respectively. The corresponding CI values were 0.4225, 0.2841, 0.2745, and 0.1620 for $IC_{05}$ Combo, $IC_{10}$ Combo, $IC_{15}$ Combo, and $IC_{20}$ Combo, respectively.

#### 2.3 Reversibility of resistance states

Having established that the five-drug ULDC retains collective growth-inhibitory activity in a gefitinib-resistant cellular context, we next asked whether the resistant states generated under five-drug ULDC conditions differ in their stability from those induced by single-agent treatment.

To assess the stability and reversibility of resistance induced by prolonged drug selection, resistant PC-9 cells generated by either repeated single-agent EGFR-TKI treatment or the five-drug ULDC were cultured in drug-free complete medium and subsequently reassessed for drug sensitivity. Single-agent resistant cells were generated after 15 repeated drug exposure cycles with gefitinib, erlotinib, afatinib, neratinib, or osimertinib, whereas ULDC-resistant cells were generated after 40 repeated drug exposure cycles. Cells were maintained in drug-free medium for five consecutive recovery periods (two cell passages per period), and drug sensitivity was reassessed after each recovery period using the same treatment conditions employed during the original resistance-selection experiments. The treatment concentrations corresponded to those producing approximately 50% growth inhibition in parental PC-9 cells, namely the $IC_{50}$ concentration for each individual EGFR-TKI and the matched growth-inhibition condition for the five-drug ULDC.

Following withdrawal of drug selection, resistance generated under five-drug ULDC conditions showed substantially greater reversibility than resistance induced by individual EGFR-TKI treatment. Cell viability of ULDC-resistant cells decreased significantly after the first recovery period ($P < 0.05$) and continued to decline during subsequent drug-free passages, approaching the parental sensitivity level after three recovery periods. In contrast, although drug sensitivity gradually recovered in all single-agent groups, none returned to the parental response level even after five recovery periods (Fig. 4b). Because resistance emergence was substantially delayed under five-drug ULDC treatment, ULDC-resistant populations had not yet emerged at the time point when single-agent-resistant populations were already established. Consistent with this difference in resistance kinetics, cell viability remained substantially lower in the five-drug ULDC group than in all five single-agent groups after 15 exposure cycles and increased only with further prolonged ULDC exposure (Supplementary Table 15). Therefore, ULDC-resistant cells analysed in these experiments were obtained after 40 drug exposure cycles, whereas single-agent-resistant cells were obtained after 15 cycles. Notably, despite undergoing substantially more prolonged drug selection, ULDC-resistant cells exhibited markedly faster restoration of drug sensitivity following withdrawal of drug selection, consistent with a less stable resistant state than that observed after single-agent treatment. In this context, the five-drug ULDC-selected resistant population exhibited a cell viability of 70.68% before drug-free recovery, comparable to those of the afatinib- and osimertinib-selected resistant populations (72.42% and 67.55%, respectively). Despite these similar initial viability levels, the ULDC-selected population recovered drug sensitivity more rapidly than either single-agent-selected population (Fig. 4b and Supplementary Table 15).

Together, these findings indicate that resistance generated under five-drug ULDC conditions was less stable than resistance induced by prolonged single-agent treatment. Although five-drug ULDC treatment substantially delayed resistance emergence during repeated drug exposure, the resistant states that eventually arose exhibited substantially greater reversibility following withdrawal of drug selection. These findings support the possibility that therapeutic perturbation architecture may influence not only the timing of resistance emergence but also the stability of the resistant states that subsequently arise.

#### Supplementary Note 3. Transcriptomic profiling reveals distinct biological responses induced by the five-drug ULDC

To further characterize the transcriptional responses associated with five-drug ULDC treatment, we performed transcriptomic profiling of PC-9 cells treated with individual EGFR-TKIs or the five-drug ULDC under matched growth-inhibitory conditions (~50%). Three biological replicates were analysed for each treatment group.

##### 3.1 The five-drug ULDC induces a distinct transcriptomic profile compared with individual EGFR-TKIs

Principal component analysis (PCA) showed clear separation between the transcriptomic profiles induced by the five-drug ULDC and those induced by individual EGFR-TKIs. Biological replicates within each treatment group clustered closely, indicating good reproducibility of the RNA-seq datasets. This clustering pattern indicates that the transcriptional response associated with the five-drug ULDC differs from those associated with individual inhibitors under comparable inhibitory conditions (Supplementary Fig. 18).

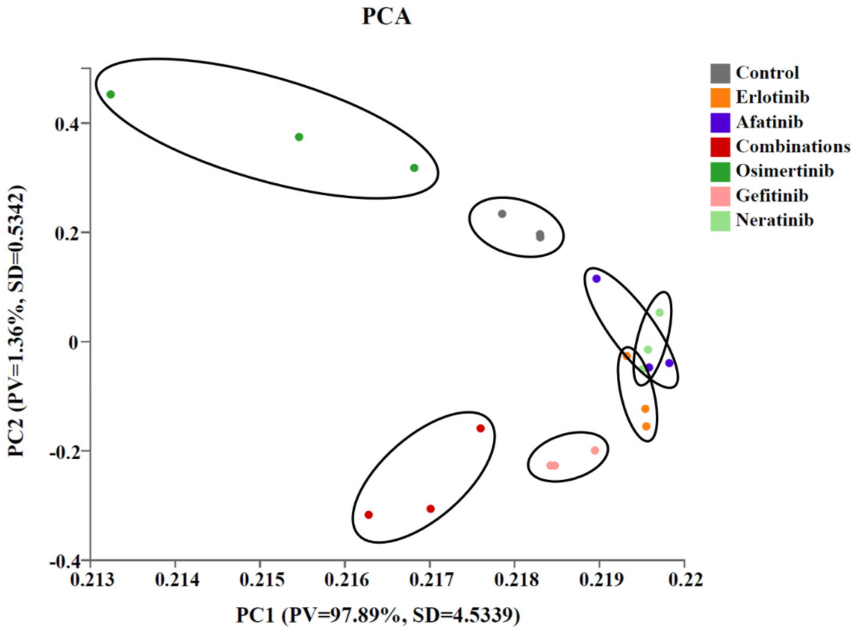


**Supplementary Fig. 18. Principal component analysis of RNA-seq samples from PC-9 cells treated with individual EGFR-TKIs or the five-drug ULDC.** Biological replicates clustered closely within each treatment group, whereas ULDC-treated samples exhibited distinct separation from individual inhibitor-treated groups. Control, PC-9 cells in drug-free complete medium; Combination, five-drug ULDC group.

#### 3.2 The five-drug ULDC produces broader transcriptional changes than individual inhibitors

Differential expression analysis revealed that individual EGFR-TKIs altered the expression of hundreds of genes, whereas the five-drug ULDC induced substantially broader transcriptional changes, including 827 upregulated and 733 downregulated differentially expressed genes (DEGs) (Supplementary Fig. 19).

These findings indicate that the transcriptional response elicited by the five-drug ULDC extends beyond the responses observed with individual inhibitors under matched growth-inhibitory conditions.

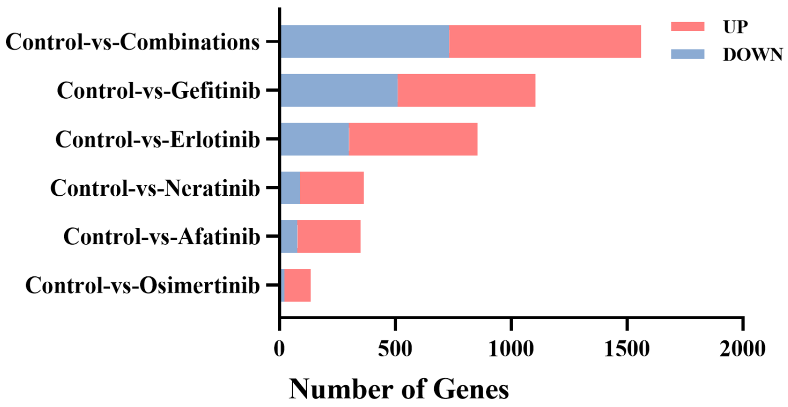


**Supplementary Fig. 19. Numbers of upregulated and downregulated differentially expressed genes induced by individual EGFR-TKIs or the five-drug ULDC.** The ULDC induced broader transcriptional changes than individual inhibitors. Control, PC-9 cells in drug-free complete medium; Combination, five-drug ULDC group.

##### 3.3 ULDC-associated transcriptional responses

To further characterize transcriptional changes associated with the five-drug ULDC, differentially expressed genes identified in the five-drug ULDC group were compared with those identified in individual EGFR-TKI groups. Further comparison across treatment groups identified 470 differentially expressed genes (DEGs) detected in the five-drug ULDC group but not in any of the individual EGFR-TKI DEG sets, whereas only 98 genes were shared across all treatment conditions (Supplementary Fig. 20). The presence of this large set of ULDC-associated transcriptional changes further supports a transcriptional response that is not fully recapitulated by the responses observed with individual EGFR-TKIs.

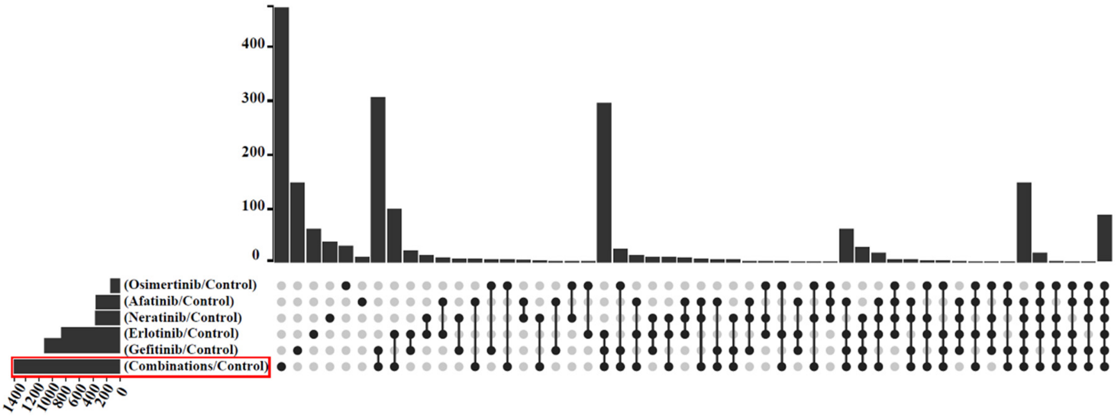

**Supplementary Fig. 20. UpSet plot showing the intersections of differentially expressed genes (DEGs) identified in PC-9 cells treated with individual EGFR-TKIs or the five-drug ULDC compared with control cells.** The left bar chart indicates the number of genes contained in each DEG set, with the y-axis representing the corresponding treatment groups and the x-axis representing the number of DEGs. The upper right bar chart displays the size of each intersection among the DEG sets, with the x-axis representing different DEG intersections and the y-axis representing the number of genes. The lower right matrix indicates the DEG sets contributing to each intersection, with connected filled circles representing the sets included in each intersection. For example, a column containing six connected filled circles represents genes shared among all six treatment conditions. This analysis identified 470 DEGs present in the five-drug ULDC DEG set but not in any of the individual EGFR-TKI DEG sets, and 98 DEGs shared across all six treatment conditions. Control, PC-9 cells in drug-free complete medium; Combination, five-drug ULDC group.

##### 3.4 ULDC-induced transcriptional changes are associated with tumour-relevant biological processes

To investigate the biological significance of genes altered by the five-drug ULDC, KEGG pathway enrichment analysis was performed. ULDC-associated differentially expressed genes were enriched in pathways related to DNA replication, genome maintenance, cancer-associated signalling and other tumour-relevant cellular processes (Supplementary Fig. 21). These enrichment patterns provide molecular context for the broader transcriptional response associated with five-drug ULDC treatment.

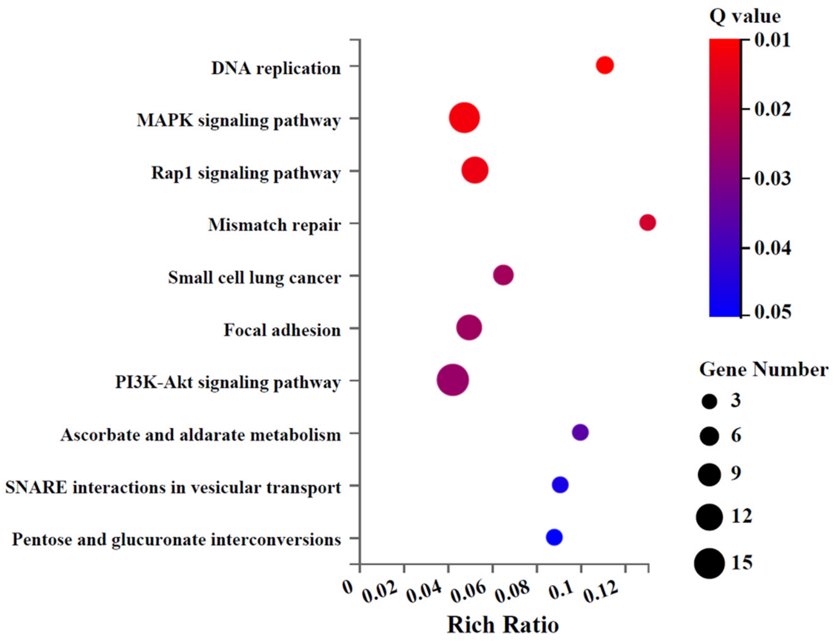


**Supplementary Fig. 21. KEGG pathway enrichment analysis of ULDC-associated differentially**

**expressed genes.** Enriched pathways were related to DNA replication, genome maintenance, cancer-associated signalling pathways, and other tumour-relevant cellular processes.

##### 3.5 Validation of RNA-seq findings by RT-qPCR

Representative differentially expressed genes identified through transcriptomic profiling were further validated by RT–qPCR. The expression trends measured by RT–qPCR were consistent with the RNA-seq results, supporting the reliability of the observed expression changes for the validated genes (Supplementary Fig. 22). Notably, several of the validated genes have previously been implicated in tumour progression or drug resistance. For example, CCND1 has been linked to EGFR-TKI resistance in lung cancer [21], HMGB1 has been associated with treatment resistance in NSCLC and with TKI-resistance-related processes in lung cancer brain metastasis [22,23], and CFL1 has been reported as a prognostic and drug-resistance marker in NSCLC [24]. These genes were downregulated in the five-drug ULDC group relative to the control group, providing additional biological context for the ULDC-associated transcriptional response. Primer sequences used for RT–qPCR validation are provided in Supplementary Table 19.

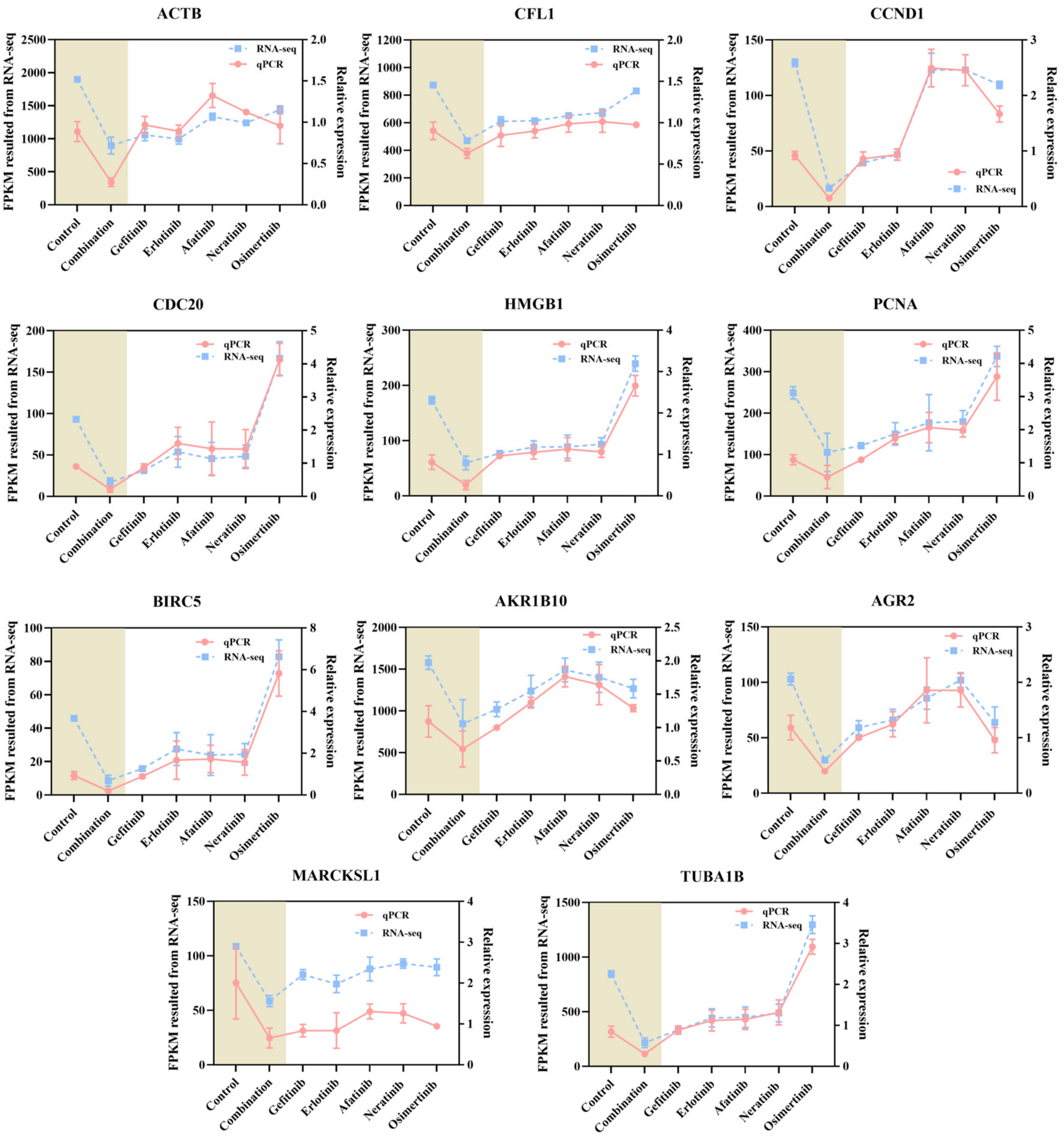


**Supplementary Fig. 22. Validation of RNA-seq findings by RT–qPCR.** Representative genes identified from transcriptomic profiling were selected for validation by RT–qPCR. Expression patterns measured by RT–qPCR were consistent with RNA-seq results. Control, PC-9 cells in drug-free complete medium; Combination, five-drug ULDC group.

**Supplementary Table 19. Primer sequences used for RT–qPCR validation of RNA-seq findings**

| Gene | Forward primer (5′–3′) | Reverse primer (5′–3′) |
|---|---|---|
| CCND1 | AGAGGCGGAGGAGAACAAACAG | GCGGTAGTAGGACAGGAAGTTGTT |
| CDC20 | GACGCCCACCAAGAAGGAAC | CTGATAACCCTCTGGCGCATT |
| HMGB1 | CAAACTTGTCGGGAGGAGCATA | TTGGGCGATACTCAGAGCAGAA |
| PCNA | AGCCATATTGGAGATGCTGTTG | CTGAGTGTCACCGTTGAAGAGAG |
| MARCKSL1 | GTGAACGGAACAGATGAGGCA | TTCTTCTTCTTGGGGGTCTCC |
| AGR2 | CAGTGTCAGCATTCTTGCTCCT | TGTGTGGGCACTCATCCAAGT |
| TUBA1B | GATGAAGTTCGCACTGGCAC | GTTCCATGAGCAGGGAGGTG |
| CFL1 | ATAAGGACTGCCGCTATGCC | ACCTCCTCGTAGCAGTTTGC |
| BIRC5 | CGCATCTCTACATTCAAGAACTG | GGTTTCCTTTGCATGGGGT |
| AKR1B10 | AGGTTCTGATCCGTTTCCATATC | GGTTGCCATCTCCTCATCACTC |
| ACTB | ATGGTAATCGTGCGAGAGGGC | CATCCCAGTTGGTGACAATACC |
| GAPDH | GGAAGCTTGTCATCAATGGAAATC | TGATGACCCTTTTGGCTCCC |

Note: ACTB was included as a target gene for RT–qPCR validation; GAPDH was used as the reference gene for normalization.

### <u>Supplementary Note 4.</u> In vivo antitumour activity and tolerability of the five-drug ULDC

##### 4.1 In vivo collective antitumour activity of the five-drug ULDC

To determine whether the collective growth-inhibitory activity observed in vitro was retained in vivo, we evaluated the five-drug ULDC in a PC-9 xenograft model.

The first five EGFR-TKIs tested in this study, namely gefitinib, erlotinib, afatinib, neratinib, and osimertinib, were administered either individually at ultra-low doses or simultaneously as the five-drug ULDC. The individual doses used in the combination were selected as ultra-low-dose conditions; after conversion to human-equivalent doses using standard Km factors, they corresponded to approximately 5%–10% of the respective WHO-defined daily doses (DDDs), as described in Methods. The gefitinib positive-control group received gefitinib at 25 mg/kg, a conventional dose. Nude mice were treated for 3 weeks, with oral administration for 5 consecutive days followed by 2 drug-free days each week. At these individually ultra-low doses, none of the five EGFR-TKIs significantly inhibited tumour growth relative to the model group. In contrast, the five-drug ULDC markedly inhibited PC-9 xenograft growth throughout the treatment period compared with the model group (Fig. 5a). At the end of treatment, tumour burden in the ULDC group was substantially lower than that in any individual treatment group and was comparable to that observed with gefitinib at 25 mg $kg^{-1}$ (Fig. 5a).

Consistent with the tumour-growth curves, endpoint tumour weights were markedly reduced by the five-drug ULDC. Individual ultra-low-dose EGFR-TKIs produced only modest reductions in tumour weight (approximately 4%–45% inhibition), whereas the five-drug ULDC achieved approximately 90% inhibition of tumour weight, comparable to the approximately 85% inhibition observed with the gefitinib positive-control treatment (Fig. 5b).

Together, these findings demonstrate that multiple individually weak same-target perturbations can generate substantial collective antitumour activity in vivo, extending the collective effects observed in cultured cells to an EGFR-driven xenograft model.

##### 4.2 Reduced EGFR phosphorylation and tumour-cell proliferation in vivo

To assess whether the in vivo antitumour activity of the five-drug ULDC was accompanied by reduced EGFR phosphorylation and tumour-cell proliferation, xenograft tissues were analysed by immunohistochemistry for phosphorylated EGFR (p-EGFR) and the proliferation marker Ki67.

Tumours from the model group showed prominent p-EGFR staining and high Ki67 expression. By contrast, tumours from nude mice treated with the five-drug ULDC showed markedly reduced p-EGFR staining together with substantially lower Ki67 expression (Figs. 5c and 5d).

Collectively, these observations indicate that the collective antitumour activity of the five-drug ULDC was associated with reduced EGFR phosphorylation and tumour-cell proliferation in vivo.

##### 4.3 In vivo tolerability assessment

In vivo tolerability was assessed in the same animal experiment used to evaluate antitumour activity (Section 4.1), by monitoring body weight during treatment and performing histopathological examination of major organs and serum biochemical analyses at the study endpoint.

Body weight increased gradually during treatment in nude mice from both the model and five-drug ULDC groups, with no apparent treatment-associated body-weight loss in the ULDC group (Supplementary Fig. 23). Histopathological examination of the heart, liver, spleen, lung, and kidney showed no overt treatment-associated abnormalities in nude mice receiving the five-drug ULDC or the corresponding individual ultra-low-dose EGFR-TKIs (Supplementary Figs. 24 and 25). In contrast, nude mice treated with gefitinib at 25 mg $kg^{-1}$ showed mild histopathological alterations in the liver and kidney, including focal inflammatory infiltration and mild tubular epithelial swelling.

Serum biochemical analyses were consistent with these histological observations. The five-drug ULDC did not produce significant changes in serum alanine aminotransferase (ALT), aspartate aminotransferase (AST), total protein (TP), or albumin (ALB) relative to the control and model groups. By contrast, the gefitinib positive-control group showed a significant increase in AST (Supplementary Fig. 26).

Similarly, serum creatinine (CREA), uric acid (UA), and urea showed no significant alterations in nude mice treated with the five-drug ULDC, whereas creatinine and uric acid were elevated in the gefitinib positive-control group (Supplementary Fig. 27). No significant treatment-associated differences were observed in lactate dehydrogenase (LDH), creatine kinase (CK), or blood glucose levels (Supplementary Figs. 28 and 29).

Together, these findings indicate that the five-drug ULDC was well tolerated during the treatment period under the conditions tested, with no apparent treatment-associated body-weight loss, overt histopathological abnormalities, or significant alterations in the measured serum biochemical parameters.

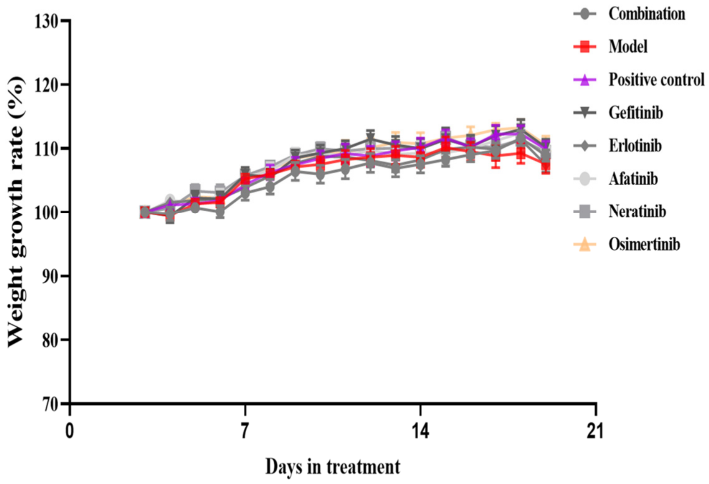


**Supplementary Fig. 23. Body weight expressed as a percentage of baseline during treatment of nude mice in each group.** PC-9 xenograft-bearing nude mice were treated by oral gavage for 3 weeks (5 consecutive treatment days followed by 2 drug-free days per week). Each group comprised ten mice (five females and five males; n = 10). Individual EGFR-TKIs were administered at ultra-low doses that, after conversion to human-equivalent doses using standard Km factors, corresponded to approximately 5%–10% of their respective WHO-defined daily doses (DDDs). Body weight was monitored throughout treatment and expressed as a percentage of baseline. Data are presented as mean ± SD. Model, model group; Combination, five-drug ULDC group; Positive control, gefitinib 25 mg/kg group.

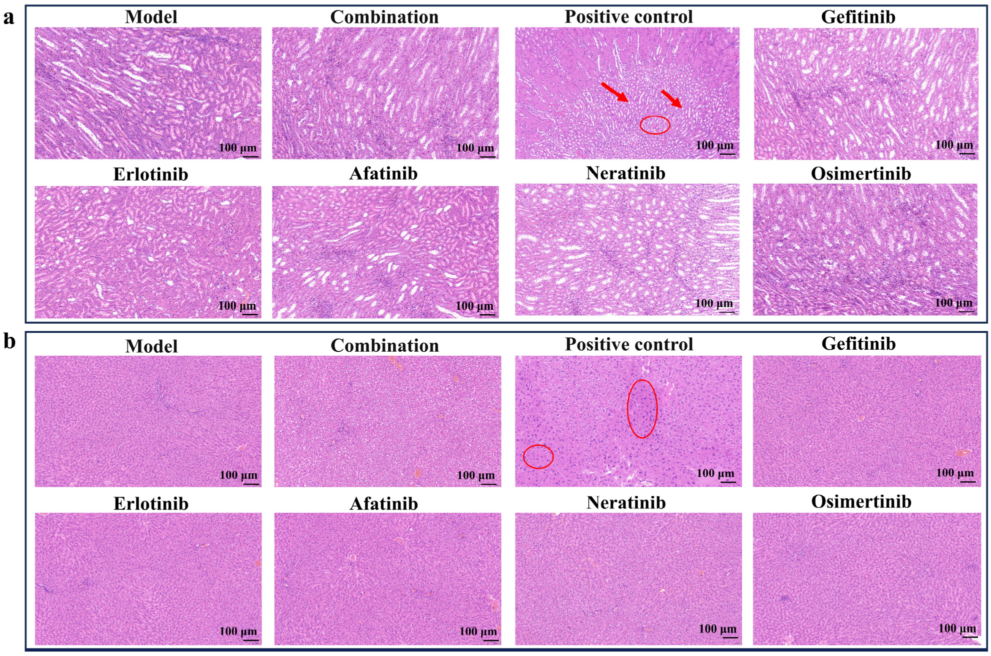

**Supplementary Fig. 24. Representative haematoxylin and eosin (H&E) staining of liver and kidney across experimental groups** (scale bar, 100 μm). **a**, liver; **b**, kidney. Red arrows and/or circles indicate focal inflammatory infiltration in the liver and mild swelling of renal tubular epithelial cells, respectively, in the gefitinib positive-control group. Other annotations, experimental groups, and treatment conditions are described in Supplementary Fig. 23.

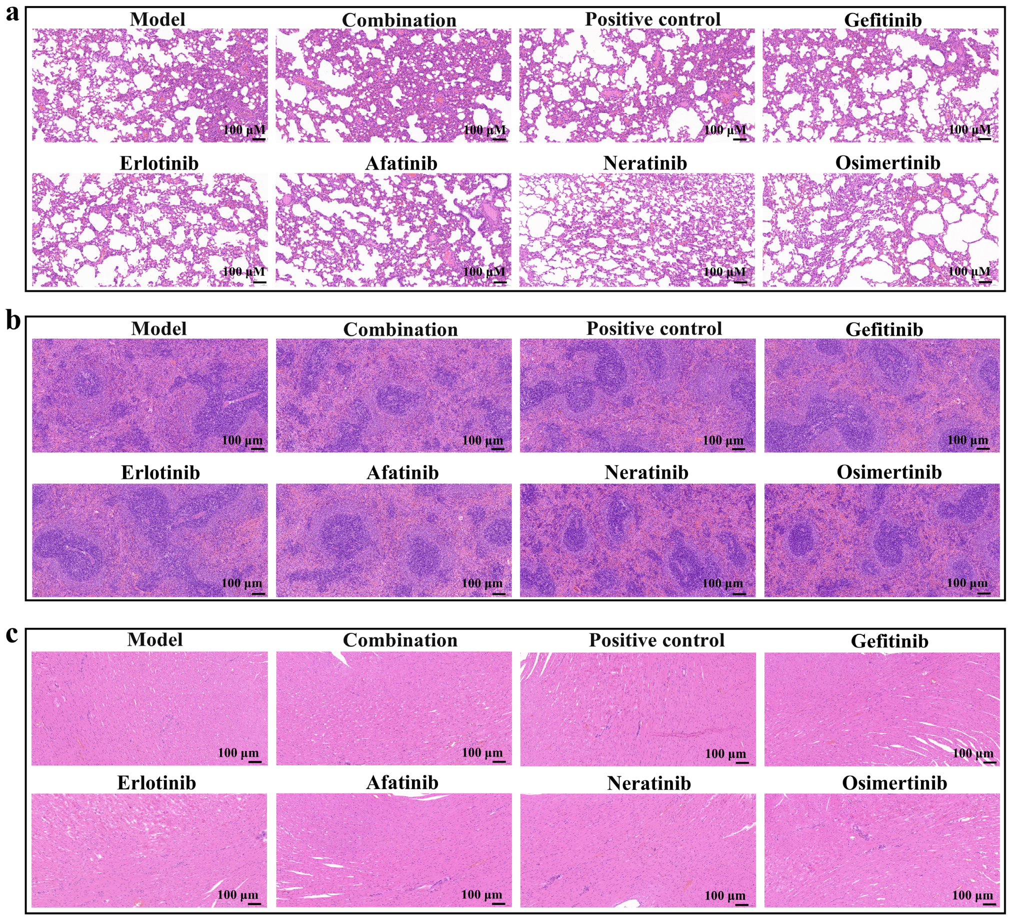


**Supplementary Fig. 25. Representative H&E staining of lung, spleen, and heart across experimental groups** (scale bar, 100 μm). **a**, lung; **b**, spleen; **c**, heart. No apparent treatment-associated pathological changes were observed. Other annotations, experimental groups, and treatment conditions are described in Supplementary Fig. 23.

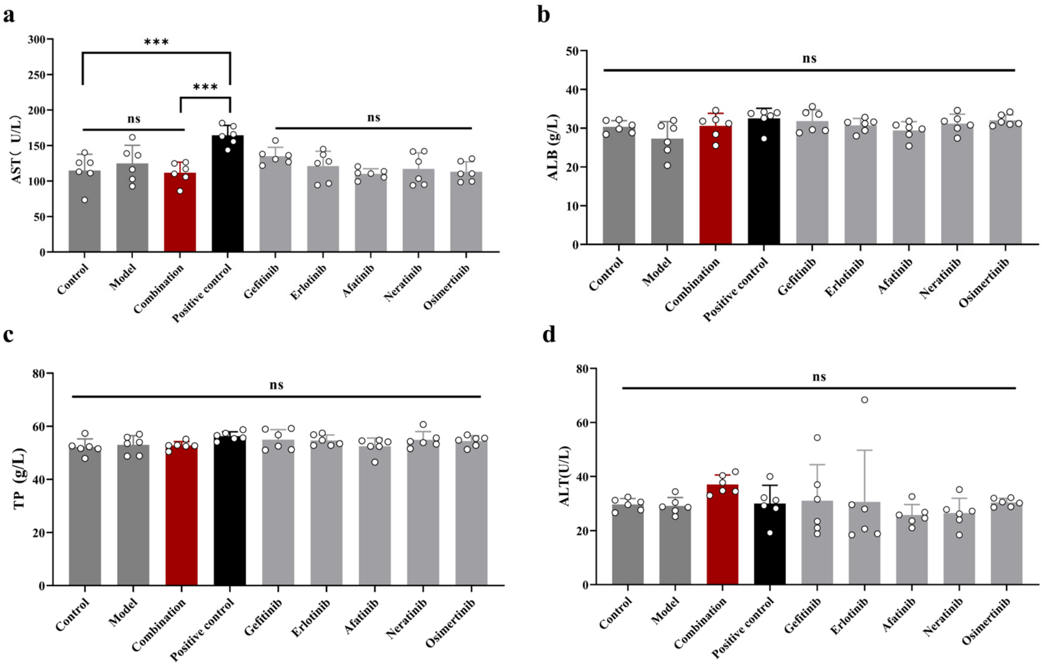


**Supplementary Fig. 26. Assessment of liver function parameters. a**, aspartate aminotransferase (AST); **b**, albumin (ALB); **c**, total protein (TP); **d**, alanine aminotransferase (ALT). Statistical comparisons among groups: ***P < 0.001; ns, not significant. Data are presented as mean ± SD (n = 6). Control, untreated control group with physiological saline. Other annotations, experimental groups, and treatment conditions are described in Supplementary Fig. 23.

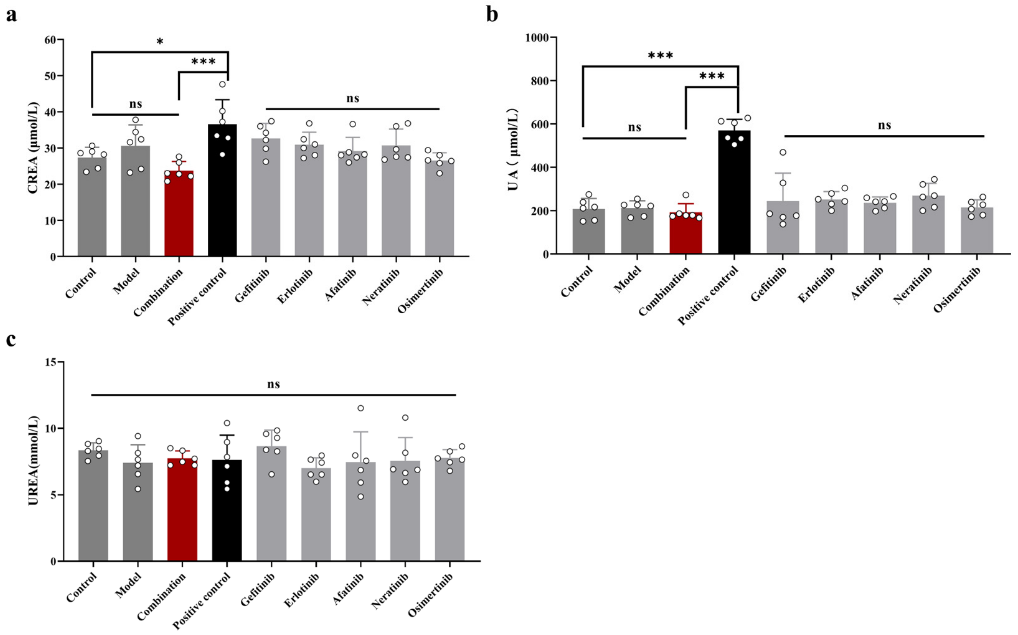


**Supplementary Fig. 27. Assessment of renal function parameters**. **a**, creatinine (CREA); **b**, uric acid (UA); **c**, urea (UREA). Statistical comparisons among groups: ***P < 0.001; *P < 0.05; ns, not significant. Data are presented as mean ± SD (n = 6). Other annotations, experimental groups, and treatment conditions are described in Supplementary Fig. 26.

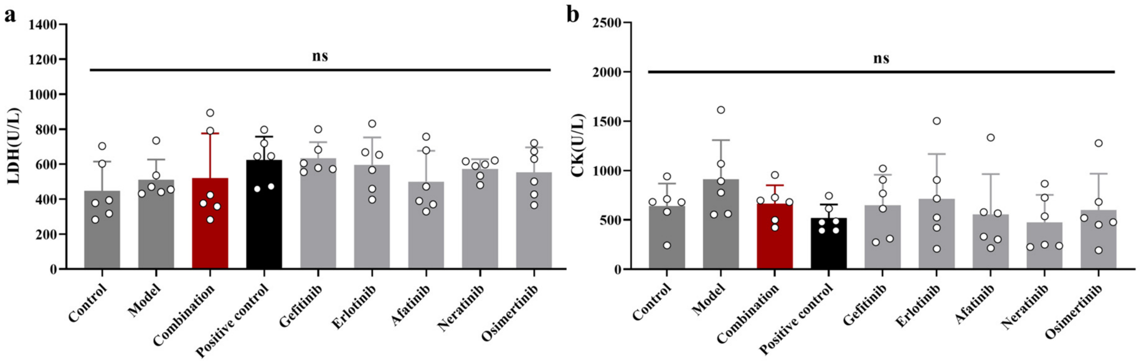


**Supplementary Fig. 28. Assessment of serum lactate dehydrogenase (LDH) and creatine kinase (CK) levels. a**, lactate dehydrogenase (LDH); **b**, creatine kinase (CK). Statistical comparisons among groups: ns, not significant. Data are presented as mean ± SD (n = 6). Other annotations, experimental groups, and treatment conditions are described in Supplementary Fig. 26.

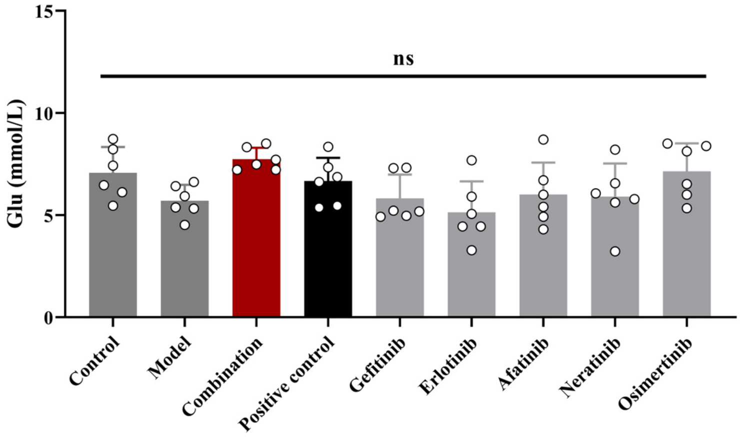


**Supplementary Fig. 29. Assessment of blood glucose levels.** Statistical comparisons among groups: ns, not significant. Data are presented as mean ± SD (n = 6). Glu, glucose. Other annotations, experimental groups, and treatment conditions are described in Supplementary Fig. 26.

#### Supplementary Discussion. Conceptual origin of the same-target combination strategy

The ULDC strategy originated from our earlier work on multicomponent pharmacology. In studying the pharmacological basis of traditional Chinese medicines, we considered the possibility that multiple bioactive compounds, including chemical constituents and their metabolites in vivo, each present at a relatively low concentration, could collectively generate substantial biological activity when acting on the same molecular target.

This led us to propose that individually weak perturbations directed against the same target could collectively produce a biologically meaningful response, a phenomenon described in our earlier work as “additive effects” [42,43,44,45]. In the present study, we refer to this broader concept as “same-target collective effects”. Previous studies from our group using defined bioactive compounds derived from traditional Chinese medicines provided experimental support for same-target collective effects, including combinations of multiple compounds acting on a shared molecular target [46,47].

We subsequently sought to examine this principle in more experimentally controlled pharmacological systems using chemically defined drugs with established molecular targets. This allowed the number, identity, and dose of individual components to be systematically controlled while preserving the central feature of the original concept: multiple individually weak pharmacological perturbations directed against the same target. This concept was subsequently extended to chemically defined anticancer combinations, including an ultra-low-dose combination of eight DNA topoisomerase inhibitors. In these studies, multiple individually weak drug perturbations generated substantial collective antitumour activity while markedly reducing treatment-associated toxicity, an effect termed the “toxicity-scattering effect” in our previous work [18,48,49]. These observations suggested that the architecture of a pharmacological perturbation—not only the potency or dose of an individual drug—might influence the biological properties of a treatment.

The present study extends this concept beyond antitumour activity and toxicity to the evolution of drug resistance. We asked whether distributing individually weak perturbations across multiple inhibitors of the same oncogenic target could alter long-term resistance outcomes compared with a single inhibitor at a concentration producing a similar level of growth inhibition. EGFR-mutant lung cancer provided a tractable system in which to test this question because multiple structurally diverse EGFR inhibitors with differing pharmacological properties are available against the same clinically validated oncogenic target. This design further allowed systematic variation of same-target inhibitor diversity under approximately matched levels of growth inhibition while maintaining low component doses. The resulting experiments therefore tested not simply whether multiple EGFR inhibitors could generate substantial collective growth-inhibitory activity, but whether increasing same-target inhibitor diversity could influence resistance emergence and whether resistant states arising under five-drug ULDC treatment differed from those generated under prolonged single-agent treatment. Portions of the experimental work underlying the present study were previously included in Z.-G. Zhao's doctoral dissertation [50].

The findings reported here support this extension of the same-target combination concept. Individually weak EGFR-TKI perturbations produced substantial collective growth-inhibitory activity when combined, while no detectable resistance emerged in the higher-diversity same-target ULDC groups within the observation window. The five-drug ULDC also retained substantial growth-inhibitory activity against gefitinib-resistant cells, and resistant cells arising during prolonged five-drug ULDC treatment showed greater reversibility after drug withdrawal than resistant cells generated under prolonged single-agent treatment. Together, these observations led us to formulate the broader framework developed in this study: same-target collective suppression concentrates multiple individually weak perturbations on a shared oncogenic target, while increasing same-target inhibitor diversity is proposed to impose progressively greater adaptive constraint by restricting the range of trajectories compatible with continued target function and tumour growth.

Importantly, the present experiments do not directly evaluate the efficacy, toxicity, or resistance properties of traditional medicines. Rather, earlier studies of multicomponent pharmacology provided the conceptual starting point for asking whether individually weak perturbations directed against a shared target could collectively generate substantial biological effects. By translating this conceptual question into chemically defined and experimentally controlled drug systems, the present work develops that initial idea into a testable framework for investigating how the architecture of therapeutic perturbation may influence both antitumour activity and resistance evolution.

#### REFERENCES


**1**. Li, J., Hu, J., Yang, Y. *et al.* Drug resistance in cancer: molecular mechanisms and emerging treatment strategies. *Mol. Biomed*. **6**, 111 (2025).

2. Dagogo-Jack, I. & Shaw, A. T. Tumour heterogeneity and resistance to cancer therapies. *Nat. Rev. Clin. Oncol.* **15** (2), 81–94 (2018).

3. Marine, J.-C., Dawson, S.-J., & Dawson, M. A. Non-genetic mechanisms of therapeutic resistance in cancer. *Nat. Rev. Cancer* **20** (12), 743–756 (2020).

4. Jin, H., Wang, L., & Bernards, R. Rational combinations of targeted cancer therapies: background, advances and challenges. *Nat. Rev. Drug Discov.* **22** (3), 213–234 (2023).

**5**. Obenauf, A. C. Mechanism-based combination therapies for metastatic cancer. *Sci. Transl. Med.* **14** (655), eadd0887 (2022).

**18**. Wang, Y., Xu, F., Zhao, Q. *et al.* Combining same-target drugs exhibits additive efficacy with minimal toxicity [preprint]. *Research Square* (2024)**:** https://doi.org/10.21203/rs.3.rs-4822017/v1

**19**. Chou, T. C. Drug combination studies and their synergy quantification using the Chou-Talalay method. *Cancer Res.* 70 (2), 440–446 (2010).

**20**. Chou, T. C. Theoretical basis, experimental design, and computerized simulation of synergism and antagonism in drug combination studies. *Pharmacol. Rev*. 58 (3), 621–681 (2006).

**42**. Xu, F., Yang, D.-H., Shang, M.-Y., Wang, X., & Cai, S.-Q. Effective forms, additive effect, and toxicity-scattering effect of pharmacodynamic substances of TCMs: some reflections evoked by the study on the metabolic disposition of traditional Chinese medicines (TCM). *World Sci. Technol.-Mod. Tradit. Chin. Med.* 12, 688–703 (2014).

**43**. Cai, S.-Q., Wang, X., Shang, M.-Y., Xu, F., & Liu, G. "Efficacy theory" may help to explain characteristic advantages of traditional Chinese medicines. *Zhongguo Zhongyao Zazhi = China J. Chin. Mater. Medica* 40, 3435–3443 (2015).

**44**. Cai, S.-Q., Feng, X., Shang, M.-Y., & Wang, X. Design idea of ultra-hypotoxic multidrug based on the efficacy theory of traditional Chinese medicines. *Planta Medica* **82** (S 01), S1–S381 (2016).

**45**. Cai, S.-Q., Xu, F., Wang, Y. *et al.* A combination drug with both additive effect on the efficacy and toxicity-scattering effect. *China patent announcement* 2016, CN105288629A,1–26 (2016); Patent application No. 201410234283.6 (2014); Patent No. ZL 2014 1 0234283.6 (2021).

**46**. Liu, L.-J. Effective forms of bioactive substances from Astragalus flavonoids and their additive effects. *Doctoral dissertation, Peking University*, May 2019.

**47**. Han, X. Additive effects of flavonoids from Astragali Radix on α-glucosidase inhibition and their antihyperglycemic activity. *Doctoral dissertation, Peking University*, October 2024.

**48**. Wang, Y. Additive effect and toxicity-scattering effect of eight antitumour drugs acting on the same target. *Doctoral dissertation, Peking University*, May 2017.

**49**. Cai, S.-Q., Xu, F., Wang, Y. *et al.* A pharmaceutical combination used for preventing and/or treating tumors. *China patent announcement* 2025, CN 120983458 A; Patent application No. 202511167867.0 (2025).

**50**. Zhao, Z.-G. Low-dose EGFR-TKI combinations for overcoming drug resistance in NSCLC and their co-delivery using exosome-based nanosystems. *Doctoral dissertation, Southwest Jiaotong University*, May 2025.
